# Molecular diversity maintained by long-term balancing selection in mating loci defines multiple mating types in fungi

**DOI:** 10.1101/2021.09.10.459787

**Authors:** David Peris, Dabao Sun Lu, Vilde Bruhn Kinneberg, Ine-Susanne Methlie, Malin Stapnes Dahl, Timothy Y. James, Håvard Kauserud, Inger Skrede

**Affiliations:** Section for Genetics and Evolutionary Biology, Department of Biosciences, University of Oslo, N-0316 Oslo, Norway; Department of Health, Valencian International University (VIU), 46002 Valencia, Spain; Department of Ecology and Evolutionary Biology, University of Michigan, Ann Arbor, Michigan 48109, USA

**Keywords:** Balancing selection, mating loci, comparative genomics, fungi, *Trichaptum*

## Abstract

Balancing selection, an evolutionary force that retains genetic diversity, has been detected in multiple genes and organisms, such as the sexual mating loci in fungi. However, to quantify the strength of balancing selection and define the mating-related genes require a large number of specimens. In tetrapolar basidiomycete fungi, sexual type is determined by two unlinked loci, *MATA* and *MATB*. Genes in both loci defines mating type identity, control successful mating and completion of the life cycle. These loci are usually highly diverse. Previous studies have speculated, based on culture crosses, that species of the non-model genus *Trichaptum* (Hymenochaetales, Basidiomycota) possess a tetrapolar mating system, with multiple alleles. Here, we sequenced a hundred and eighty specimens of three *Trichaptum* species. We characterized the chromosomal location of *MATA* and *MATB*, the molecular structure of *MAT* regions and their allelic richness. Our sequencing effort was sufficient to molecularly characterize multiple *MAT* alleles segregating before the speciation event of *Trichaptum* species. Our analyses suggested that long-term balancing selection has generated trans-species polymorphisms. Mating sequences were classified in different allelic classes based on an amino acid identity (AAI) threshold supported by phylogenetics. The inferred allelic information mirrored the outcome of *in vitro* crosses, thus allowing us to support the degree of allelic divergence needed for successful mating. Even with the high amount of divergence, key amino acids in functional domains are conserved. The observed allelic classes could potentially generate 14,560 different mating types. We conclude that the genetic diversity of mating in *Trichaptum* loci is due to long-term balancing selection, with limited recombination and duplication activity. Our large number of sequenced specimens highlighted the importance of sequencing multiple individuals from different species to detect the mating-related genes, the mechanisms generating diversity and the evolutionary forces maintaining them.

**Author summary:** Fungi have complex mating systems, and basidiomycete fungi can encode thousands of mating types. Individuals with the same mating type cannot mate. This sexual system has evolved to facilitate sexual mating, increasing the chances to recombine into advantageous allelic combination and prune deleterious alleles. We explored the genomes of hundred and eighty specimens, combined with experimental mating studies of selected specimens, from a non-model organism (*Trichaptum*). We characterized the genomic regions controlling sex. The mating ability of the specimens confirmed the role of the mating alleles observed in the genomic data. The detailed analyses of many specimens allowed us to observe gene duplication and rearrangements within the mating loci, increasing the diversity within these loci. We supported previous suggestions of balancing selection in this region, an evolutionary force that maintains genomic diversity. These results supports that our fungal specimens are prone to outcross, which might facilitate the adaptation to new conditions.

## Introduction

Balancing selection is an evolutionary force that maintains genetic diversity [1]. Due to the importance of balancing selection to generate diversity, it has received long-term attention in evolutionary biology [2]. Heterozygote advantage [1], pleiotropy [3], negative frequency-dependent selection [4], rapid temporal fluctuations in climate [5], and segregation distortion balanced by negative selection [6, 7] are modes of balancing selection. These different modes of balancing selection leave similar genomic signatures, such as an increased number of polymorphic sites around the region under balancing selection, and sometimes an enrichment of intermediate-frequency alleles around the selected genomic region [1]. When balancing selection has persisted for a long period, coalescent time of alleles may predate speciation events, and polymorphisms can become shared among distinct species, leading to trans-species polymorphisms [8]. Phylogenetic trees for balanced regions are characterized by the presence of long internal branches [9], and clades with a mixture of species caused by trans-species polymorphisms [10]. The development of methods to detect the genomic footprints of balancing selection [11–13] has unraveled, also with a low number of individuals due to sequencing costs, multiple loci under this type of selection. Well-known examples include: the major histocompatibility locus (MHC) in vertebrates [8]; the ABO histo-blood [14]; non-MHC genes, such as *TRIM5* and *ZC3HAV1* in humans [15, 16]; self-incompatibility (SI) loci in plants [17, 18] and self/nonself-recognition during vegetative growth in fungi [19]; multilocus metabolic gene networks, such as the *GAL* network in *Saccharomyces* [20, 21]; and sexual mating loci in fungi [22].

In basidiomycete fungi, there are numerous examples of balancing selection acting on loci regulating the sexual cycle [22–26]. In this phylum, the sexual cycle involves fusion (plasmogamy) of two genetically distinct monokaryotic hyphae (n or one set of chromosomes), generating a dikaryotic (n+n) hyphae [27–29]. The dikaryon is considered a more stable and long-lived state than the monokaryotic phase, but there are controversies about this assumption due to limited studies [30, 31]. Due to this dikaryotic state, plasmogamy is normally separated in time from karyogamy, the fusion of both parental nuclei [32]. In basidiomycetes, karyogamy and meiosis normally occur in specialized structures, the fruit bodies [32]. Mating between two monokaryotic hyphae is determined by one or two sets of multiple allelomorphic genes in the mating (*MAT*) loci. Two different mating systems have evolved among basidiomycetes, referred to as bipolar or tetrapolar mating systems [33]. Mating-type identity in some basidiomycetes, such as *Cryptococcus neoformans,* and members of the sister phylum Ascomycota i.e. *Saccharomyces cerevisiae*, is governed by a single *MAT* locus [34]. This case corresponds to the bipolar system, resembling the sexual system (male or female) in metazoans [35]. However, the ancestor of basidiomycetes developed an evolutionary innovation, the tetrapolar mating system, where two *MAT* loci regulate mating [36]. This new system hinders inbreeding more effectively, since only 25% of the spores from the same individual can mate, compared to 50% for the bipolar species [37]. At the same time, having multiple mating alleles in each *MAT* locus enables extremely effective outcrossing, where most monokaryotic spores or mycelia (derived from different individuals) can establish a dikaryotic mycelium when a compatible partner is found [38].

In strict tetrapolar organisms, the *MATA* locus (syn. *b* or *HD*) contains a series of linked pairs of homeodomain-type transcription factor genes (*HD1-HD2*, syn. *bW-bE*), whereas the *MATB* locus (syn. *a* or *P/R*) is composed of tightly linked G-pheromone receptors (*STE3*, syn. *Rcb, pra*) and pheromones (*Phe3,* syn. *Ph, mfa*) [23,39–46]. These genes define mating type identity [34], which controls successful mating and completion of the life cycle [32]. Nucleotide differences in mating-related genes, without sufficient amino acid changes in key functional domains, belong to the same allelic class [22]. These allelic classes define the mating type in *MATA* and *MATB*. When monokaryotic (haploid) hyphae of compatible allelic classes, different *MATA* and *MATB* types, conjugate, a structure involved in transferring one of the nuclei during cell division can be observed, called clamp connection, indicating a successful mating [47]. Proteins encoded by *MATA* genes initiate the pairing of the two parental nuclei within dikaryons, they promote clamp development, synchronize nuclear division and septum formation. Proteins encoded by *MATB* genes coordinate the completion of clamp fusion with the subapical cell after synchronized nuclear division and the release of the nucleus, which was initially trapped within the unfused clamp cell [48, 49]. Once monokaryons have fused, the *MATB* proteins facilitate septum dissolution and nuclear migration [39]. Experimental crossings in various basidiomycetes, such as *Coprinopsis* and *Schizophyllum*, have been used to infer the number of *MATA* and *MATB* alleles, and results suggest that 12,800-57,600 mating types may exist in some species [50].

However, the molecular confirmation and the knowledge of the diversity of such genomic regions are far behind, as multiple specimens must be sequenced. One of the reasons to this delay, is the high nucleotide divergence among *MAT* alleles, which has complicated the study of molecular evolution of the fungal mating systems, where e.g. primer design has been a challenge. Moreover, until now, only a limited number of specimens from different species have been analyzed, mainly due to sequencing costs, limiting the quantification of the strength of balancing selection, the presence of trans-species polymorphisms and the detection of mating and non-mating related genes. Due to limited availability of sequenced specimens, how each genes within mating loci are involved in mating is unknown.

Speculations about the mating system in two non-model *Trichaptum* sister species, *Trichaptum abietinum* and *Trichaptum fuscoviolaceum* (Hymenochaetales, Basidiomycota), have been done in the past, likely because their fruit bodies readily produce monokaryotic spores that germinates and grows *in vitro*, making it easy to conduct crossing experiments in the lab [51]. *Trichaptum abietinum* and *T. fuscoviolaceum* are wood-decay fungi with circumboreal distributions [52]. Although, we know their life cycle (Figure 1A), details about how long these organisms spend in monokaryotic or dikaryotic states are still unknown. Previous mating studies have suggested a tetrapolar mating system for *Trichaptum* with an inferred number of 385 *MATA* and 140 *MATB* alleles in *T. abietinum* [53]. The mating studies have also revealed that three intersterility groups (ISGs) occur in *T. abietinum* [50–54]. However, so far we have no information about the underlying genomic architecture and molecular divergence of *Trichaptum* mating genes.

**Figure 1.**
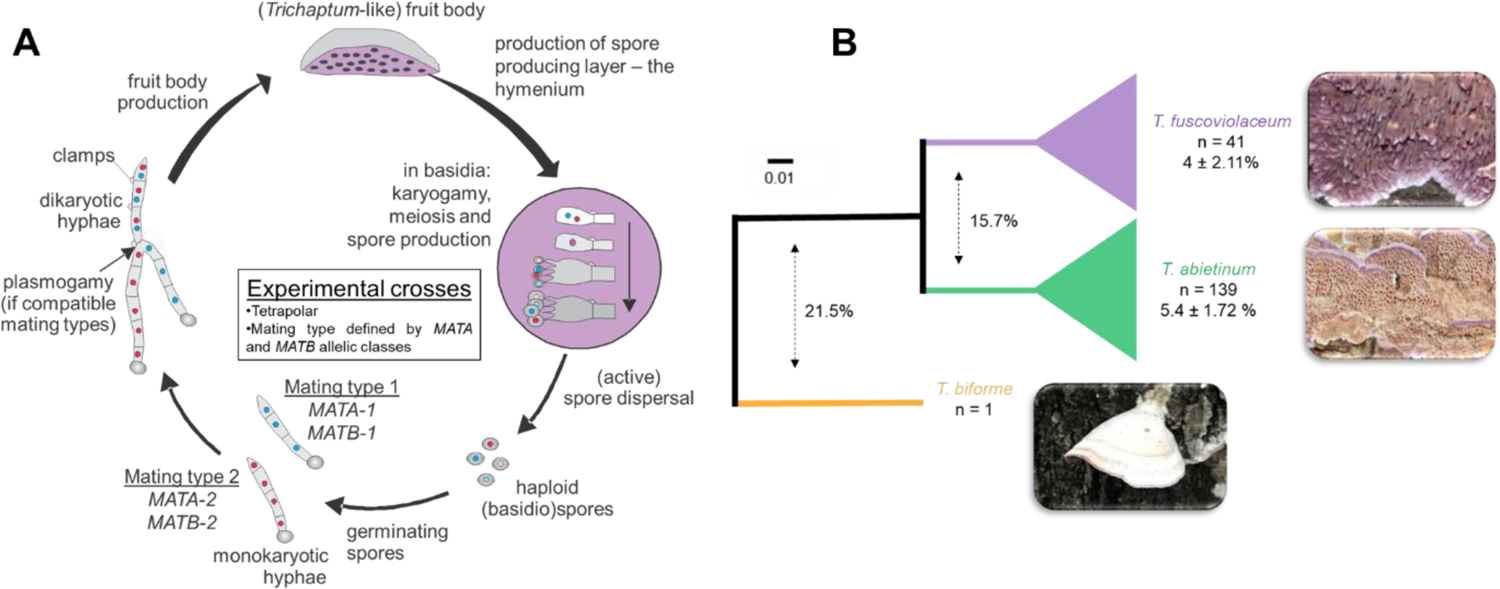
Trichaptum abietinum and T. fuscoviolaceum are sister-species. A) Schematic representation of the *Trichaptum* life cycle. As an example, allelic classes, for *MATA* and *MATB,* generating two compatible mating types are indicated. B) Schematic Neighbor-Joining (NJ) phylogenetic tree reconstructed using 100 - ANI values. More detailed (uncollapsed species clades) NJ and ASTRAL phylogenetic trees can be found in Supplementary Figure 1 and in iTOL: https://itol.embl.de/shared/Peris_D. The number of specimens (n) and the average 100 - ANI within species are indicated for each species clade. The L15831 genome is included increasing the *T. abietinum* collection to 139 specimens. Dashed arrows indicate the average 100 - ANI of pairwise specimen comparisons for the compared species. Colors highlight the species designation after the whole genome sequencing analysis.

Here, we study the molecular evolution of the *MAT* genes in tetrapolar basidiomycetes, using a non-model organism. By combining, full genome sequencing of a large set of new established monokaryotic cultures from sporulating fruit bodies, collected at different circumboreal locations, bioinformatics and *in vitro* crosses, we want to: i) unravel the genomic location and the structure of the mating-related genes; ii) assess the allelic richness of *MAT* genes; iii) the divergence needed among the alleles in order for the fungi to recognize different mating types, then test whether the genotypic information mirrors phenotypic outcomes of *in vitro* sexual mating; iv) and reveal molecular signals of balancing selection.

## Results

### Mating regions are highly dynamic in Trichaptum species

To locate the chromosomal position of *MATA* and *MATB* and the genes delimiting the mating regions, we generated PacBio assembly genomes (Table 1) for one *T. abietinum* and one *T. fuscoviolaceum* specimen. These two species genomes differed with an average 15.7% in a converted ANI (average nucleotide identity) value to divergence value (Figure 1B).

**Table 1.** PacBio assembly stats.

|  |  |  |  | Before ultrascaffolding |  |  | After correction & ultrascaffolding |  |  |
| --- | --- | --- | --- | --- | --- | --- | --- | --- | --- |
| Specimen name | Descended from | Species | Assembler | Contigs | N50 (Kb) | L50 | Scaffolds | N50 (Kb) | Bases (Mb) |
| TA10106M1 | TA-1010-6 | *T. abietinum* | Canu | 26 | 4,268.52 | 5 | 12 | 4,354.20 | 49.43 |
| TF100210M3 | TF-1002-10 | *T. fuscoviolaceum* | Canu | 118 | 2,011.66 | 10 | 12 | 5,547.79 | 59.09 |

Both species potentially contained twelve chromosomes. The genome size of *T. abietinum* and *T. fuscoviolaceum* was 49 Mbp and 59 Mbp, respectively. Both genomes were highly syntenic with a few small inversions (Supplementary Figure 2). The *MATA* and *MATB* loci were located on chromosomes 2 and 9, respectively. *MATA* homeodomains genes were flanked by *bfg, GLGEN* on one end and *MIP1* coding sequences on the other (Figure 2). The *MATA* region, defined from *bfg* to *MIP1*, was 17.9 and 19.6 Kbp long in *T. abietinum* and *T. fuscoviolaceum,* respectively. Both reference genomes contained two homeodomain complexes: alpha- (*aHD*) and beta-complexes (*bHD*). In the reference *T. fuscoviolaceum MATA* region, one homeodomain pair, the *bHD1*, was lost, *bHD2* was inverted, and between the alpha and beta-complexes there was a gene encoding an *ARM*-repeat containing protein (Figure 2). *MATB* pheromone receptors and pheromones were flanked by *PAK, RSM19, DML1, RIC1* and *SNF2* genes. All these genes together were defined as the *MATB* region, which was 30.3 Kbp long in both species. Four putative pheromone receptors and two pheromone genes were annotated. The *MATB* region was syntenic between both species, except an inverted block containing *STE3.2* and *Phe3.2* genes in the *T. fuscoviolaceum* reference (Figure 2).

**Figure 2.**
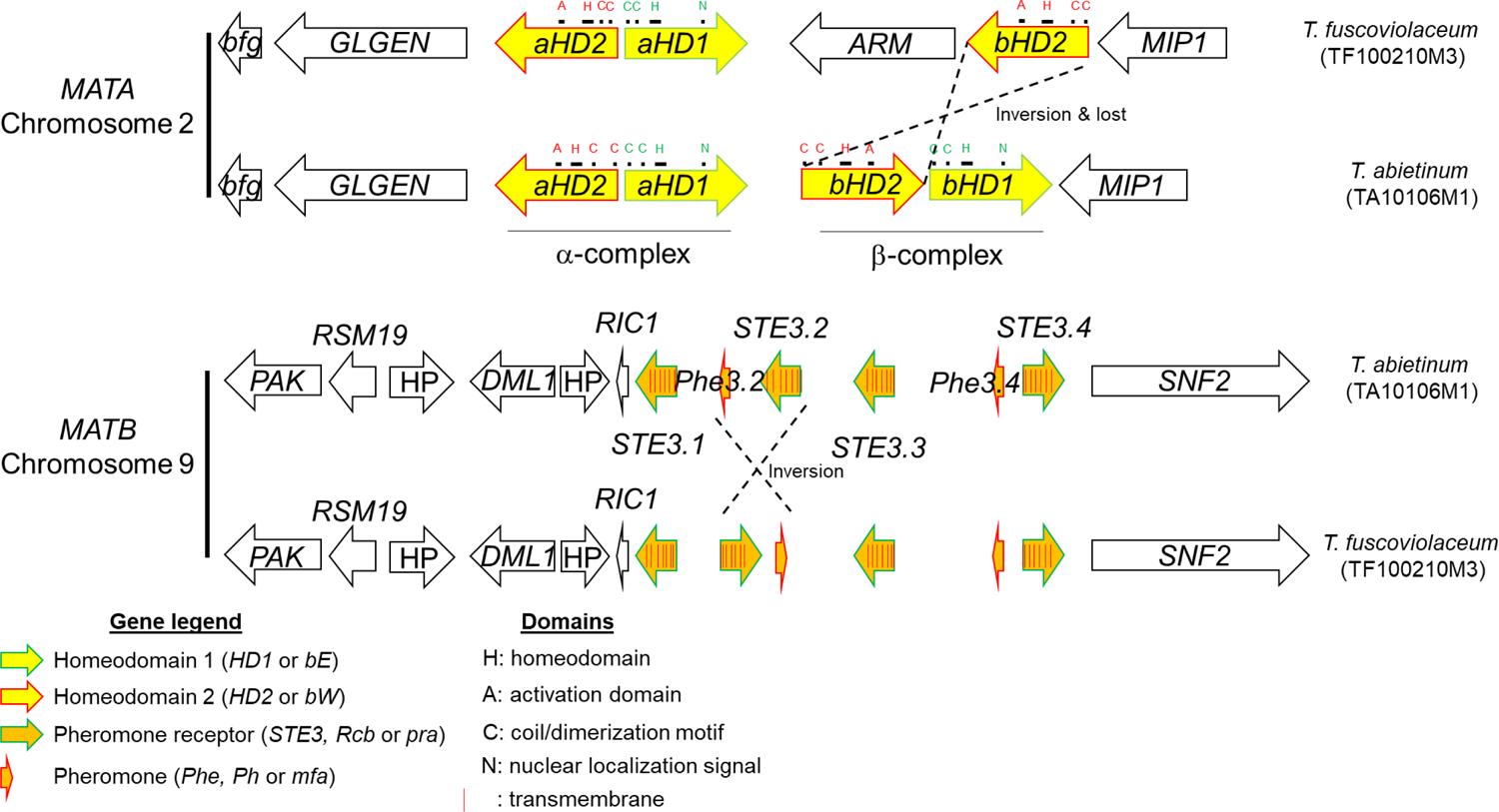
Two homeodomain complexes in *MATA* and four putative pheromone receptors in *MATB* were detected in *T. abietinum* and ***T. fuscoviolaceum*.** Schematic representation of the gene composition and direction in both PacBio reference genomes. Homeodomain, pheromone and pheromone receptors genes are represented as indicated in the legend. The rest of the genes were colored in black, and the gene names were indicated inside the arrows. *aHD*: alpha-complex homeodomain*; ARM:* ARM-repeated containing protein; *bfg* beta-flanking gene; *bHD:* beta-complex homeodomain; *DML1*: mtDNA inheritance protein; *GLGEN:* glycogenin-1; HP: hypothetical protein; *MIP1:* mtDNA intermediate peptidase; *PAK:* serine/threonine protein kinase; *RSM19:* 37S ribosomal protein S19*; RIC1:* RIC1-domain containing protein*; SNF2:* Snf2 family dna-dependent ATPase; *STE3:* GPCR fungal pheromone mating factor.

### MAT genes displayed multiple alleles

The annotated mating genes in the reference genomes were used to search for those genes in the 178 Illumina sequenced specimens, collected at circumboreal regions (Figure 3) and a *T. abietinum* assembly downloaded from JGI (Supplementary Table 1). *Trichaptum abietinum* was the most diverse species (average converted ANI 5.4%) (Figure 1B). *MATA* genes were assembled in one contig for 75 *T. abietinum,* 25 *T. fuscoviolaceum* and 1 *T. biforme*. In the case of *MATB* genes, genes in that region were found in one contig for 116 *T. abietinum,* 27 *T. fuscoviolaceum* and 1 *T. biforme*. For these specimens, the mating genes have potentially the same chromosomal location than in our reference specimens. For the rest of the sequenced specimens, the mating genes were found in multiple contigs due to assembly limitations using short reads. Most of those fragmented mating regions might be organized as in our reference specimens; however, we observed unexpected coding sequences for 6 specimens in the *MATA* region and 2 specimens in the *MATB* region, which could suggest that these regions have split and were translocated to different chromosomes or positioned in a new chromosomal location (Supplementary Table 2).

**Figure 3.**
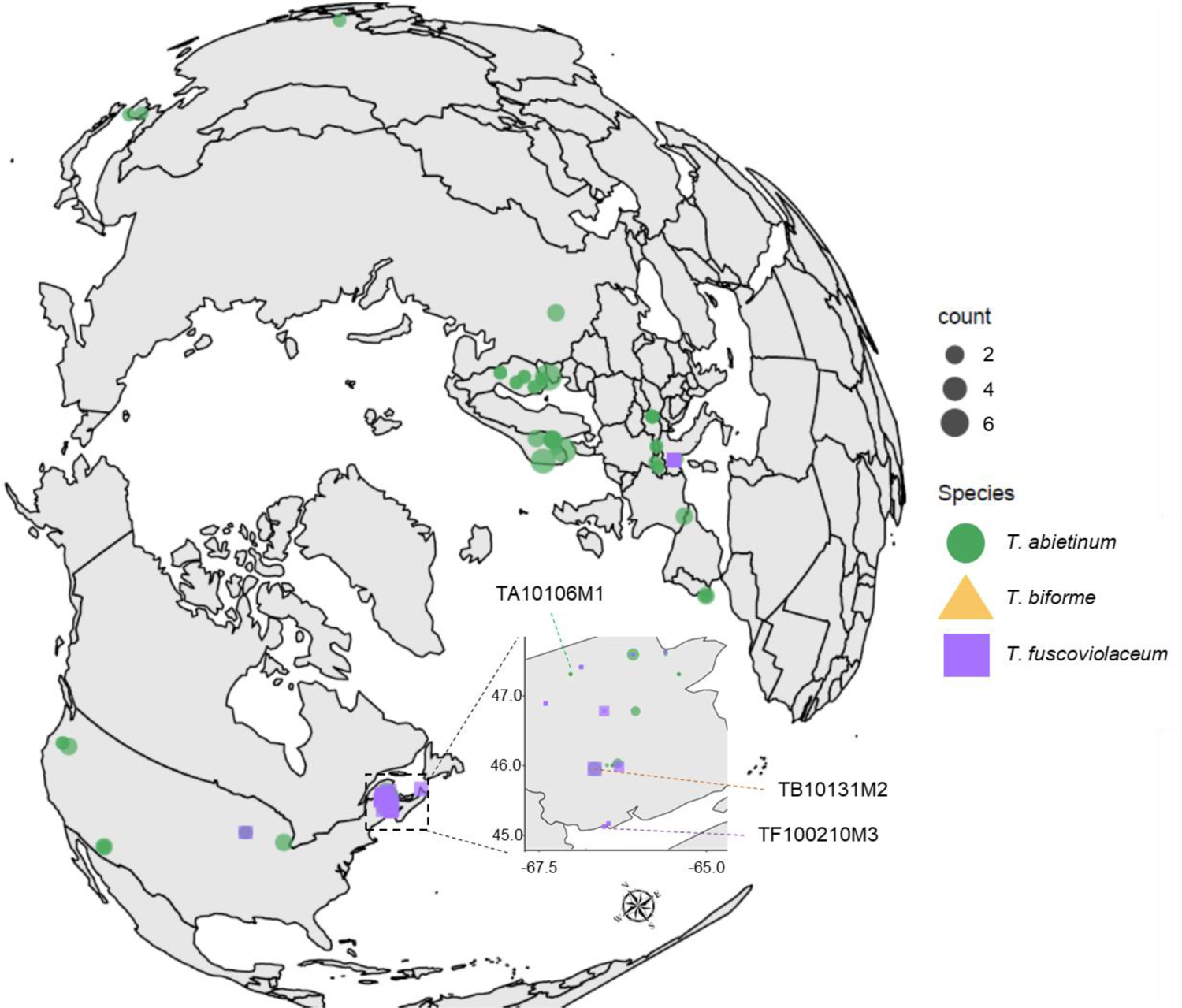
Circumboreal distribution of *Trichaptum* specimens. Geographic distribution of our *Trichaptum* specimens.

An initial analysis of nucleotide conservation of the mating regions indicated that flanking genes were conserved, as well as *STE3.1* and *STE3.3.* However, the rest of putative mating genes were highly diverse (Figure 4). Gene order comparison among specimens highlighted that the most common *MATA* and *MATB* syntenic blocks were both present in *T. abietinum* and *T. fuscoviolaceum*, and the frequent *MATB* syntenic block was present in the three species (Figure 5). *Trichaptum biforme* and five other *Trichaptum* specimens, differentiated from the most frequent *MATA* configuration by the presence of a hypothetical protein (Figure 5A). All this suggest that the most frequent *MATA* and *MATB* gene configurations, represented in Figure 2 for *T. abietinum,* were present in the ancestor of these three *Trichaptum* species. The gene order of *HDs* in the alpha complex was conserved among all *Trichaptum* specimens. However, frequent inversions of the *bHD2* gene and absence of one of the two *bHD* genes were detected. An interesting observation was the presence of an additional *HD2* gene (*xHD2*) upstream the alpha complex in six *T. abietinum* specimens (Figure 5A). The coding sequence of *xHD2* looks truncated, indicating an ongoing process of pseudogenization. In the *MATB* region, all specimens contained two pheromones, one located between *STE3.1* and *STE3.2*, and a second between *STE3.3* and *STE3.4.* The orientation of *STE3.2, STE3.4* and pheromone genes varied among specimens (Figure 5B).

**Figure 4.**
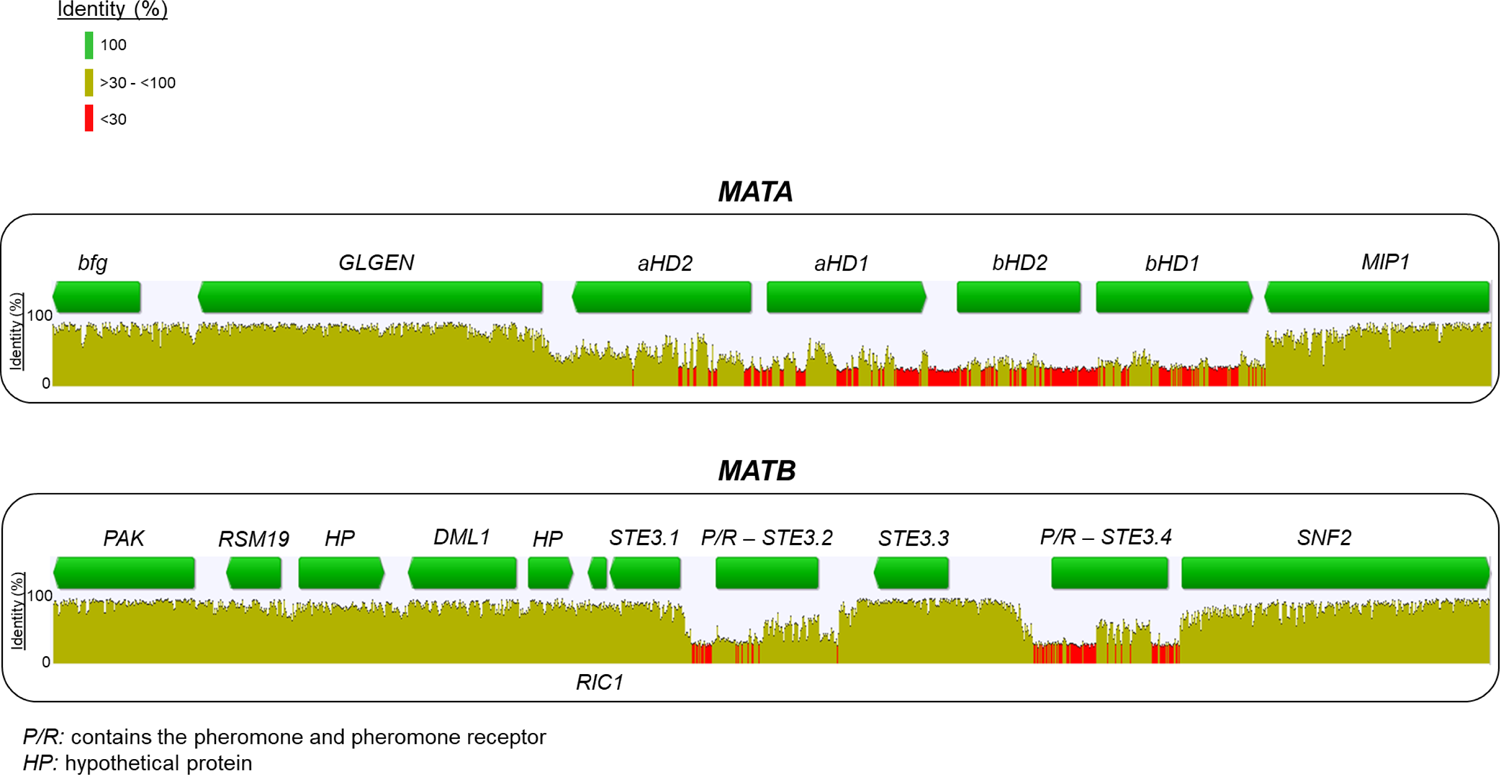
High nucleotide diversity among mating genes. Identity values of nucleotide alignments for *MATA* and *MATB* regions are displayed. Gene arrows indicate the coding direction; however, when gene direction was different (Figure 5) in specimens, we represented a green rectangle. Bar colors represented the level of identity according to the legend.

**Figure 5.**
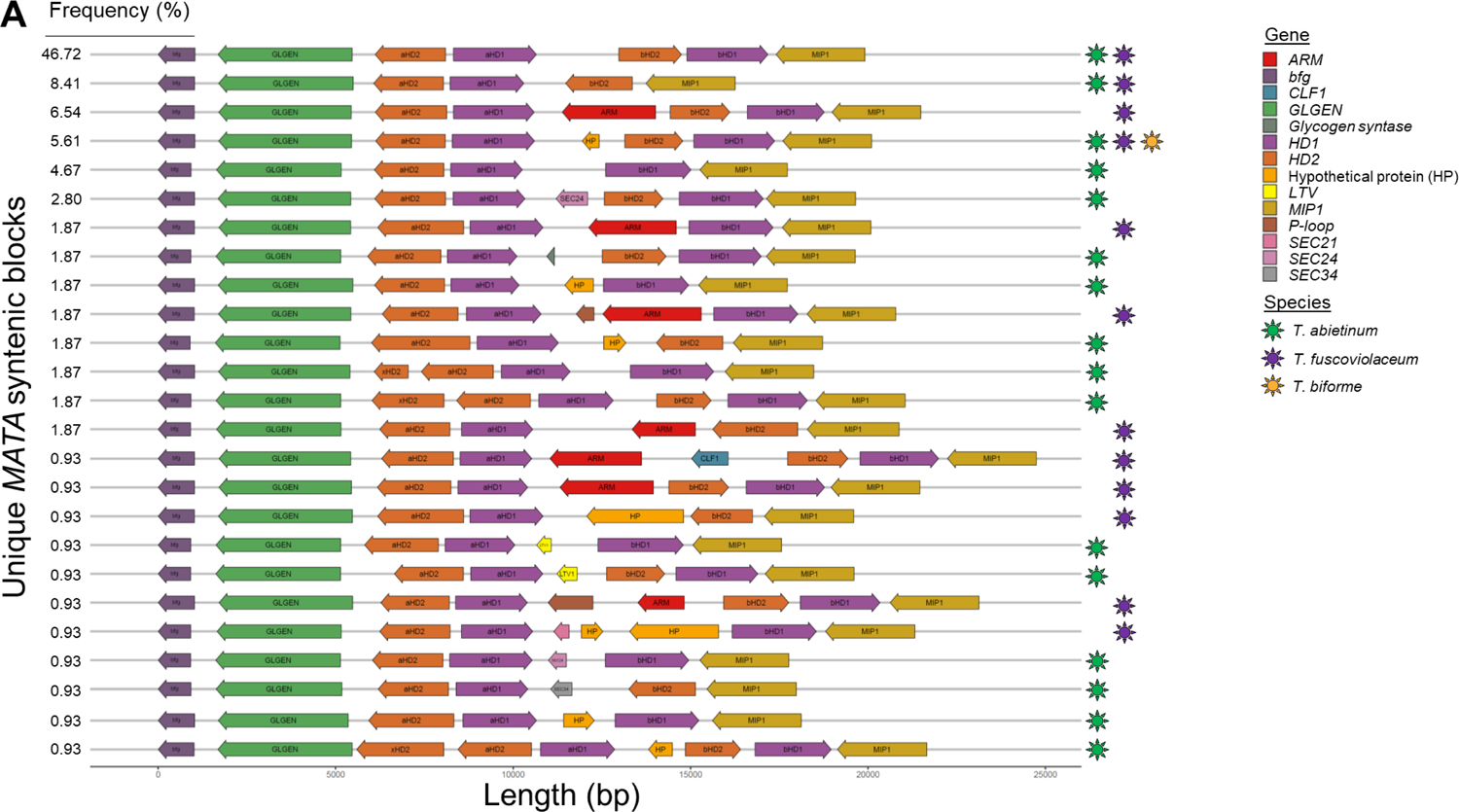
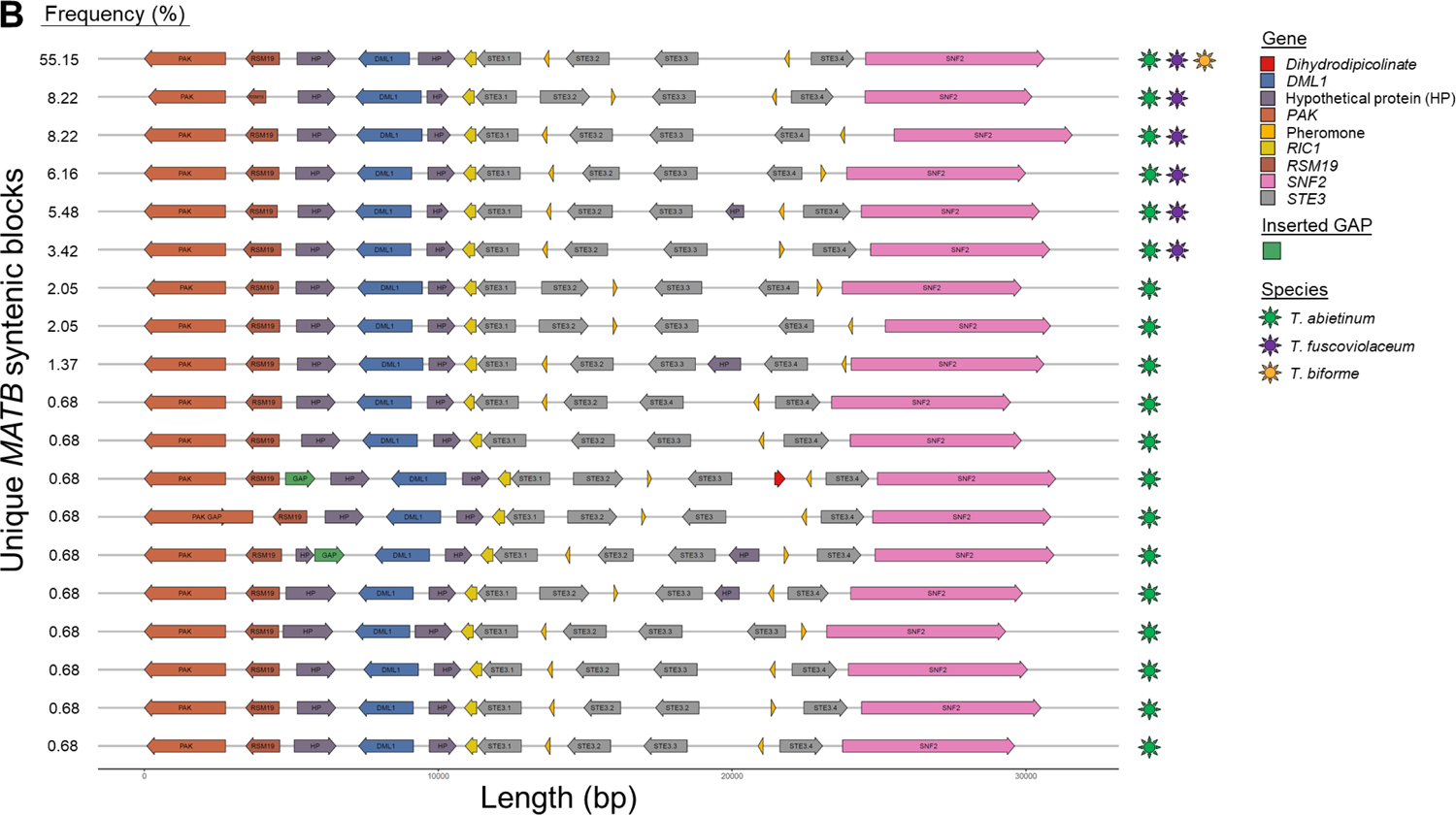
Mating regions are highly dynamic and show multiple rearrangements among *Trichaptum* specimens. Panel A) *MATA* gene order representations for *Trichaptum* specimens with *MATA* genes assembled in one contig. Panel B) *MATB* gene order representations for *Trichaptum* specimens with *MATB* genes assembled in one contig. In the *MATB* case, we considered assembled in one contig when region was assembled contiguously from *RIC1* to *SNF2.* For that reason GAP label is also drawn in this panel B). The percentage of specimens containing a specific *MAT* block order is indicated in the left. Genes were colored according to the legend. Species containing a particular *MAT* block are represented by colored stars at the right of the *MAT* block and were colored according to the legend. Coding sequence direction is represented by the arrows.

We were able to infer several domains and motifs in mating genes. *HD1* and *HD2* homeodomain genes contained three and four exons, respectively, whereas *STE3* genes, characterized by the presence of seven transmembrane domains, included 4 to 6 exons. Homeodomain genes were characterized by the presence of the typical homeobox domain (Figure 2). In each homeodomain protein alignment, we found conserved amino acid sequences in at least 75% of the protein sequences. aHD1 homeobox contained <u>WL</u>X_3_jXN<u>PYP</u>X_4_<u>K</u>X_2_<u>J</u>X_8_KX_4_<u>WF</u>SX_2_<u>RRR</u> and bHD1 <u>WL</u>X_3_jHX<u>PYP</u>X_4_<u>K</u>X_2_<u>J</u>X_13_<u>WF</u>VX_2_<u>RRR</u>, showing highly conserved amino acids between both proteins (underlined amino acids). Similar results were found for aHD2 homeobox domains <u>W</u>X_7_AX_29_<u>FN</u>X_2_YX<u>P</u>X<u>LE</u>XFFXEEQF<u>P</u>SR<u>A</u>DKX_2_<u>LA</u>XKXG<u>M</u>XYR<u>Q</u>IHV<u>WFQNRR</u> and bHD2 <u>W</u>X_29_<u>FN</u>X_4_<u>P</u>X<u>LE</u>X_8_<u>P</u>X_2_<u>A</u>X_4_<u>LA</u>X_2_SX<u>M</u>X_3_<u>Q</u>X_3_<u>WFQN</u>X<u>R</u>X<u>R</u>. In all four types of proteins, it was common to find a tryptophan (W) at the start of the homeobox and two arginines (R) at the end. Inside the homeobox domain, a conservation of a proline (P), a tryptophan and a phenylalanine (F) is likely essential for the activation of the expression of target genes. The nuclear localization signal was detected in HD1 proteins, with the presence of bipartite sequences, <u>KR</u>X_2_SX_8_<u>KR</u> in aHD1 and <u>KR</u>RJX_12_<u>KR</u> in bHD1. Regions enriched in prolines are indicative of putative activation domains (AD), which were conserved in HD2 proteins. The potential AD region contains <u>P</u>XKYP<u>P</u>BFDX_3_D<u>P</u> amino acids in aHD2 and <u>P</u>X_4_<u>P</u>X_2_YPPX_6_F<u>P</u> in bHD2. It is important to note an additional conserved region at the C-terminal of these highly divergent HD1 proteins, where aHD1 contained <u>KL</u>X<u>R</u>INX<u>L</u>LX<u>E</u>AAXLQXEVF amino acids and bHD1 contained <u>KL</u>E<u>R</u>LX_2_<u>L</u>XE<u>E</u>X_3_JXZZEX_2_L. Coiled coils related with heterodimerization were likely located at the N-terminal (Figure 2).

Using the pheromone_seeker.pl script, we were able to detect most of the pheromones. However, some pheromones were not detected due to unexpected amino acids in the CaaX motif (Supplementary Figure 3). We found multiple examples in both pheromones (Phe3.2 and Phe3.4), where the canonical CaaX motif contained a polar amino acid (threonine), displaying an uncommon CpaX motif. Most of the pheromones contained an aspartic acid amino acid following the starting methionine. The presence of both aspartic and glutamic amino acids in the maturation site was highly conserved in *Trichaptum* pheromones.

These results highlight that despite the dynamic nature of both mating regions (Figure 5), where rearrangements and gene losses were frequent, and a high nucleotide diversity (Figure 4), the conservation of functional domains was essential for the activity of mating proteins.

*Phylogenetic analyses demonstrate long-term balancing selection in* HDs *and two* STE3 *genes* To infer the evolutionary history of mating genes and the flanking genes, and to test whether they agree with our species tree (Figure 1B, Supplementary Figure 1), we reconstructed Maximum Likelihood (ML) individual protein trees (Supplementary Figure 4). For most proteins encoded in flanking genes and for both STE3.1 and STE3.3 proteins, phylogenetic trees clustered specimens according to their species designation (Figure 6A,B, Supplementary Figure 4A,B,H-L,N). However, protein trees for homeodomains (aHDs and bHDs), two pheromone receptors (STE3.2 and STE3.4), MIP1 and SNF2 disagreed with the species tree (Figure 6C, Supplementary Figure 4C-G,M,P,O). These trees were characterized by long internal branches and a mixture of species-specific sequences in different clades. All these results pointed to the presence of trans-species polymorphisms due to long-term balancing selection.

**Figure 6.**
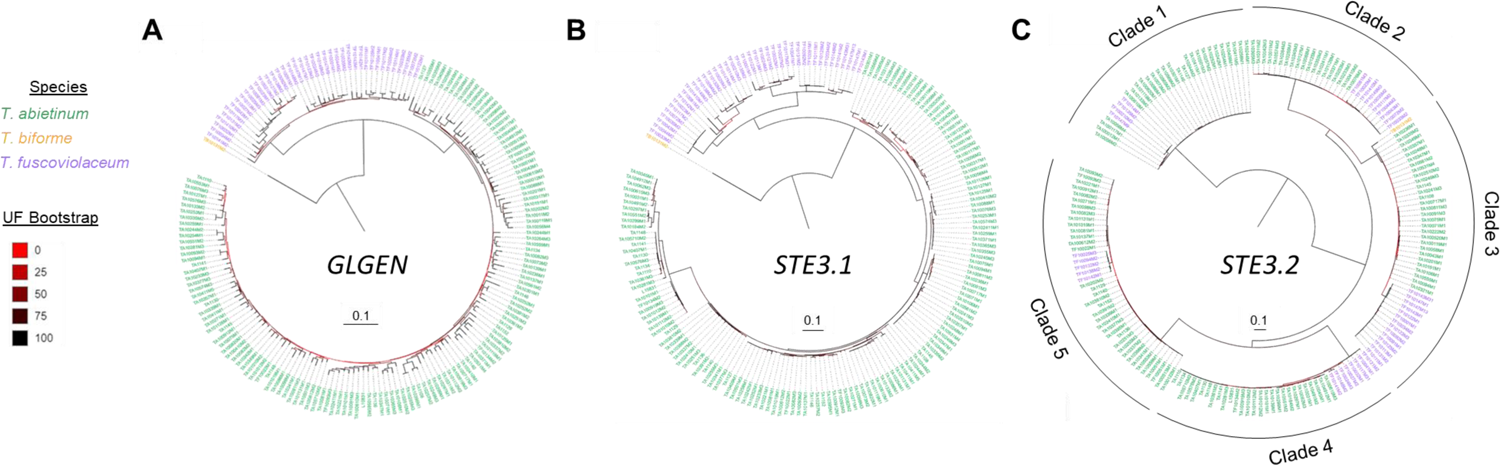
ML phylogenetic tree topology of mating proteins suggests balancing selection and trans-species polymorphisms. ML phylogenetic protein trees of *GLGEN* (a flanking gene), *STE3.1* (a potential non-mating pheromone protein) and *STE3.2* (a mating pheromone receptor protein) are represented in panel A, B and C, respectively. Species designations are indicated by colored bars according to the legend. Branch support was assessed using the ultrafast bootstrap (UF bootstrap) method. UF bootstrap is indicated in each branch by a gradient color according to the legend. Scale bar is represented in number of amino acid substitutions per site. The rest of phylogenetic protein trees and more detailed trees for the represented here are found in Supplementary Figure 4.

To define mating types, we first quantified the number of clades in each phylogenetic tree (see the Material and Methods section for details). The number of clades in the phylogenetic trees varied from 5 to 28. Each clade was considered as a different allelic class for our mating experiments (Supplementary Table 2). Sequences in the same allelic class encoded for proteins with an AAI higher than 86% (Supplementary Figure 5). The highest number of allelic classes was found among alpha complex homeodomain genes where we detected evidence of recombination (Supplementary Table 3). Additionally, once we defined the mating types of our samples, we calculated the AAI by pairwise comparisons of protein sequences of specimens containing the same mating type. We detected high conservation within species for all proteins (AAI = 100%), and higher conservation of pheromone receptors between species (AAI > 95-98%) than for homeodomain genes (AAI > 78-83%), suggesting pheromone receptors were more constrained to accumulate non-synonymous mutations compared to homeodomains (Supplementary Figure 6).

Two alleles were found for the xHD2 protein. A ML phylogenetic tree of all HD2 sequences clustered xHD2 proteins in two aHD2 allelic classes, aHD2.8 and aHD2.10. The limited presence of *xHD2* genes in other specimens and the high similarity of the proteins with two aHD2 proteins points to two recent *aHD2* gene duplications (Supplementary Figure 7). Phylogenetic analyses with other fungal sequences indicated that beta complex HD proteins were much older than Hymenochaetales (Supplementary Figure 8A), which was in accordance with the lower identity values observed for pairwise comparisons within bHD than within aHD (Supplementary Figure 5). Except alpha complex aHD1.12, the rest of aHD proteins were *Trichaptum*-specific. A similar result can be observed for pheromone receptors, where most *Trichaptum* pheromone receptor proteins were closely related, except two proteins, encoded in *STE3.2* and *STE3.4* genes, which were related to pheromone receptor proteins from other fungal species (Supplementary Figure 8B).

The geographic distribution of *MATA* and *MATB* alleles did not suggest a bias towards a particular continent (Supplementary Figure 4, 9), supporting an evolutionary scenario of long-term balancing selection for mating genes.

### Long-term balancing selection left footprints in the mating regions

To further test whether long-term balancing selection is acting on the mating regions, we quantified nucleotide statistics and performed a multilocus HKA test using the mating genes and a collection of universal single-copy orthologs (BUSCO) genes. We first tested the reciprocal monophyletic nature of our BUSCO collection. As expected from the species tree (Supplementary Figure 1B), most of our annotated BUSCO genes (eighty-three percent) showed reciprocal monophyly for both species, *T. abietinum* and *T. fuscoviolaceum*, and 98.64% of the rest of genes (174 genes of 1026 BUSCO genes) showed complete monophyly for one of the two species. This BUSCO dataset suggests a clear diversification of both *Trichaptum* species, and supports the utility of our dataset to set the neutral evolution values of the next analyzed nucleotide statistics.

We observed an elevated number of the average number of synonymous substitutions per synonymous sites (median dS > 1.71) and non-synonymous substitutions per non-synonymous sites (median dN > 0.22) for the mating genes compared to the flanking and BUSCO genes (Supplementary Figure 10, median dS < 0.55, median dN < 0.10). dS and dN values in mating genes were more than 20x and 3x higher than values for BUSCO genes, respectively (Supplementary Table 4). This was an additional support that balancing selection acts on the mating regions. Moreover, similar levels of dS and dN (Supplementary Figure 10, ratio comparison of 0.95-1.03) were observed within and between species in pairwise comparisons of mating genes, indicating that these polymorphisms were not species-specific and recent introgressions were not involved in the generation of trans-species polymorphisms. This was coherent with a scenario where alleles segregated before the diversification of the species. It is important to note that dS and dN values for two putative receptors, *STE3.1* and *STE3.3*, differed from the other mating genes and that they displayed similar low values as most flanking and BUSCO genes (Supplementary Figure 10). In addition, for these two putative non-mating pheromone receptor genes, the dS and dN values were 1.41-3.17 times higher between than within species pairwise comparisons, as we would expect if most of the mutations accumulated after the speciation of *T. abietinum* and *T. fuscoviolaceum*. *MIP1* and *SNF2* dS values were slightly more elevated than BUSCO genes (Supplementary Table 4), but values from between species comparisons were more elevated than within pairwise comparisons (Supplementary Figure 9). This indicates that the elevated dS values are caused by linkage disequilibrium, where the effects of balancing selection in the closest mating gene are not completely broken by recombination.

To infer whether other nucleotide statistics supported balancing selection, we explored gene values deviating from the rest of the genome (Figure 7). Homeodomain (*HD1s* and *HD2s*) and pheromone receptor genes (*STE3.2* and *STE3.4*) deviated from the distribution of 99% of values in at least four nucleotide statistics (elevated pi/dxy ratio, high dS values, low Fst and high Tajima’s D), all in agreement with a balancing selection scenario maintaining trans-species polymorphisms for multiple alleles (Figure 7).

**Figure 7.**
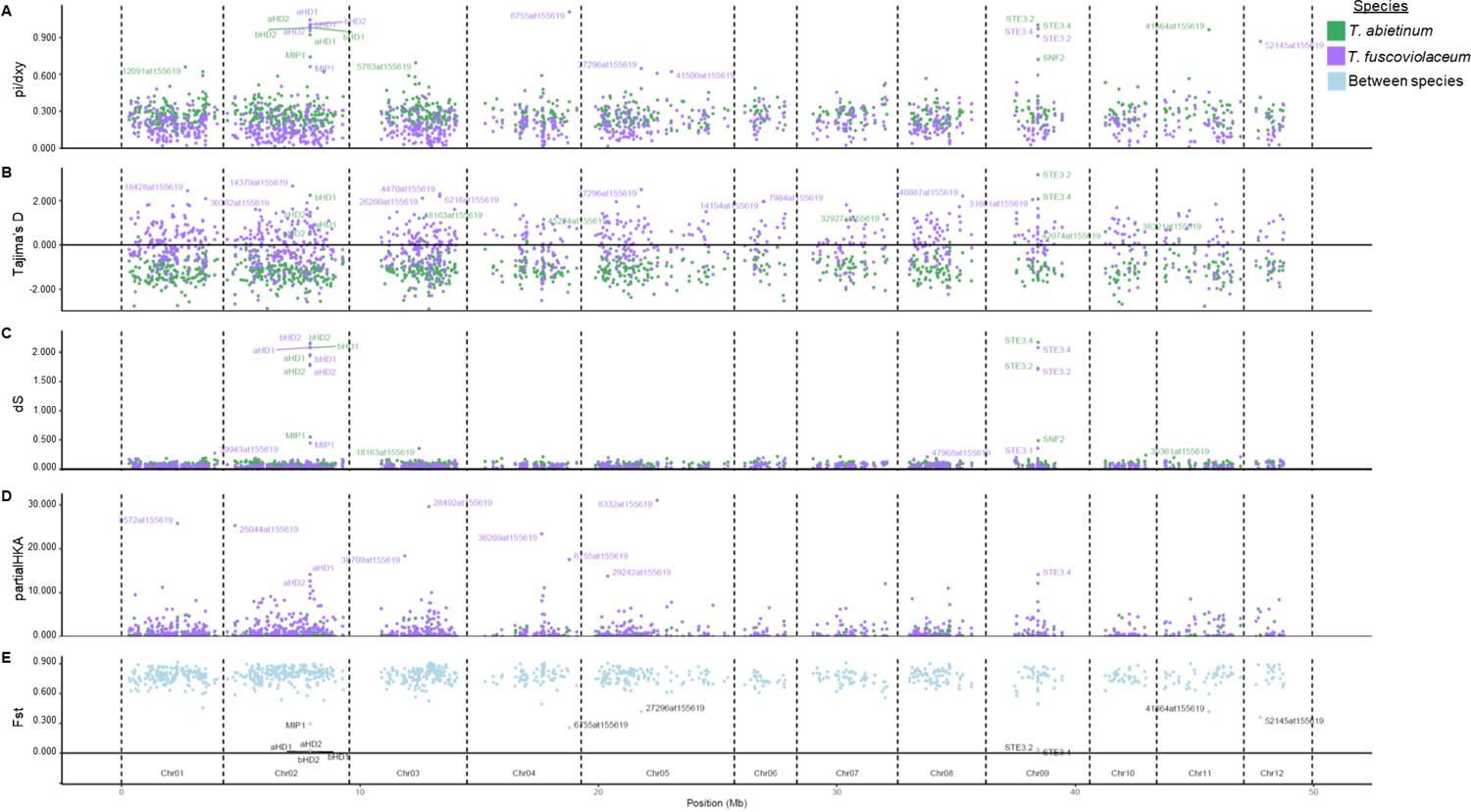
Multiple nucleotide statistics support long-term balancing selection in genes located in the mating region. Nucleotide diversity (Pi), Tajima’s D, average number of synonymous substitutions per synonymous sites (dS), absolute divergence (dxy) and relative divergence (Fst) values are reported in panels A), B), C), E) and F), respectively. Gene contribution to the significance of a HKA test (partial HKA) are represented in panel D). Gene names containing 1% of the highest values (A, B, C, D) or 1% of the lowest values (E, F) are displayed in each panel. *T. fuscoviolaceum* gene names with the highest partial HKA values are displayed due to the significant result of the HKA test (*p*-value = 3.13 x 10^-39^). Each dot represents a gene. Dots were colored according to within species calculations (green or purple for *T. abietinum* and *T. fuscoviolaceum,* respectively) or between species comparison (cyan).

Five BUSCO genes were detected in at least two statistics, deviating from the rest of the genome (Figure 7). Those five genes were also detected to show a phylogenetic topology incongruent with a complete reciprocal monophyly, except 18163at155619 where only *T. fuscoviolaceum* specimens were monophyletic (Supplementary Figure 11). The detected genes encoded for an acetolactate synthase (27296at155619), a ribosomal protein L38e (52145at155619), a non-specific serine/threonine protein kinase (6755at155619), a protein kinase-domain-containing protein (18163at155619) and a NF-kappa-B inhibitor-like protein 1 (41864at155619).

### Distinct mating allelic classes generate compatible mating crosses within species

Based on our allelic class classification (Supplementary Figure 4, Supplementary Table 2, Material and Methods section) we defined the mating types. We tested the outcome of crosses between selected monokaryons from the same species and between species (Supplementary Table 6). We assumed a successful mating when clamp connections were formed (Supplementary Figure 12). Our expectations, based on the molecular characterization, were confirmed in all the performed within species crosses. Crosses using monokaryons with identical *MATA* alleles did not generate clamps when *MATB* alleles were expected to be compatible, and vice versa. These results demonstrate that identical (AAI > 86%) *MAT* alleles generate the first mating barrier. We also included some monokaryons derived from the same dikaryotic isolate (Supplementary Table 2), where most of them showed at least a pair of compatible *MATA* alleles and/or *MATB*. These monokaryons helped us to unfold the original allelic class composition of the parental dikaryon (Supplementary Table 2). Due to the unlinked nature of *MATA* and *MATB* regions and limited number of studied monokaryons from the same dikaryon, some monokaryons had identical mating types, thus did not reveal the original mating type composition of the parental dikaryon.

No clamps were observed in crosses between species with compatible mating types suggesting other mechanisms are involved in the generation of pre-zygotic barriers between *Trichaptum* species.

## Discussion

### Mating genes diversity was maintained by balancing selection

Retaining multiple mating alleles appears to be beneficial as it promotes outcrossing [36]. The multiallelic character of mating types promotes a potential outcross event to occur in 98% of crosses [36, 55]. How this mating diversity originated is not clear, but we demonstrated that some levels of recombination and duplications might play a role. Fifteen recombinant variants in the alpha complex and two recent *aHD2* duplications were detected in *Trichaptum*. It was previously thought that recombination was suppressed or limited in the mating regions [56], and that duplication and diversification events were limited to Agaricales [42]. Recombination is suppressed by the presence of inversions and/or gene losses, which might generate hemizygous specimens, observed in mating loci and genomic regions under balancing selection [57]. The rearrangements observed in our *Trichaptum* beta complex brings another layer of complexity to *MATA* region, which is comparable to the complexity previously described for *MATB* genes [36]. Rearrangements in both *MAT* loci might be an important factor suppressing recombination in these genes. On the contrary, the gene order conservation of the alpha complex does not completely suppress recombination, in accordance with evidence of ongoing recombination between mating genes [58] and their flanking genes in other fungal organisms [59]. Our observations highlight how studying a high number of specimens of the same species, as we have done here for the first time in fungal literature, can unravel previously underestimated mechanisms that generate diversity in mating genes.

We have demonstrated that balancing selection is likely the main force retaining genetic diversity in the mating genes. Evidence of balancing selection has been proposed for homeodomain genes in the pathogenic root decay fungus *Heterobasidion* (Russulales) [26], as well as in pheromone receptors of *Mycrobotryum* species (Mycrobotryales) [24]. The action of balancing selection in *Trichaptum* and in other fungi appears to have occurred before the speciation event, generating multiple cases of trans-species polymorphisms [26]. The genetic signatures of balancing selection highlighted that two pheromone receptors in *Trichaptum* specimens are likely non-mating genes, this could only have been unraveled by including multiple specimes as we have done here. In Agaricomycotina, it is frequent to detect multiple pheromone receptors, some of them not involved in mating functions [40,42,60]. The role of these non-mating pheromone receptors will deserves further investigation.

It has long been speculated about the action of balancing selection in the *MATA* flanking gene, *MIP1* [25, 59]. *MIP1* encodes a mitochondrial intermediate peptidase 1, which is a thiol-dependent metallopeptidase involved in the last step of protein maturation targeted to the mitochondria, where MIP1 cleaves off an octapeptide of immature proteins [61]. The genomic footprints detected in *MIP1* are likely due to the action of linkage disequilibrium, as *MIP1* is close to the beta complex *HD* genes. It has been speculated that *MIP1* signals of balancing selection might be due to a role in mating, such as MIP1 involvement in mitochondrial inheritance, functioning as a suppressor of selfish mtDNA [62]. However, this function is not well-supported. Other genes encoding proteins involved in mitochondrial functions have been found linked to mating genes [59]. In *T. abietinum and T. fuscoviolaceum*, we found *RSM19,* a 37S ribosomal protein S19, linked to *MATB*. However, we did not detect signals of balancing selection in this gene. In addition, some signals of balancing selection were detected in *SNF2*, a gene located in the *MATB* region, encoding a DNA-dependent ATPase protein. The analogous signals of balancing selection between *SNF2* and *MIP1* might support that the balancing selection signal in both genes is due to linkage disequilibrium, and the signal is just a consequence of the action of balancing selection in the neighbor mating genes [59].

### Mating genes and organization resembles other basidiomycetes suggesting similar origin

Sampling and studying the genomes of a wide collection of *Trichaptum* specimens have unraveled the dynamic nature of mating gene architectures. With two homeodomain complexes, *Trichaptum MATA* gene organization is similar to other Hymenochaetales, such as *Phellinus lamaoensis, Phellinus sulphurascens* (both species from the *Phyrrhoderma* genus) and *Schizopora paradoxa* [63]. In other Hymenochaetales species, such as *F. mediterranea* and *Porodaedalea pini*, the location of *GLGEN* gene is more distant and interrupted by multiple *ORFs* [63, 64]. Notably, the *Phyrrhoderma* species and *F. mediterranea* [63, 65] are bipolar, in contrasts to the tetrapolar *Trichaptum* specimens. *Trichaptum* and other Hymenochaetales species, such as *Hypodontia* and *S. paradoxa*, have conserved the ancestral tetrapolar system of basidiomycetes [36]. According to mating studies, the formation of clamp connections is facilitated by the presence of at least one different allele at one of the multiple *MATA HD* complexes and one at the *MATB* P/R loci. Here, we demonstrated by mating experiments and genomic analyses that protein identity must be lower than 86% to function as different mating type, although important protein domains and motifs are conserved.

We inferred that around 224 *MATA* types (28 alpha x 8 beta) and 65 *MATB* types (5 *STE3.2* x 13 *STE3.4*) are present in *Trichaptum* species, which indicates around 14,560 mating types. These numbers are close to the estimated number of alleles, 20,000 mating-types, in a previous study of *T. abietinum* [50], suggesting that our sequencing efforts molecularly characterized most of the *Trichaptum* mating alleles. In other tetrapolar basidiomycete species, such as the model species *Coprinopsis cinerea* and *Schizophyllum commune*, the number of mating types is also similar, around 12,800 (160 *MATA* x 81 *MATB*) and 23,328 (288 *MATA* x 81 *MATB*), respectively [51]. We inferred that beta complex HD alleles were segregating in other Agaricomycetes, suggesting that these HD proteins are much older than alpha HD, a result that is supported by the ongoing recombination events in the alpha HD. Moreover, we cannot discard that alpha complex alleles may be exclusively specific of *Trichaptum*. Allele *aHD1.12* points to potential alpha complex alleles segregating in other Hymenochaetales, but just thirteen Hymenochaetales species have been genome sequenced, and usually only one representative of each species, except for the three sequenced *Pyrrhoderma noxium* specimens. Thus, there are few available genomes to compare.

A new pheromone motif containing a polar amino acid in CaaX motifs was detected by our sequencing efforts. We are not aware of CpaX motifs in other Basidiomycetes, although this motif was observed in pheromones of some Ascomycetes species [66, 67]. The whole genome sequence of other Hymenochaetales and other fungal orders, and the increased number of specimens from multiple species, will clarify the evolutionary history of the alpha complex and protein patterns observed here.

### Trichaptum - a candidate model system for genomics

Using whole genome sequencing, we confirmed the sister species relationship between *T. fuscoviolaceum* and *T. abietinum*, as suggested in previous studies using few molecular markers [68–70]. *Trichaptum biforme* is an early divergent species. Our dataset contributes with a large number of genome assemblies from two non-model species.

The existence of at least two North American intersterility groups (ISGs) that are partially compatible with a third European group in *T. abietinum* indicates three potential differentiated lineages [52–54]. Even though we did not perform a population genomic analysis in this study, multiple well-differentiated clades can be inferred in our ANI and BUSCO phylogenetic species trees, supporting some population structure in our specimen collection. The presence of ISG in *T. fuscoviolaceum* is not previously confirmed based on mating studies [52, 54]. However, we hypothesize that there are at least two potential lineages due to the presence of two well-differentiated *T. fuscoviolaceum* clades, as suggested by Seierstad *et al.* [52]. ANI dissimilarity values between these lineages were nearly as high as values detected in *T. abietinum*, supporting the hypothesis about population structure in *T. fuscoviolaceum*. However, the difference in the levels of populations and the presence of clear ISG in one species and not in the other might be the reason of the differences in the distribution of Tajima’s D values, with more BUSCO genes with negative Tajima’s D values in *T. abietinum* than in *T. fuscoviolaceum*. These results highlight how *Trichaptum* species are suitable for population genomic studies and has the potential to offer new insights into mechanisms of speciation in fungi and how evolutionary mechanisms shape the genome.

## Conclusion

We have demonstrated the importance of sequencing several specimens of fungal species to detect mating-related genes, and to unravel the strength and footprints of long-term balancing selection in mating genes. Events previously thought of as uncommon in mating genes, such as recombination and duplications, have been detected in mating-related genes with conserved gene order. Our *Trichaptum* dataset highlights how diverse and dynamic the mating loci are. These mating genes play a fundamental role in promoting outcrossing events and have consequently been targets of long-term balancing selection. The action of balancing selection leaves signatures of multiple trans-species polymorphisms beyond the genus level. Comparative genomics and phylogenomics were important tools to locate mating genes and characterize the number of alleles retained by balancing selection. Mating proteins with less than 86% identity generated compatible mating types, as we demonstrated by experimental crosses. Despite the number of alleles and the high diversity among them, important domains and motifs are still conserved due to their critical role during the life cycle. Questions regarding the effects of mutations in the interaction between homedomain proteins or receptors and pheromones, especially the presence of non-aliphatic amino acids in the CaaX motif (i.e. a CapX motif), and which role the linked mating genes, such as *MIP1,* are playing during the life cycle are exciting areas of research. Our new sequenced collection of *T. abietinum* and *T. fuscoviolaceum* makes a step-forward to re-establish these fungal organisms as a model system in evolutionary research.

## Material and Methods

### Trichaptum *collection*

A total of 180 *Trichaptum* specimens from the northern hemisphere were included in the study: 138 *T. abietinum*, 41 *T. fuscoviolaceum* and one *T. biforme* (Supplementary Table 6). Specimen GPS coordinate format conversion was generated with `GMScale 0.5.1` to plot the specimen geographic distribution in `R`, using `ggmap 3.0.0`, `ggplot2`, `ggrepel 0.8.2`, and `mapdata 2.3.0`. These specimens were dikaryons (n+n) due to the ability to form fruiting bodies. This result suggest these *Trichaptum* specimens spend most of the time in a dikaryotic state.

### Monokaryon generation and genomic DNA isolation

To facilitate the study of highly diverse regions, such as the mating loci, and to rid out of heterozygosity issues in other genomic regions we established monokaryotic cultures. Monokaryotic cultures were made by hydrating dried field collected specimens in the lab, and collecting single spores that were ejected from these moist fruit bodies onto 3% malt extract agar plates with 10 mg/L tetracyclin, 100 mg/L ampicillin, 25 mg/L streptomycin and 1 mg/L benomyl. Germinated single spores were transferred to new 3% malt extract agar plates with identical mixture of antibiotics and benomyl. Before DNA extraction, monokaryon cultures were grown for 2-3 weeks on nitex nylon (Sefar AG, Heiden, Switzerland) on 3% malt extract agar plates. Two different DNA extraction protocols were used depending on the sequencing protocol. For Illumina sequencing, tissue from 1/4^th^ plate was scraped off the nylon and directly homogenized in 2 ml Lysing Matrix E tubes (MP Biomedicals, Santa Ana, CA, USA) on a FastPrep-24 (MP Biomedicals, Santa Ana, CA, USA) for 2 x 20 seconds at 4.5 m/s^2^. Genomic DNA was extracted using the E.Z.N.A HP Fungal DNA kit (Omega Bio-Tek, Norcross, GA, USA) supplemented with 30 µl RNaseA (Qiagen, Hilden, Germany). For PacBio sequencing, tissue from 10 plates were scraped off the nylon and directly homogenized in a mortar with liquid N_2_. Genomic DNA was extracted using a phenol-chloroform protocol followed by a macro (500 µg) Genomic tip (Qiagen, Hilden Germany) protocol, as described in Skrede *et al* [71].

### Genome sequencing and assembly

In order to get the chromosome location and sequences of mating genes, we first Illumina sequenced and provided the first PacBio sequences for the *Trichaptum* genus. We sequenced one specimen from *T. abietinum* (TA-1010-6-M1) and one from *T. fuscoviolaceum* (TF-1002-10-M3) (Supplementary Table 1).

Illumina libraries were generated by the Norwegian Sequencing Centre using the following protocol: 1 µg of genomic DNA was sheared using 96 microTUBE-50 AFA Fiber plates (Covaris Inc., Woburn, MA, USA) on a Covaris E220 system (Covaris Inc., Woburn, MA, USA). The target fragment size was 300-400 bp. gDNA samples were cleaned on a small volume Mosquito liquid handler (TTP labtech) with a 1:1 ratio of Kapa Pure beads (Roche, Basel, Switzerland) and eluted in Tris-Cl, pH 8.0. Library preparation was carried out with 500 ng sheared DNA using Kapa Hyper library prep kit (Roche, Basel, Switzerland). Barcodes were added using the Illumina UD 96 index kit (Illumina). Final libraries were PCR-amplified during 5 cycles with Kapa HIFI PCR kit (Roche, Basel, Switzerland) before standard library quality control with standard sensitivity NGS Fragment kit (Agilent, Santa Clara, CA, USA). Quantification was performed in a qPCR with Kapa Library quantification kit (Roche, Basel, Switzerland). The first batch of library specimens were sequenced with HiSeq 4000 system, and the second with NovaSeq I (Supplementary Table 1). 2×150 paired-end Illumina reads were generated by both systems. Barcodes and adapters were trimmed from final Illumina sequences using `Trim_galore 0.6.5` [72].

### PacBio libraries were prepared by the Norwegian Sequencing Centre using Pacific

Biosciences Express library preparation protocol (Pacific Biosciences of California, Inc, USA) without any prior fragmentation. Size selection of the final PacBio libraries was performed using BluePippin (Sage Science, Beverly, USA) and 15 Kbp cut-off. PacBio libraries were sequenced on one 1M SMRT cell using Sequel Polymerase v3.0 and sequencing chemistry v3.0. Loading was performed by diffusion and movie time was 600 min for *T. abietinum* and 900 min for both *T. fuscoviolaceum* runs.

We assembled the genome of *T. abietinum* using PacBio reads by different assemblers: `Flye 2.6` [73], `Canu 1.9` [74], `MECAT2` [75], `SMARTdenovo 1.0.0` [76] and `wtdbg2 2.5` [77]. Quality of the draft PacBio genome and percentage of consensus between draft genome and Illumina reads were quantified by `quast 5.0.2` [78] and `polca` [79], respectively. The best draft PacBio assembly based on the previous quality statistics was selected and Illumina-corrected using `HyPo` [80]. Scaffolds with less than 100 PacBio reads of support and less than 10 Kbp of length were removed from the final corrected genome assembly. *T. abietinum* ultrascaffolding was done using a Hymenochaetales species, *P. noxium* KPN91 PacBio genome assembly, as reference (Accession number GCA002287475, [81]. We first checked chromosome correspondence using `D-Genies` [82] and manually ultrascaffolded in `Geneious 6.1.6` [83]. Chromosomes were named according to *P. noxium* chromosome similarity. We applied the same pipeline to the *T. fuscoviolaceum* PacBio assembly, except that ultrascaffolding was performed using `RaGOO` [84], and the *T. abietinum* PacBio genome assembly as reference. Visual inspection of syntenic comparisons were performed using `mummer 3.23` [85] and `D-Genies`. This approach allowed us to correct the order of the ultrascaffolded chromosome 3 of *T. abietinum*. We assumed that the chromosome 3 order must be more similar between sister-species *T. abietinum* and *T. fuscoviolaceum* than between *T. abietinum* and *P. noxium*. In both *Trichaptum* assemblies, ultrascaffolded chromosomes contain artificial 10,000 Ns separating joined scaffolds. Assembly statistics of the final genomes, such as N50, genome size, and completeness of universal single copy orthologous genes, were assessed using `quast` and `BUSCO 4.1.2` [86]. The training `BUSCO` database was `agaricomycetes_odb10`, which contains 2898 genes.

Genomes of the 178 specimens, sequenced by the Illumina platform, were assembled with `iWGS` wrapper [87]. We selected `SPAdes 3.14` [88] assemblies based on `quast` quality reports. Genome completeness was assessed with `BUSCO`. In addition, we included a DOE Joint Genome Institute (JGI) MycoCosm Illumina-sequenced and assembled *T. abietinum* specimen (L15831, [89].

### Trichaptum species classification and species tree reconstruction

Species designation of our specimens was first supported based on a fast method, `fastANI 1.1` [90]. With `fastANI,` we calculated the pairwise average nucleotide identity (ANI) among genome assemblies, whose values were then converted to a percentage dissimilarity matrix by subtracting ANI from a value of 100%. The dissimilarity data was used as distance to reconstruct a Neighbor-Joining (NJ) phylogenetic tree in `MEGA v5` [91].

The utilization of gene nucleotide and amino acid sequences of universal single copy orthologs annotated with `BUSCO` assessed the species designation by `fastANI`. Individual `BUSCO` protein alignments were generated with `MAFFT 7.455` [92]. Amino acid alignments were back translated to nucleotides using `pal2nal v14` [93]. Codon columns with gaps were removed from the alignments using `trimal 1.4.1` [94]. Gene sequences present in all specimens that retained at least 30% of positions and with more than 300 nucleotides (100 amino acids) were selected for additional analyses. In total, 1026 BUSCO genes (35% of the genes) passed our filters. Maximum Likelihood (ML) phylogenetic trees of trimmed genes were reconstructed using `IQTree 2.0.3` [95]. The best fitted evolutionary nucleotide model for each gene was estimated by `ModelFinder` [96] implemented in `IQTree`. Individual gene trees were pooled in a unique file, which was the input to reconstruct the species tree by applying a coalescent model implemented in `ASTRAL 5.7.4` [97]. Species tree branch support was assessed by calculating the gene concordance factor implemented in `IQTree`. To assess reciprocal monophyly of BUSCO genes, ML phylogenetic trees were read in R using `treeio v1.12` [98] and converted to `ape v5.4` format [99]. Once species designation were associated to phylogenetic tip labels, the trees were rooted using *T. biforme* specimen as an outgroup. Monophyly test was performed using `spider v1.5` [100]. ML phylogenetic trees of BUSCO genes detected as top 1% in at least two nucleotide diversity statistics (see below) were drawn to a pdf using `ggtree v2.2.4` [101].

### Mating gene annotation, alignments and phylogenetics

Mating regions encoding the genes involved in the sexual cycle are conserved among basidiomycetes [36]. We first searched for conserved flanking genes to delimit the mating sites in our new PacBio genomes. Mating A (*MATA*) region was located using *MIP1* (mtDNA intermediate peptidase), *bfg* (beta-flanking gene) and *GLGEN* (Glycogenin-1) gene sequences. Mating B (*MATB*) region was delimited using *PAK* (syn. *CLA4*, serine/threonine protein kinase). We found both mating regions by performing a `blast` search in `Geneious` [102] using *P. noxium* flanking gene sequences as subject. Delimitation of genes and coding sequences in mating regions were performed using `FGENESH` and the *P. noxium* gene-finding parameters [103]. Some annotated open reading frames (ORFs) required manual curation. ORFs were `blastx` in NCBI to confirm the gene designation. An additional annotation comparison to infer the number of exons in different ORFs was done using `MAKER2` [104], where we included the transcriptome dataset of L15831 *T. abietinum* as input [89].

The annotation of domains and motifs was performed using different strategies. Typical homeodomain/homeobox domains in HD proteins were annotated with `CD-search` using the CDD v3.18 – 55570 PSSMs database [105]. To differentiate *HD1* and *HD2* genes, we first screened the nuclear localization signal (NLS) domain using `NLS Mapper` [106]. NLS is characteristic of HD1 proteins [39,107,108]. Conserved regions enriched in proline amino acids were suggested as potential regions for activation domains (AD) for homeodomain proteins [109]. Coiled coil regions involved in the dimerization of the two homeodomain proteins were detected with `Coiled coils v1.1.1 Geneious plugin`. Proteins with seven-transmembrane G protein-coupled receptor superfamily domains are usually indicative of *STE3* pheromone receptors [110]. The 7 transmembrane domains of the pheromone receptor protein were annotated with `PredictProtein` [111]. Pheromones were screened in close proximity to the detected pheromone receptors using `pheromone_seeker.pl` script [112]. Briefly, the perl script searches common aminoacid features in pheromones, such as the CaaX (C, cysteine; aa, two aliphatic amino acids; X is any amino acid) motif in the C-terminal of the pheromone [40, 60]. Hits with a length shorter than 100 bp or longer than 200 bp, and/or distant to *STE3* genes were considered as false positives. Consequently, we removed those hits from the annotations. Additionally, pheromones in specimens missing at least one hit close to *STE3.2* or *STE3.4* were manually searched using pheromone amino acid sequences of specimens in the same clade for *STE3.2* or *STE3.4* phylogenetic trees. Pheromone maturation sites were located by searching glutamic/arginine (ER) or aspartic acid/arginine (DR) amino acid motifs [39].

Once we had annotated the mating regions in our PacBio reference specimens, we were able to search for these genes in the Illumina sequenced and assembled genomes of the rest of specimens. We first generated local blast databases for our Illumina genomes. We BLASTed the reference flanking genes to pull out the mating regions. In case a mating region (*MATA* or *MATB*) was not contiguous (<44% and <20% of specimens for *MATA* and *MATB*, respectively), but split on different contigs, we assumed those regions kept the same gene order as in the PacBio reference genomes, and we ultrascaffolded the contigs for each mating region accordingly. 999 Ns were added between joined contigs. Similar to the PacBio genome assemblies, we defined the mating regions to the scaffold/ultrascaffolded segment containing sequences from *bfg* to *MIP1* for *MATA* region, and from *PAK* to *SNF2* for *MATB*. Once regions were located and/or ultrascaffolded, we used the previous `FGENESH` pipeline for annotating ORFs. Gene identification was performed by BLASTing the genes from our reference genomes against the mating regions. Additional identification was performed by searching family matches in the InterPro-5-RC6 database [113]. All annotations were stored in gff3 files generated by `Geneious.` Due to limitations of our Illumina sequencing some genes in the mating regions were not detected probably because they were not covered by the Illumina reads.

For calculating the frequency of each unique gene block for each region, we followed a conservative approach. We took into account only mating regions that were assembled contiguously by `SPAdes` and did not need an ultrascaffolding step. The criteria applies from *bfg* to *MIP1* (*MATA*) and from *RIC1* to *SNF2* (*MATB*) genes. Gff3 files were the input to plot *MAT* gene order in `R` using `dplyr 1.0.2`, `gggenes 3.3.2`, `ggplot2 3.3.2`, and `rtracklayer 1.48.0`.

To calculate the nucleotide identity conservation of mating regions, we first aligned *MATA* and *MATB* sequence regions independently using FFT-NS-1 algorithm, 200PAM/k=2 score matrix and default gap opening penalty and offset value with the `MAFFT 7.017` version implemented in `Geneious`. Gaps present in more than 20% of specimens were removed with `trimal`. Identity plots for each region was generated in `Geneious`.

For phylogenetics, we first generated amino acid sequence alignments using `MAFFT` and back translated to nucleotides with `pal2nal`. Again, we were conservatives and codon columns with gaps were removed from the alignments using `trimal.` The trimmed alignment was converted to amino acid for ML phylogenetic tree reconstruction with `IQTree`. An evolutionary protein model for each protein was estimated by `ModelFinder`. Homeodomain and pheromone receptors were classified in clades/alleles according to visual inspection of ML phylogenetic trees and pairwise amino acid identity percentages calculated in `Geneious`. Note here that alleles/allelic classes refer to similar protein sequences enclosed in a clade and not to haplotype sequences.

Mating genes, flanking genes and the species tree were plotted with `iTOL 5.7` [114]. *T. biforme* was used as the outgroup to root the trees when possible. To detect whether a mating related gene was segregating before the speciation event, we selected a random protein sequence of each allelic class to infer the phylogenetic relationship with proteins from other Hymenochaetales species, two reference species of Agaricales and one species from Polyporales.

### Nucleotide statistics, tests to detect balancing selection and recombination

Trimmed codon-based sequence alignments of mating genes, their flanking genes and BUSCO genes were the input for the calculation of nucleotide statistics. Pairwise sequence estimation of synonymous and nonsynonymous substitution rates were calculated using the model of Yang and Nielsen [115] implemented in the `yn00` program of `PAML 4.9` [116]. We calculated nucleotide statistics, absolute nucleotide divergence (dxy) and relative divergence (Fst) using the `PopGenome 2.7.5` package in `R 4.0.2` [117]. Sequences were split in different alignments based on the species designation inferred from the species tree phylogeny. Each species-specific alignment was the input to calculate nucleotide diversity (π, Pi) and Tajima’s D using `PopGenome`. A multilocus test for detecting balancing selection was performed with `HKAdirect 0.70b` [13]. We generated species-specific input tables for `HKAdirect` using `PopGenome.` The input tables consisted of the number of samples (nsam), segregating sites (S), absolute divergence (Divergence) and length for each species-specific gene (length_pol and length_div). We set factor_chrm to 1 because our genes are encoded in the nuclear genome. The input tables were necessary to run the multilocus test.

dS and dN boxplots, and genome-wide gene nucleotide statistic plots were generated in `R` using `cowplot 1.0.0`, `dplyr`, `ggplot2`, `ggrepel`, `PopGenome`, `reshape2 1.4.4`, and `rtracklayer`.

To detect evidence of recombination, homeodomain and pheromone receptor individual nucleotide alignments were analyzed in `RDPv4` [118]. Recombination events significantly detected by all seven methods (`RDP`, `GENECONV`, `Bootscan`, `Maxchi`, `Chimaera`, `SiSscan` and `3Seq`) were reported.

### Monokaryon specimen crosses

To test the compatibility of allelic classes designation for *MATA* and *MATB* alleles, we designed putative compatible and incompatible mating type crosses (Supplementary Table 5). *MAT*A mating type is defined by the allelic class classification of the two complexes. *HD1* and *HD2* genes of the *MATA* complex were highly linked, so they can be treated as a unique unit. For example, aHD1.9 and aHD2.1 defined alpha*MATA*-1. Now, the combination of alpha*MATA-1* with a beta complex can give different *MATA* mating types. For example, *MATA-2* was defined by alpha*MATA-1* plus beta*MATA*-2. There are 28 alpha and 8 beta complex allelic classes generating around 224 *MATA* types. Similarly, *MATB* mating type is defined by the allelic classes of the pheromone receptors, i.e *MATB-52* is defined by *STE3.2-5* and *STE3.4-10.* There are 5 and 13 *STE3.2* and *STE3.4* pheromone receptor allelic classes, respectively, generating around 65 *MATB* types. Finally, a mating type *MAT-2* is generated by the combination of *MATA-2* and *MATB-52*. Due to the presence of 224 *MATA* and 65 *MATB,* this suggest around 14,560 mating types. The mating classification numbering is arbitrary. For that reason, for simplicity, our selected candidates were described as having or not having a compatible alpha/beta complex and *STE3.2/STE3.4* (Supplementary Table 6). We expected a compatible cross when one of the *MATA* complexes and one of the pheromone receptors were from distinct allelic classes among the selected specimens.

A total of 21 and 10 crosses were designed for crosses within *T. abietinum* and *T. fuscoviolaceum*, respectively, and 10 crosses between both species. Crosses were performed by plating monokaryons on 3% malt extract agar plates at 4 cm distance between the two monokaryons. After 2-4 weeks, hyphal growth generated contact zones between both monokaryons. Then, a small piece from the middle area of the contact zone was extracted and re-plated on a new 3% malt extract agar plate. After one week of growth, we examined clamp connections by placing a sample of the culture on a slide under a Nikon Eclipse 50i (Nikon Instruments Europe BV, Amsterdam Netherlands). Images of the microscopic slides were acquired under a Zeiss Axioplan-2 imaging with Axiocam HRc microscope camera (Zeiss, Oberkochen Germany). All crosses were performed in triplicates.

### Bioinformatic tools

All bioinformatic tools, programs and most scripts were implemented in UNINETT Sigma2 SAGA High-Performance Computing system (technical details here: https://bit.ly/2VkIXM2), except most `R` steps. `R` analyses were performed in Windows 10 operative system, implemented in `RStudio 1.3.1073` with an `R version 4.0.2`. Bioinformatic tools were installed through `conda` [119] under the SAGA module `Anaconda2/2019.03.` Non-computational demanding and/or simple python steps were implemented in `Jupyter notebooks` using python modules installed through `conda` under `Windows 10 Anaconda 1.9.12` version.

## Acknowledgments

We thank Sebastián Ramos Onsins for the interpretation of the results provided by his program HKADirect and Alija Bajro Mujic for sharing pheromone_seeker.pl. We thank Amanda Bremner, Beatrice Senn-Irlet, Buck Castillo, Brittny Gardner, Carolina Girometta, Carolina Pina Paez, Charlotte Johnson, Daniel Andrew Lovejoy, Daniel Luoma, Hermann Voglmayr, Irmgard Krisai-Greilhuber, Jilian Myers, Jonas Oliva, Jørn-Henrik Sønstebø, Kadri Runnel, Kevin Amses, Kyle Gervers, Myung Soo Park, Otto Miettinen, Rabern Simmons, Rebecca Clemons, Sergey Volobuev, Stefan Blaser, Sara Lynch, Stephen R. Clayden, Sundy Maurice, Ursula Peintner, Vesa Salonen, Young Woon Lim and Yu-Cheng Dai for providing samples and assistance in the field. We thank Georgiana May for critical discussion about the strength of balancing selection. The sequencing service was provided by the Norwegian Sequencing Centre (NSC, www.sequencing.uio.no). NSC is a national technological platform hosted by the University of Oslo and supported by the “Functional Genomics” and “Infrastructure” programs of the Research Council of Norway and the Southeastern Regional Health Authorities. The computations were performed on resources provided by UNINETT Sigma2 - the National Infrastructure for High Performance Computing and Data Storage in Norway. This work was supported by Research Council of Norway (RCN) grant no. RCN 274337. D.P. is a postdoctoral researcher funded by the RCN grant no. RCN 274337 and a senior researcher, supported by the Valencian International University (VIU). The funders had no role in study design, data collection and analysis, decision to publish, or preparation of the manuscript.

## Declarations

Ethics approval and consent to participate

Not applicable

Consent for publication

Not applicable

Availability of data and materials

PacBio and Illumina sequencing data have been deposited in NCBI’s SRA database, Bioproject PRJNA679164. Illumina genome assemblies, MAT regions and their annotations (gff files), together with the source data underlying Figure 5, Supplementary Figs. 4,5, 7, 11 and the details about command lines used to run the programs can be found at https://perisd.github.io/TriMAT/ and Dryad repository <u>doi:10.5061/dryad.fxpnvx0t4</u>. PacBio genomes were submitted to the European Nucleotide Archive (ENA) under the project number PRJEB45061. Phylogenetic trees can be accessed following the `iTOL` link http://itol.embl.de/shared/Peris_D. All other relevant data are available from the authors upon reasonable request.

## Competing interests

DP declares receiving royalties from VIU based on publication productivity.

## Authors’ contributions

Conceived and designed the experiments: D.P., I.S., H.K., D.S.L. Collected specimens in the field, performed spore isolation and specimen identification I.S., D.S.L, V.B.K., I.S.M. Performed genomic DNA extraction V.B.K., I.S.M., I.S., D.S.L, M.S.D. Performed specimen crosses D.S.L., V.B.K. Computational analyses, plotting and statistics were performed by D.P. Wrote a first draft of the paper D.P., D.S.L., I.S. All authors read, commented and approved the final manuscript.

## Supporting information captions

**Supplementary Figure 1.** Phylogenetic trees suggest some population structure in *Trichaptum* species. A) Neighbor-Joining tree using the 100 – ANI values as distances to reconstruct the tree. Scale bar represents 100 – ANI / 100. B) Coalescent species tree using 1026 BUSCO ML phylogenetic trees. Scale bar represents coalescent units. Bar colors represent the species designation according to the legend. Circles in branches represent the concordance factor support (0: none ML tree agrees – 100: all 1028 ML trees agree).

**Supplementary Figure 2.** Genomes of *T. abietinum* and *T. fuscoviolaceum* are mostly syntenic. `D-G enies` dot-plot of our two reference PacBio genomes. Alignment matches are represented by dots and the identity values are colored according to the legend. MAT region locations are indicated.

**Supplementary Figure 3.** Non-common CpaX motifs were detected in *Trichaptum* **pheromone proteins.** Phe3.2 and Phe3.4 sequence alignments of unique pheromone proteins are represented in panel A) and B). Sequence logo is represented at the top of each alignment to highlight conserved amino acids. Polar amino acids in the CaaX motif are squared in red.

**Supplementary Figure 4.** ML phylogenetic trees reconstruction of individual proteins shows signals of balancing selection in mating genes and linked genes. ML phylogenetic trees of individual proteins from the *MATA* and *MATB* regions are represented. Species designation and continental isolation are indicated by colored bars according to the legend. Branch support was assessed using the ultrafast bootstrap (UF bootstrap) method. UF bootstrap is indicated in each branch by a gradient color according to the legend. Scale bar is represented in number of amino acid substitutions per site.

**Supplementary Figure 5.** Pairwise amino acid identity within mating proteins. Pairwise amino acid identity was calculated for protein sequences within a clade/allele and between protein sequences from different clades (allelic classes). Dots represent the average value for within or between pairwise comparisons. Median values for all proteins are represented by horizontal lines inside the boxes, and the upper and lower whiskers represent the highest and lowest values of the 1.5 * IQR (inter-quartile range), respectively. Box plots and dots were colored according to the species where the pairwise comparison was performed. Horizontal dashed line represents the maximum value of 100 - % amino acid identity. We considered 86% amino acid identity a threshold to classify sequences in an allelic class.

**Supplementary Figure 6.** Pairwise amino acid identity of mating proteins from specimens with identical mating types. Pairwise amino acid identity was calculated for protein sequences within an allelic class of the same species (2 pairwise comparison for *T. fuscoviolaceum*) and between species (2 pairwise comparisons between 2 *T. abietinum* and 2 *T. fuscoviolaceum*). Dots represent the average value for within or between pairwise comparisons. Horizontal dashed line represents the 86% amino acid identity threshold detected in Supplementary Figure 5.

**Supplementary Figure 7.** Two recent duplications of *aHD2* genes generated xHD2 proteins. ML phylogenetic trees of a protein sequence alignment containing xHD2, aHD2 and bHD2. xHD2 sequences are highlighted with red arrows. Branch support was assessed using the ultrafast bootstrap (UF bootstrap) method. UF bootstrap is indicated in each branch by a gradient color according to the legend. Scale bar is represented in number of amino acid substitutions per site.

**Supplementary Figure 8.** Some mating alleles are older than *Trichaptum* genus. Selected regions of ML phylogenetic trees of trimmed (`trimal –gt 0.8`) protein sequence alignments containing HD2-HD1 and STE3 are displayed in panel A) and B), respectively. Branch support was assessed using the ultrafast boostrap (UF bootstrap) method. UF bootrstrap is indicated in each branch by a gradient color according to the legend. Scale bar is represented in number of amino acid substitutions per site. *Trichaptum* proteins are highlighted by red arrows or enclosed in a red bar.

Protein sequences were retrieved from DOE-JGI MycoCosm and download from NCBI as indicated:

1. Hymneochaetales JGI protein list: Fomme: *Fomitiporia mediterranea* (MF3/22), Onnsc: *Onnia scaura* (P-53A), Phefer: *Phellinidium ferrugineofuscum* (SpK3Phefer14), Pheign: *Phellinus ignarius* (CCBS575), Phevit: *Phellinus viticola* (PhevitSig-SM15), Pheni: *Phellopilus* (*Phellinus*) *nigrolimitatus* (SigPhenig9), Porchr: *Porodaedalea chrysoloma* (FP-135951), Pornie: *Porodaedalea niemelaei* (PN71-100-IP13), Resbic: *Resinicium bicolor* (OMC78), Ricfib: *Rickenella fibula* (HBK330-10), Ricmel: *Rickenella mellea* (SZMC22713), Schpa: *Schizopora paradoxa* (KUC8140), Sidvul: *Sidera vulgaris* (OMC1730).
2. Downloaded from NCBI: [HYMENOCHAETALES] *Fomitiporia mediterranea* (MF3/22), *Pyrrhoderma noxium* (KPN91), *Shanghuangporus baumii* (Bpt 821), *Rickenella mellea* (SZMC22713); [AGARICALES] *Laccaria bicolor* (S238N-H82), *Coprinopsis cinerea* (Okayama7#130); [POLYPORALES] *Rhodonia* (*Postia*) *placenta* (Mad-698-R).

To remove protein redundancy in protein collection of species retrieved from JGI, a blastp using the downloaded NCBI protein sequences and our HDs and STE3s protein representatives of each clade/allele was performed. For each input sequence two hits were used for sequence alignments, a protein sequence with the lowest e-value and the protein sequence with the highest coverage value. Complete ML phylogenetic trees are deposited in a shared iTOL folder: https://itol.embl.de/shared/Peris_D

**Supplementary Figure 9.** Geographic distribution of mating alleles supports long-term segregation. Stacked bar plots are represented for each mating gene. Bars were colored according to their geographic location.

Supplementary Figure 10. dS and dN values for mating, flanking and BUSCO genes supports balancing selection in mating genes. Panels A), C) reports the pairwise dS within each species (colored according to the legend) or between species (black) for each gene in the *MATA* and *MATB* regions, respectively. Similarly, panels B), D) reports the pairwise dN. Median values for all genes are represented by horizontal lines inside the boxes, and the upper and lower whiskers represent the highest and lowest values of the 1.5 * IQR (inter-quartile range), respectively. Median values for BUSCO genes are represented by horizontal dashed lines and they are colored according to the legend, green and purple for within *T. abietinum* and *T. fuscoviolaceum* comparisons, respectively, and black between species comparisons.

**Supplementary Figure 11.** Detected BUSCO genes are shown to have some signal of non-reciprocal monophyly. Maximum-Likelihood phylogenetic trees of five detected BUSCO genes based on nucleotide statistics (Figure 7) are represented. Scale bar is represented in number of nucleotide substitutions per site.

**Supplementary Figure 12.** Allelic class classification based on phylogenetics and protein identity generated compatible and incompatible crosses. Example plate and microscope pictures of the specimen cross experiments are displayed. The rest of the pictures indicated in Supplementary Table 5 can be found in https://perisd.github.io/TriMAT/. When types were distinct in both mating loci clamp connections are observed in septae.

## Notes

https://perisd.github.io/TriMAT/

https://itol.embl.de/shared/Peris_D

doi:10.5061/dryad.fxpnvx0t4

### Summary of Updates

A short description of the main changes performed to the previous manuscript below: 1.To highlight our main findings, we modified the title to 'Molecular diversity maintained by long-term balancing selection in mating loci defines multiple mating types in fungi' where the molecular characterization of mating loci and the phenotypic validation of compatible mating types were the most novel results of our manuscript. 2.We clearly highlighted that our organism is a non-model Basidiomycete (lines 127-128, 139-140), and the previous knowledge about its tetrapolar character and number of mating types were just inferred from experimental crosses with no molecular validation (lines 133-138). The molecular characterization is performed in our study. 3.Expanded details about the Trichaptum life cycle (new Figure 1A) and we added clarity about how we defined allelic classes, how allelic classes were linked to mating types (lines 103-107, 247-251, 686-702) and highlighted that they were validated by experimental crosses (lines 325-343). 4.Clarification of text and improvement of figure details were also provided.

