## Supplementary_Figures_Tables for "Molecular diversity maintained by long-term balancing selection in mating loci defines multiple mating types in fungi": Supplementary Figure 4.pdf

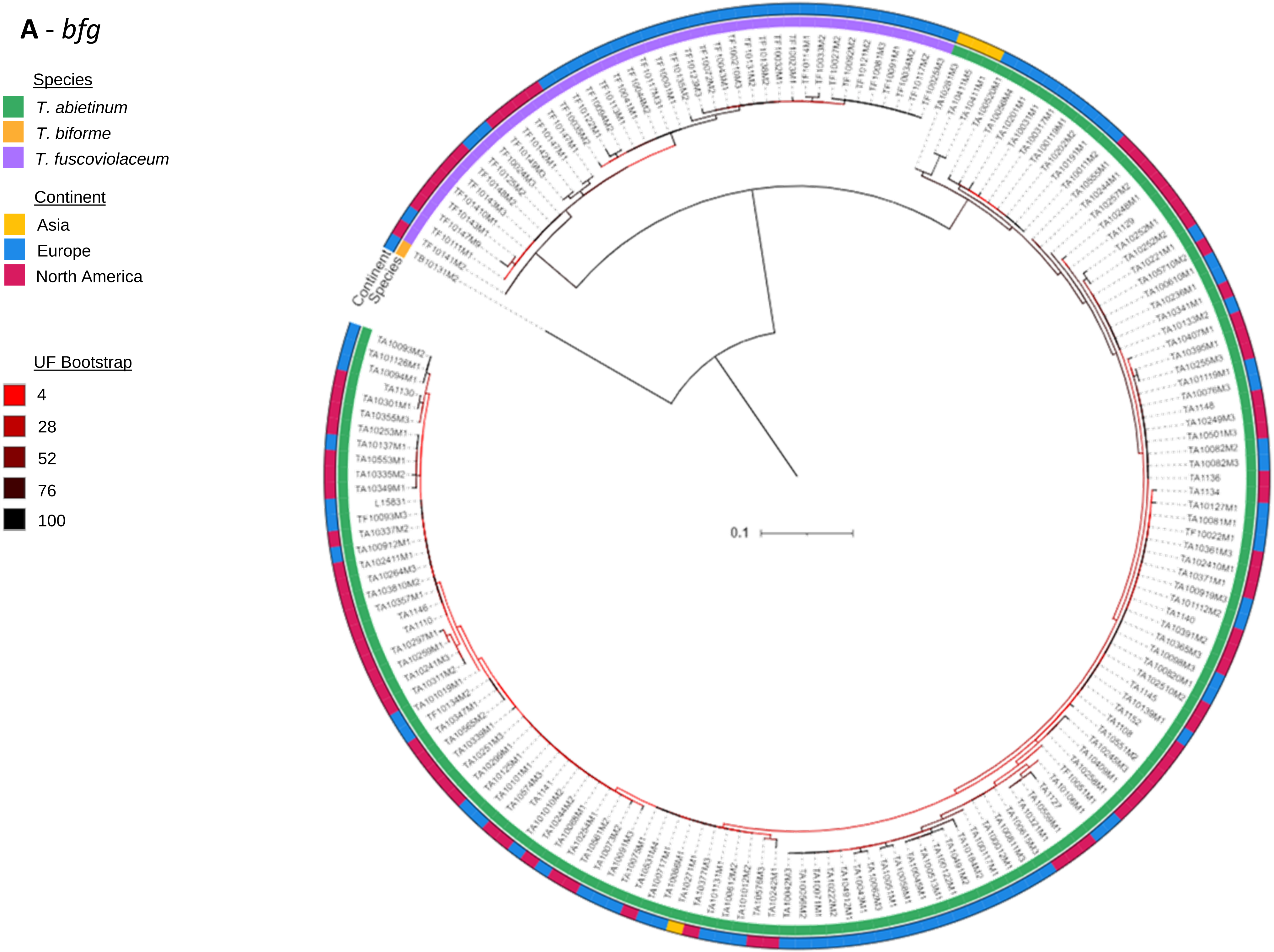
A - *bfg*
Species
T. abietinum
T. biforme
T. fuscoviolaceum
Continent
Asia
Europe
North America
UF Bootstrap
4
28
52
76
100
Continent
Species
0.1

**B** - *GLGEN*

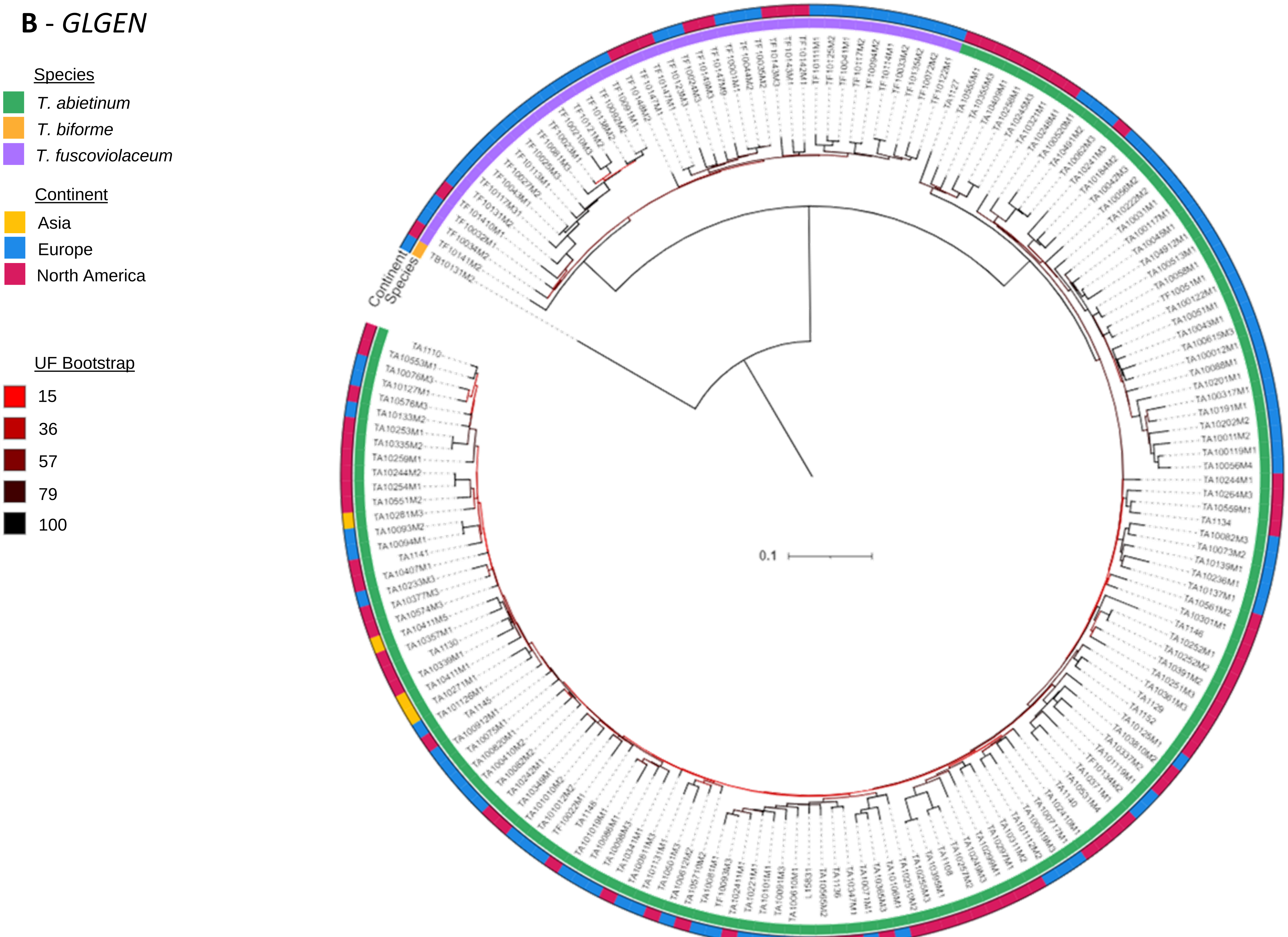
Species
T. abietinum
T. biforme
T. fuscoviolaceum
Continent
Asia
Europe
North America
UF Bootstrap
15
36
57
79
100
Continent
Species
0.1

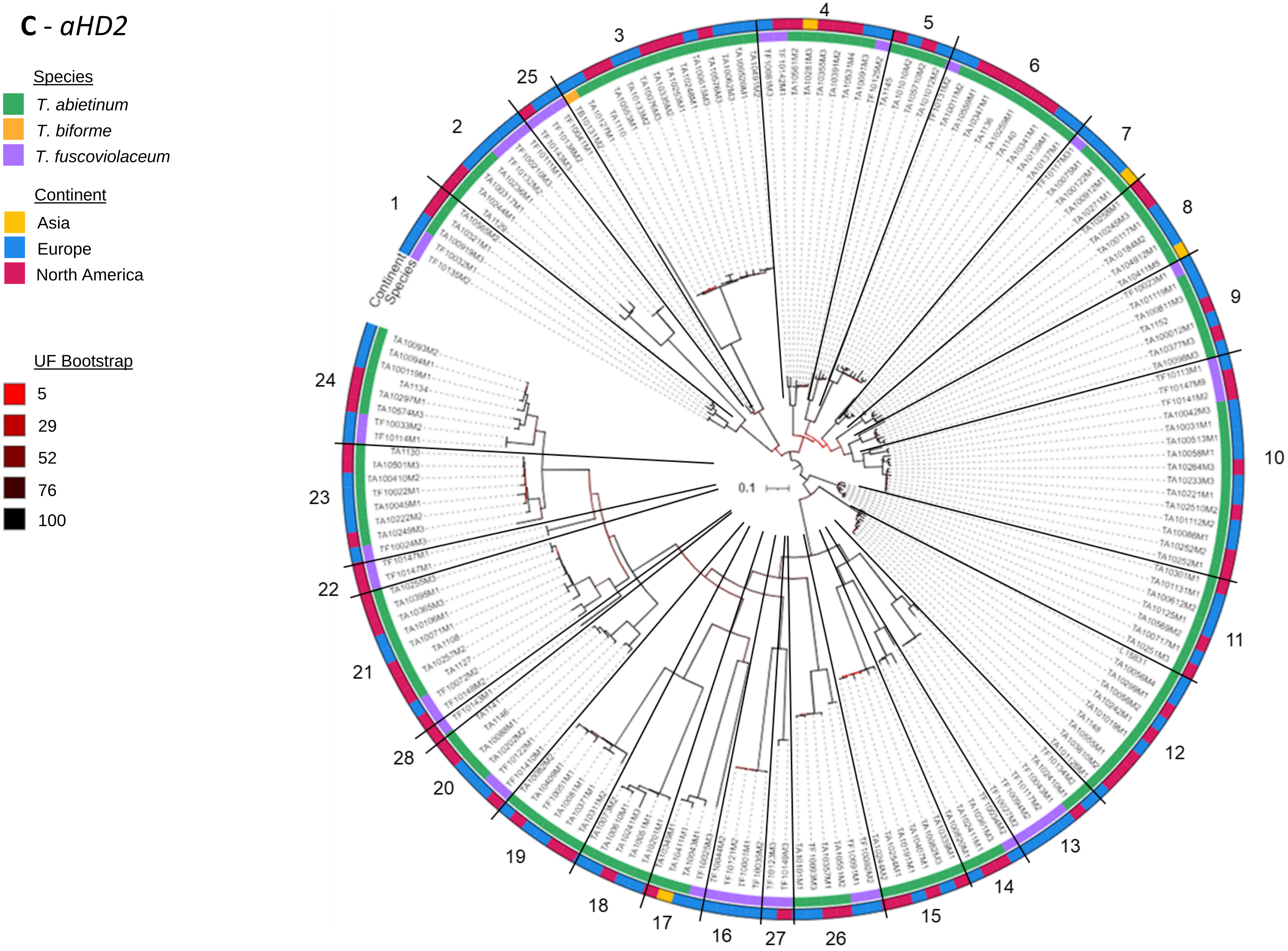
C - aHD2
Species
T. abietinum
T. biforme
T. fuscoviolaceum
Continent
Asia
Europe
North America
UF Bootstrap
5
29
52
76
100
Continent
Species
0.1

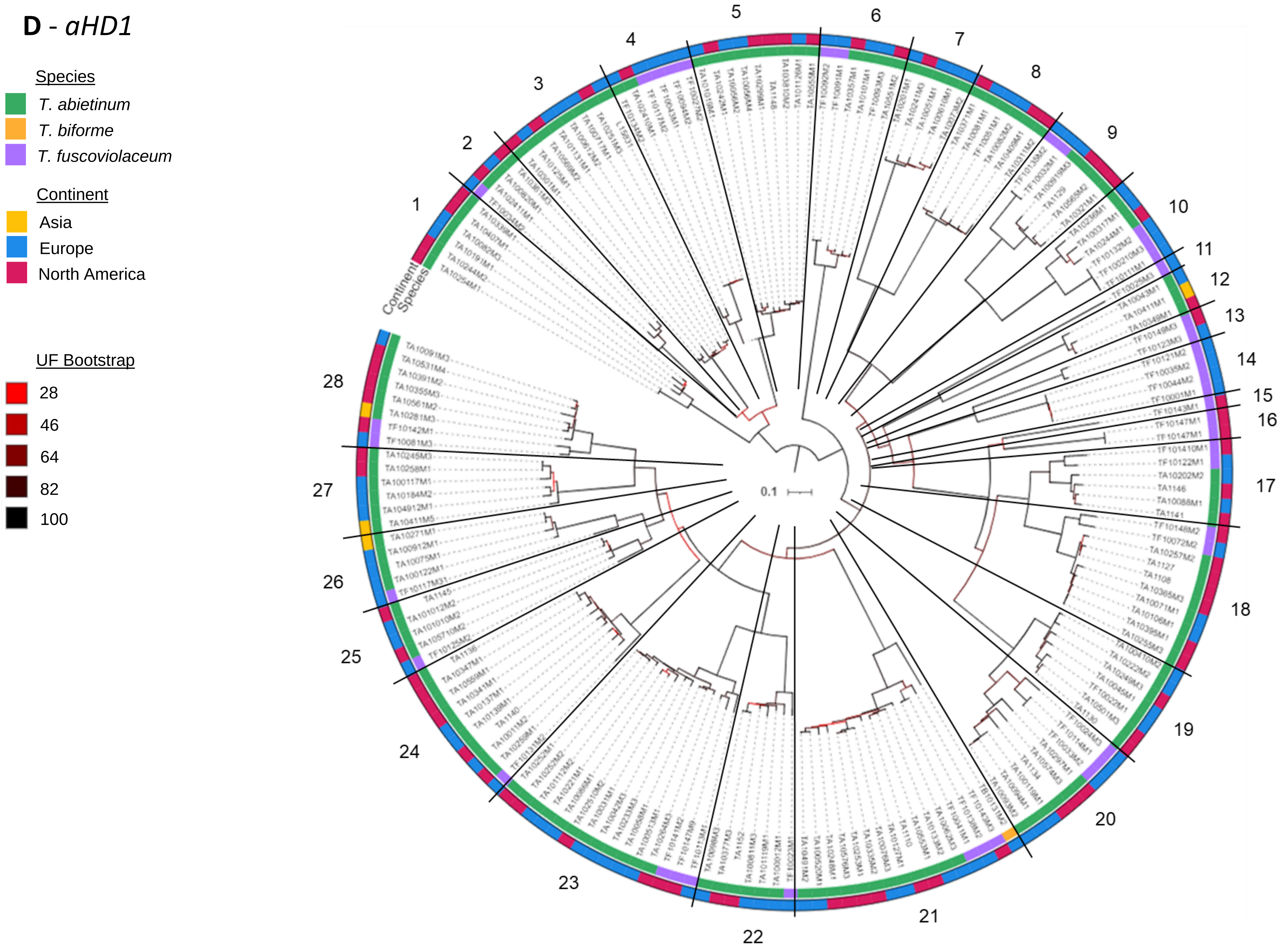
D - aHD1
Species
T. abietinum
T. biforme
T. fuscoviolaceum
Continent
Asia
Europe
North America
UF Bootstrap
28
46
64
82
100
Continent
Species
0.1

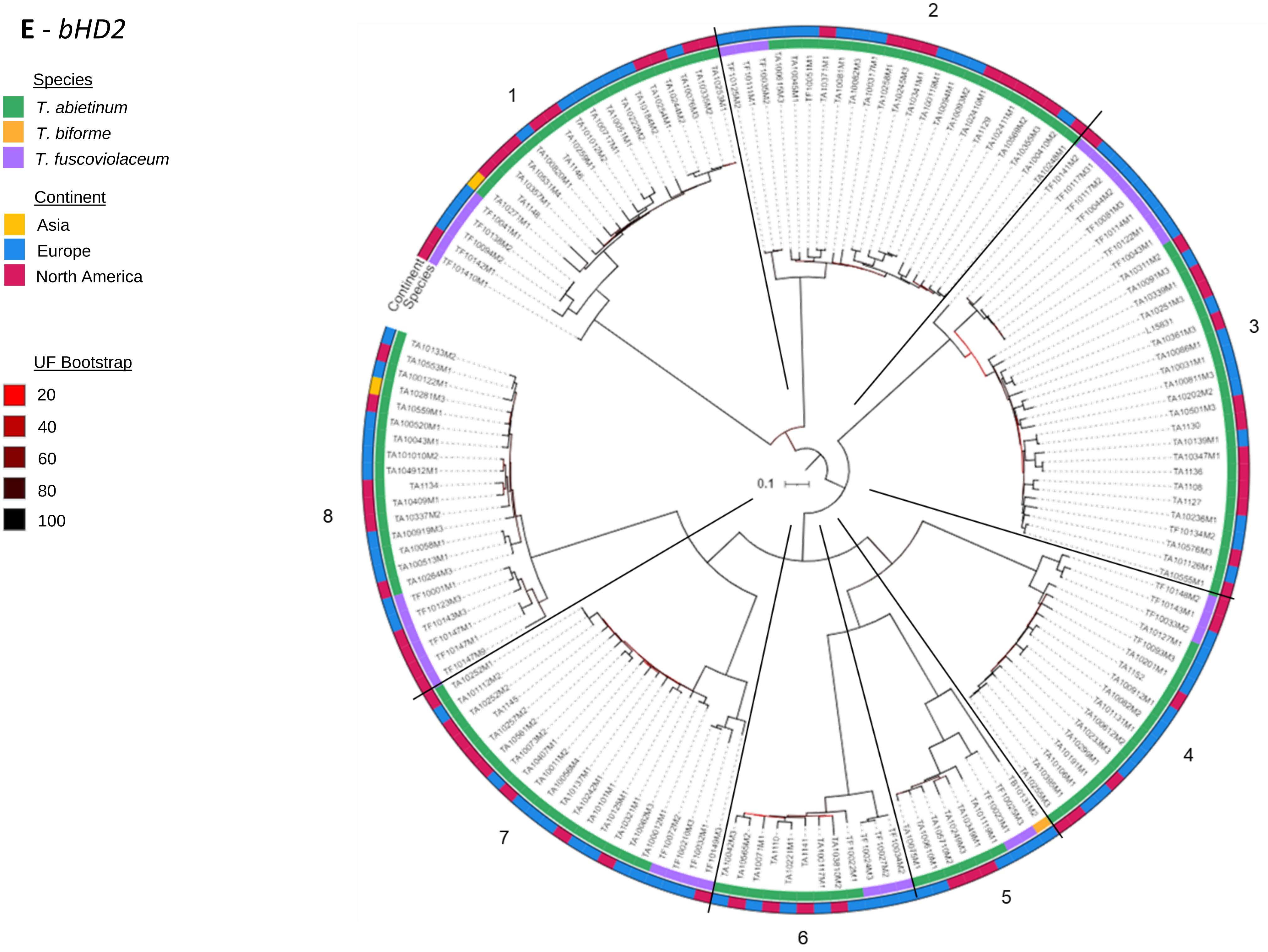

E - bHD2
Species
T. abietinum
T. biforme
T. fuscoviolaceum
Continent
Asia
Europe
North America
UF Bootstrap
20
40
60
80
100
Continent
Species
0.1
1
2
3
4
5
6
7
8

**F** - *bHD1*

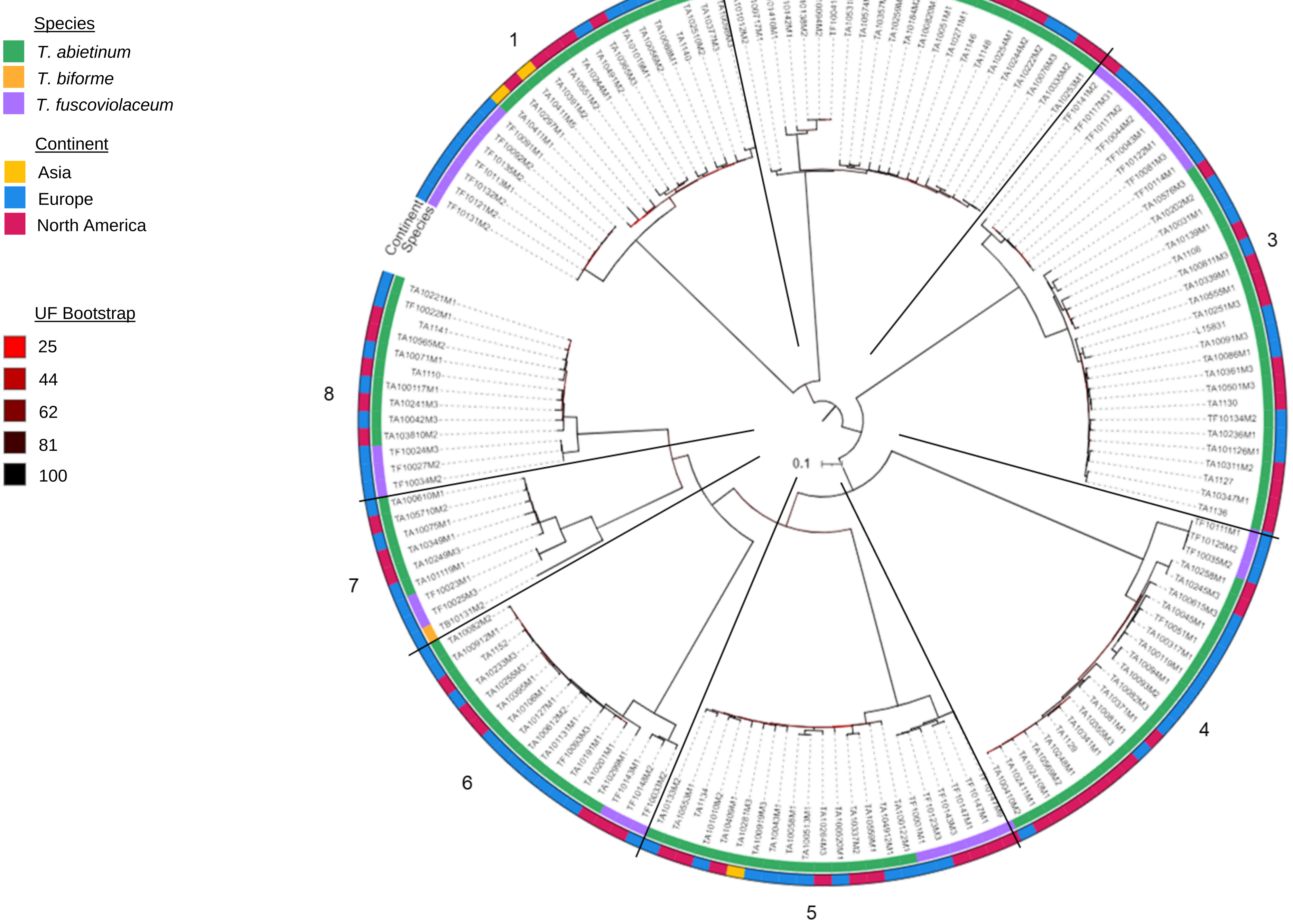

Species
T. abietinum
T. biforme
T. fuscoviolaceum
Continent
Asia
Europe
North America
UF Bootstrap
25
44
62
81
100
Continent
Species
0.1
1
2
3
4
5
6
7
8

G - *MIP1*

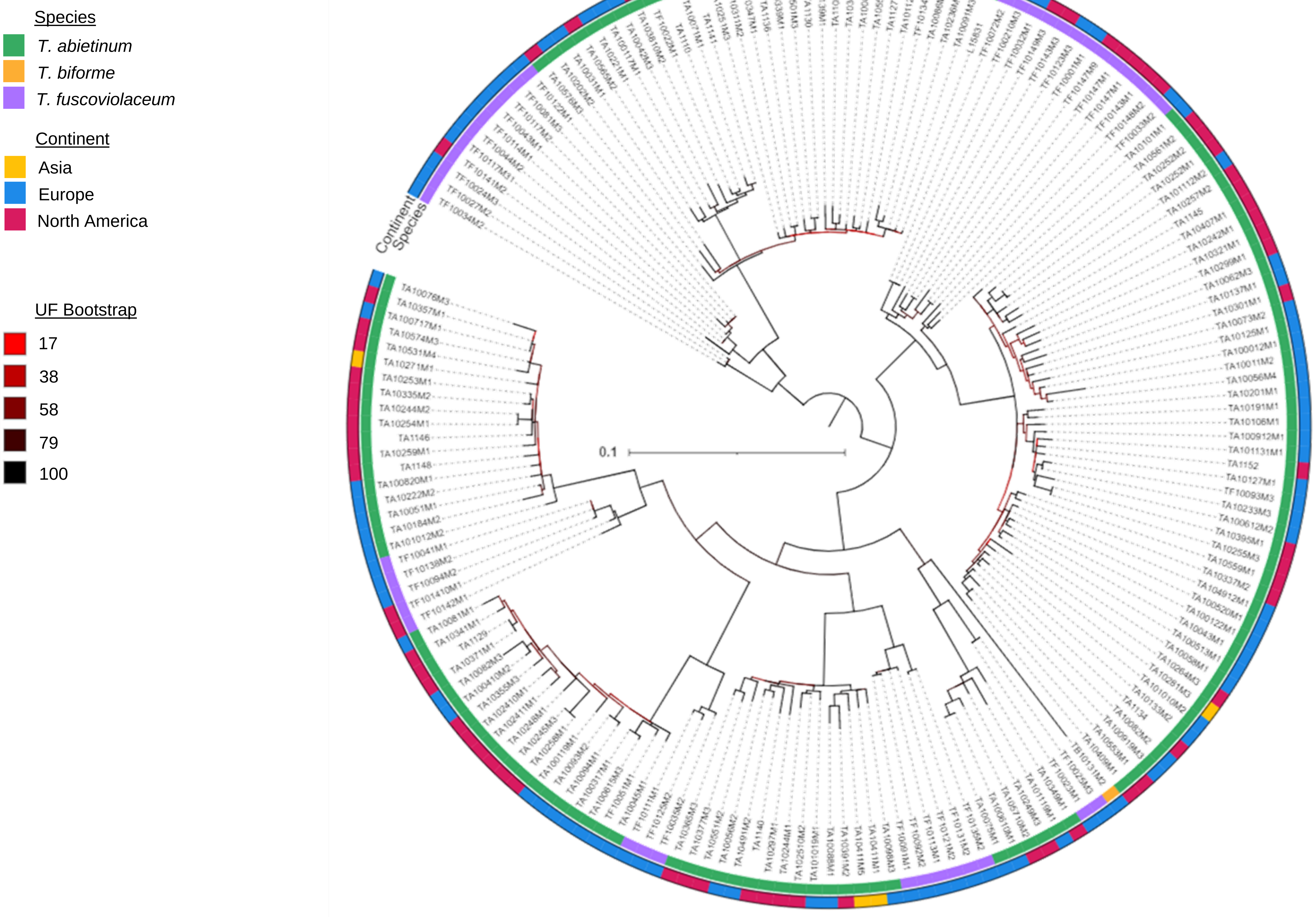
Species
T. abietinum
T. biforme
T. fuscoviolaceum
Continent
Asia
Europe
North America
UF Bootstrap
17
38
58
79
100
0.1
Continent
Species

**H** - *PAK*

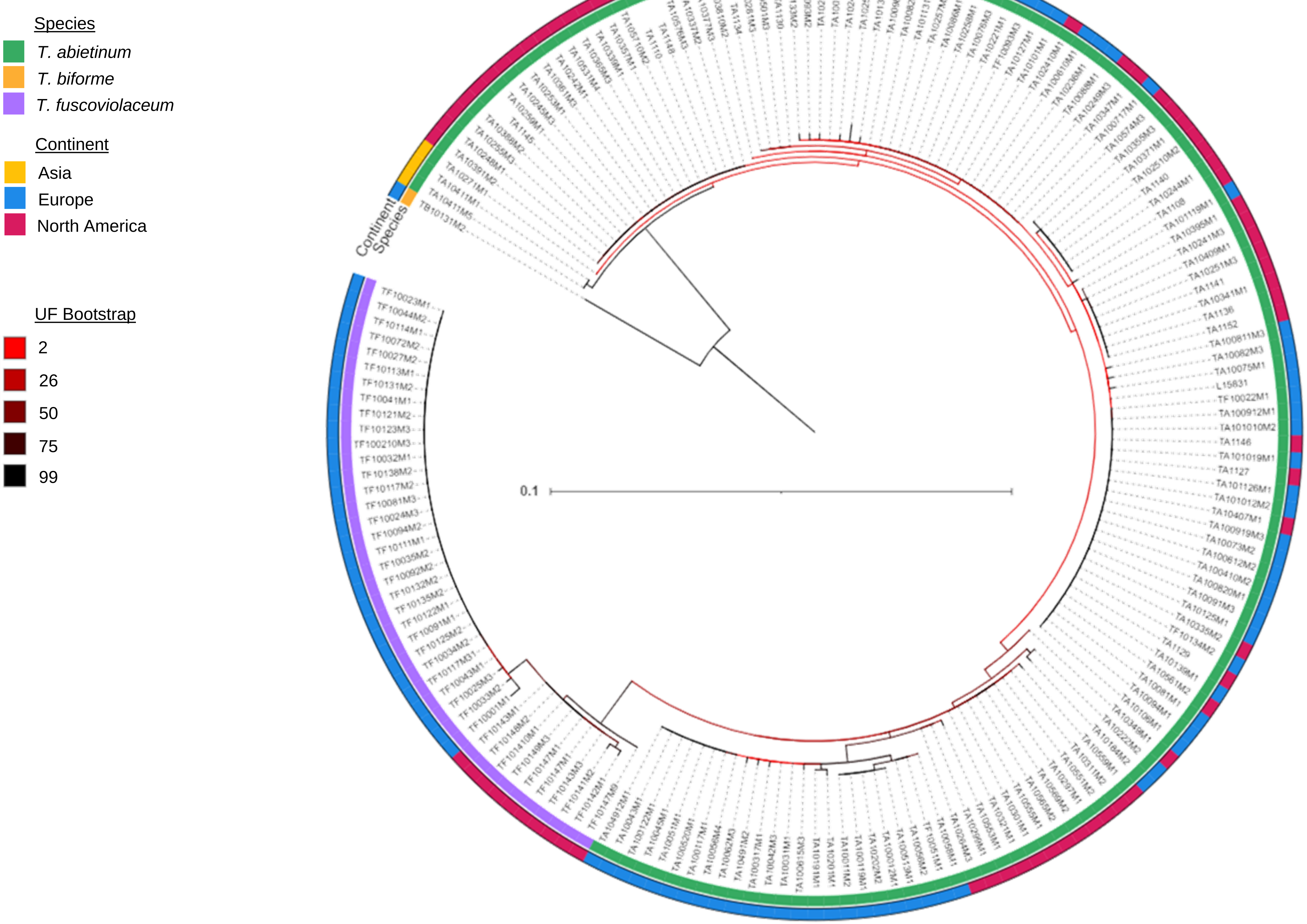
Species
T. abietinum
T. biforme
T. fuscoviolaceum
Continent
Asia
Europe
North America
UF Bootstrap
2
26
50
75
99
Continent
Species
0.1

**I** - *RSM19*

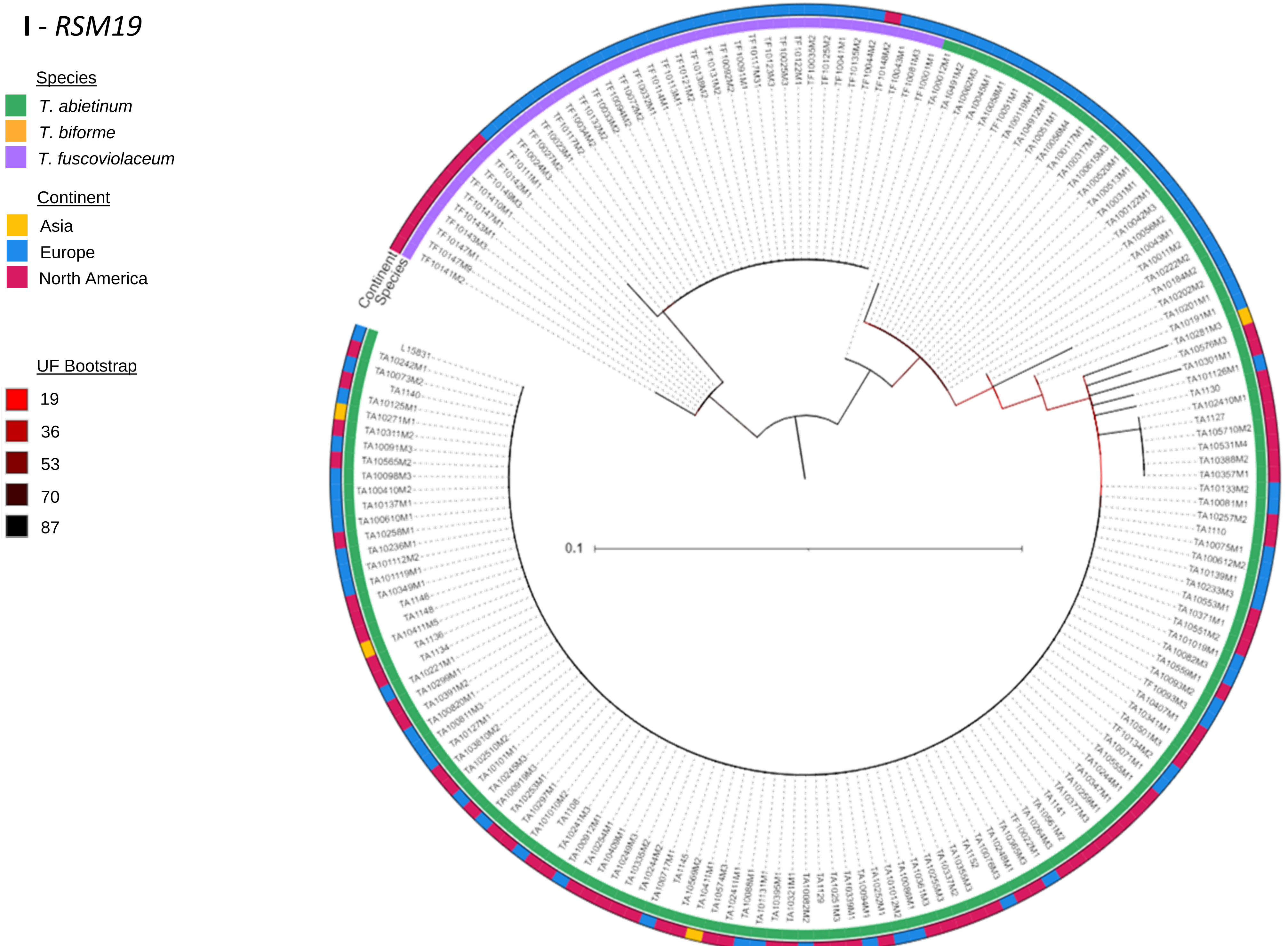

Species
T. abietinum
T. biforme
T. fuscoviolaceum
Continent
Asia
Europe
North America
UF Bootstrap
19
36
53
70
87
Continent
Species
0.1

**J** - *DML1*

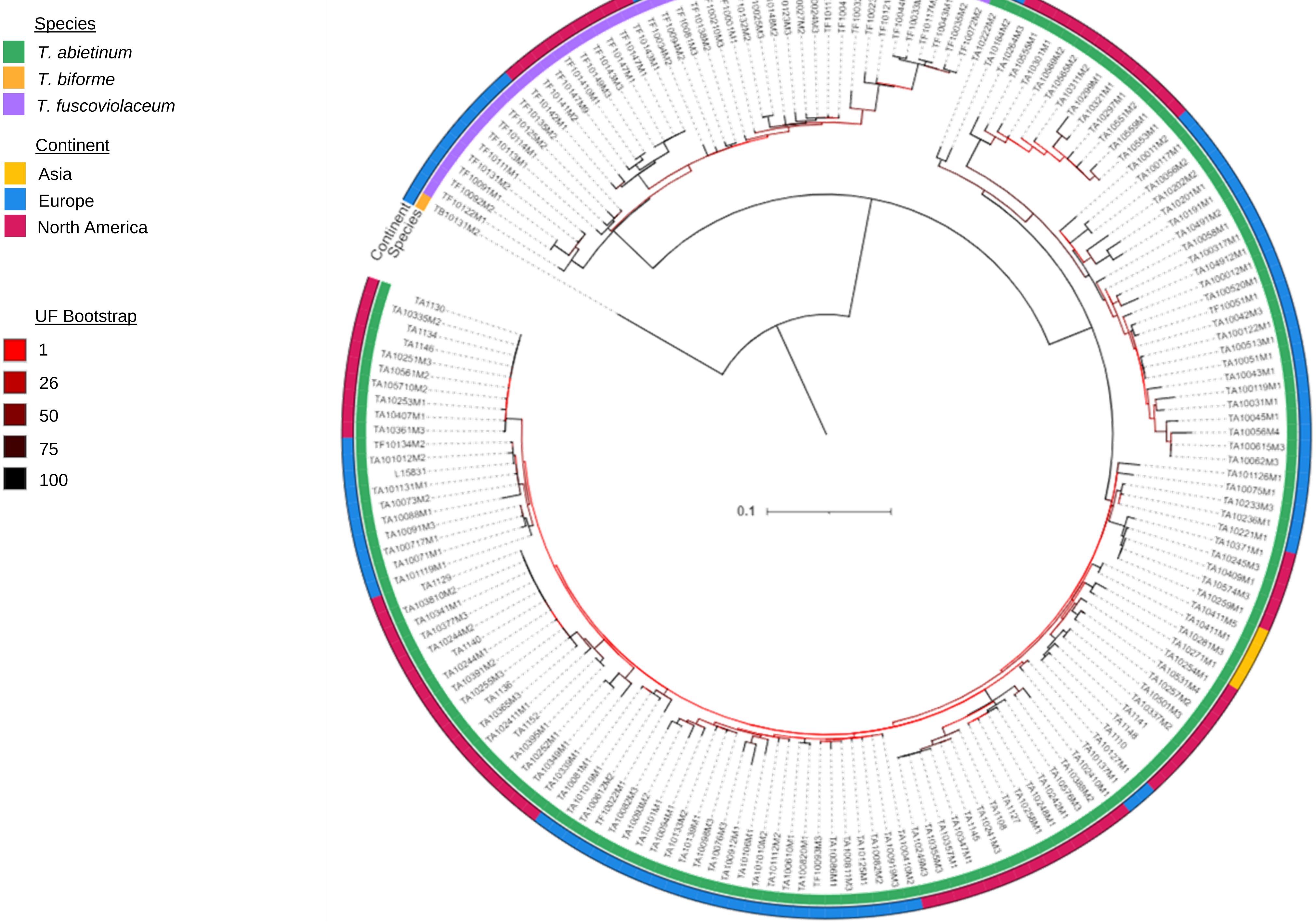
Species
T. abietinum
T. biforme
T. fuscoviolaceum
Continent
Asia
Europe
North America
UF Bootstrap
1
26
50
75
100
Continent
Species
0.1

**K** - *RIC1*

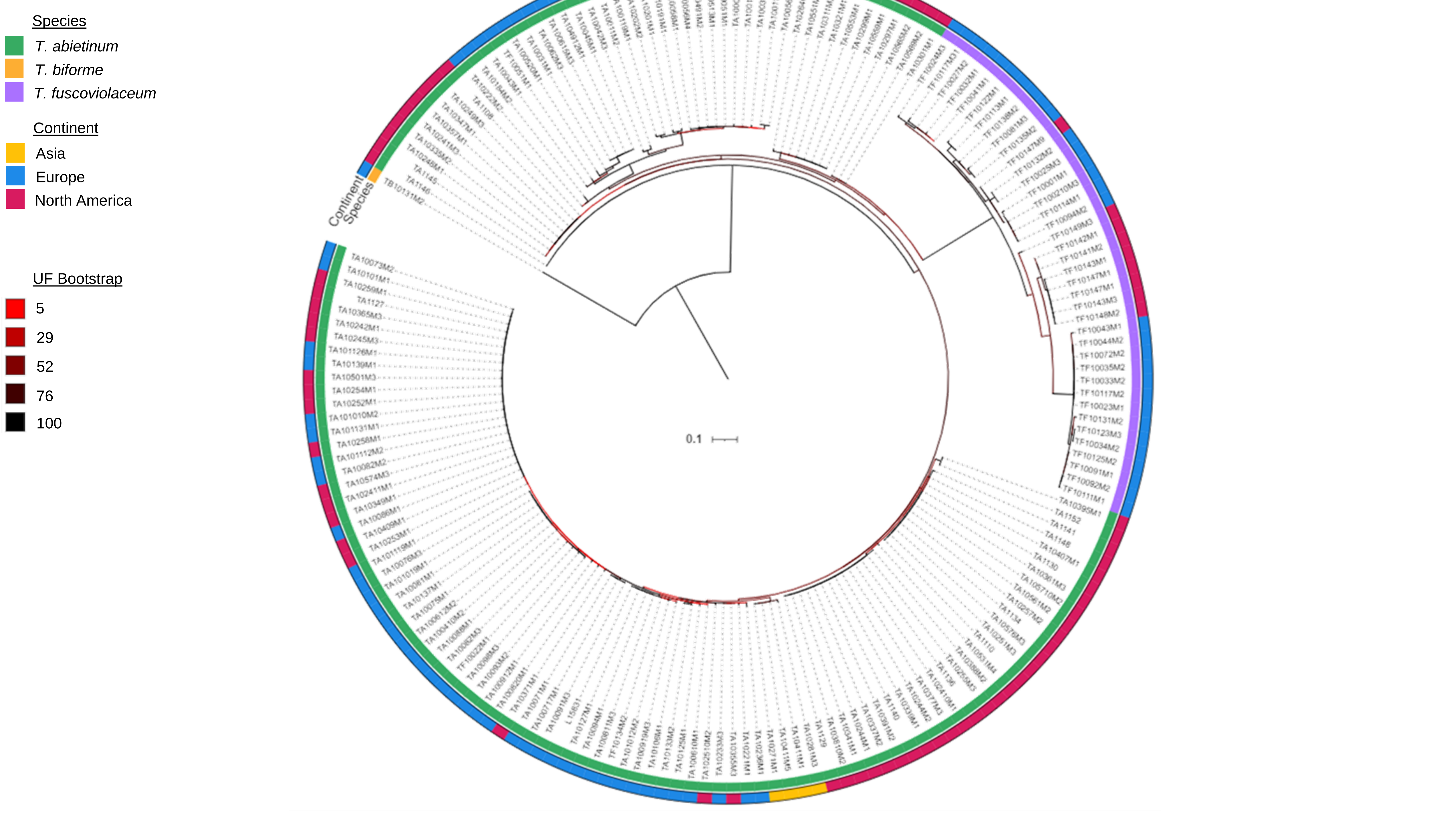

Species
T. abietinum
T. biforme
T. fuscoviolaceum
Continent
Asia
Europe
North America
UF Bootstrap
5
29
52
76
100
Continent
Species
0.1

**L** - *STE3.1*

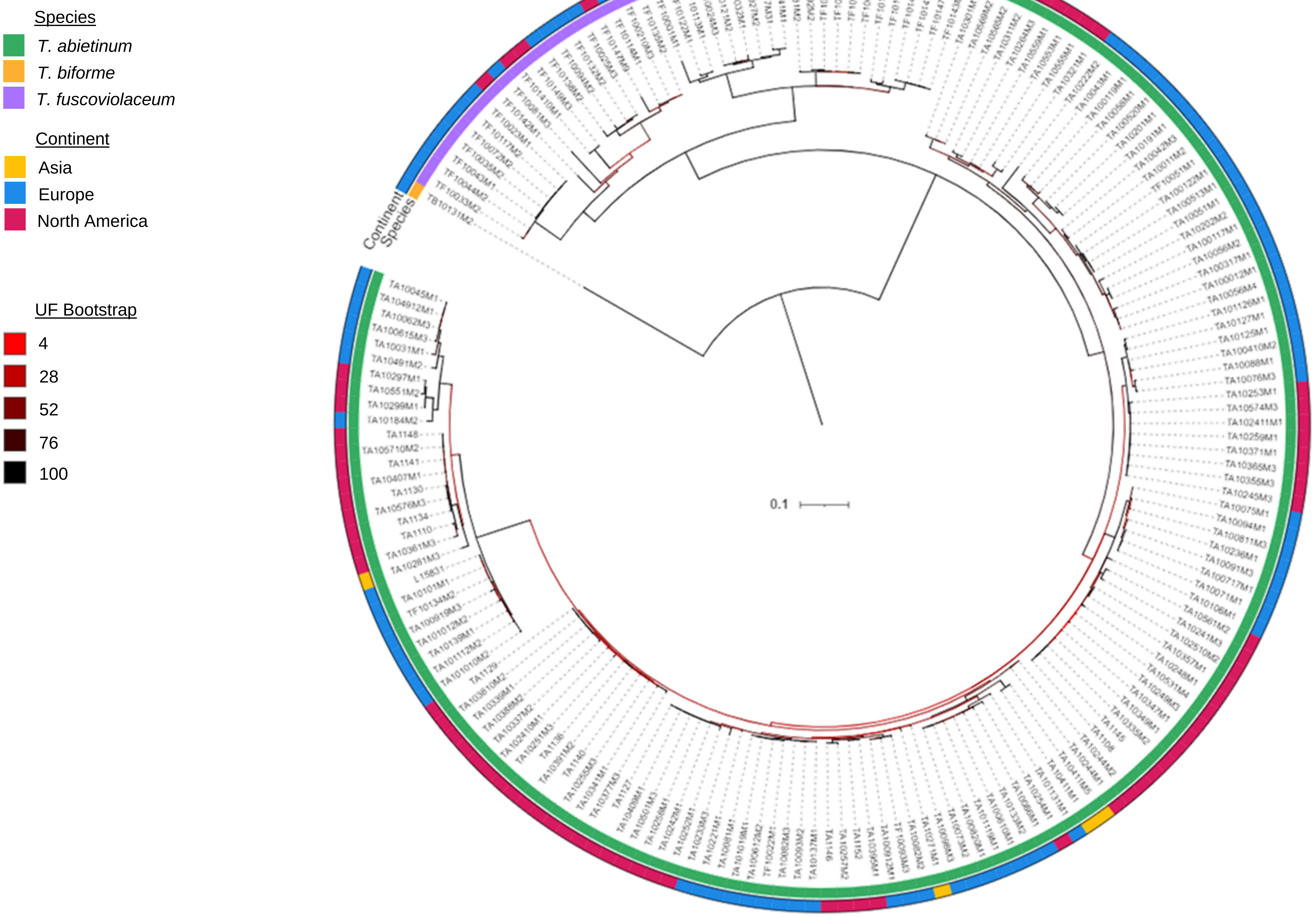
Species
T. abietinum
T. biforme
T. fuscoviolaceum
Continent
Asia
Europe
North America
UF Bootstrap
4
28
52
76
100
Continent
Species
0.1

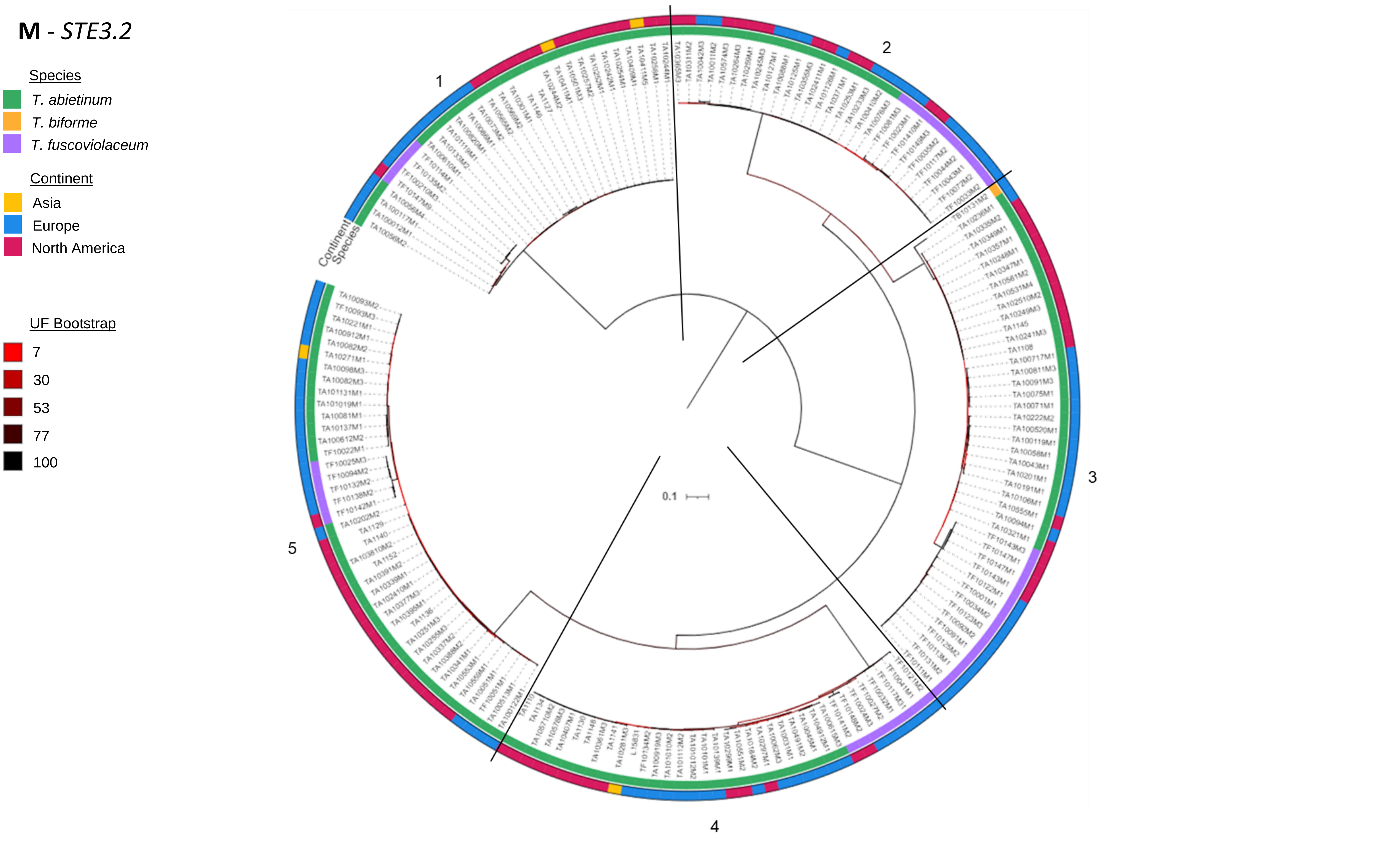
M - STE3.2
Species
T. abietinum
T. biforme
T. fuscoviolaceum
Continent
Asia
Europe
North America
UF Bootstrap
7
30
53
77
100
Continent
Species
1
2
3
4
5
0.1

**N** - *STE3.3*

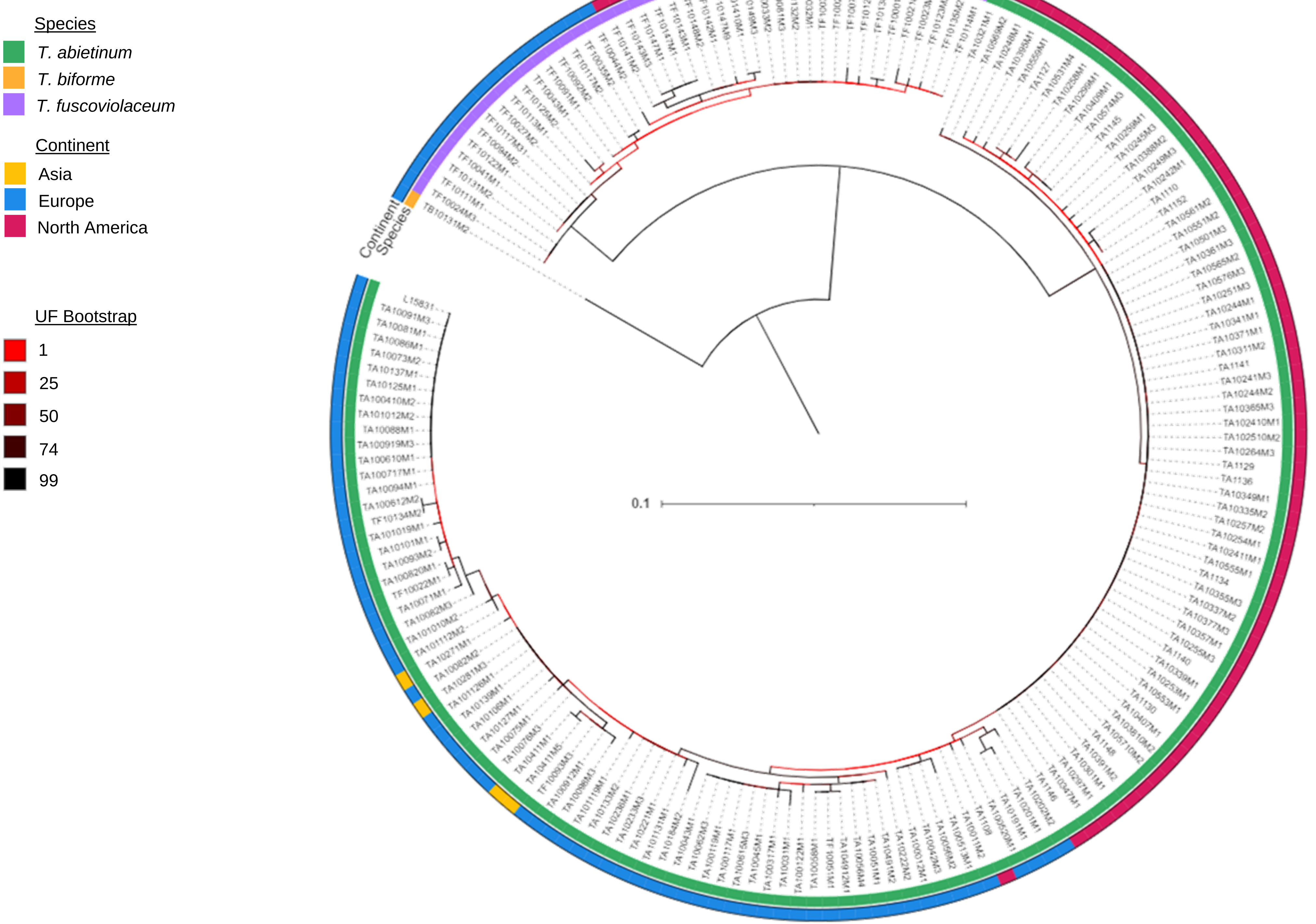
Species
T. abietinum
T. biforme
T. fuscoviolaceum
Continent
Asia
Europe
North America
UF Bootstrap
1
25
50
74
99
Continent
Species
0.1

**O** - *STE3.4*

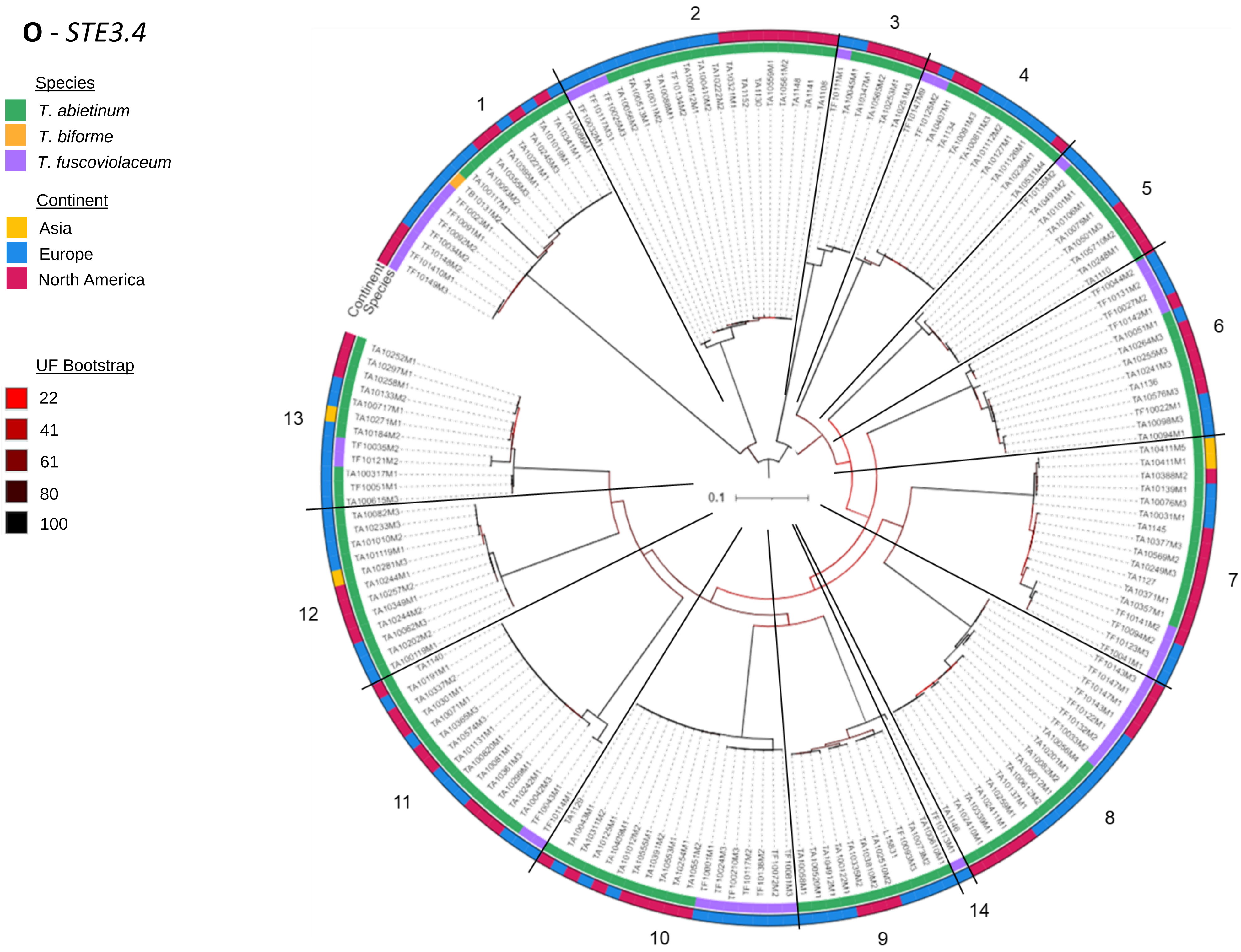
Species
T. abietinum
T. biforme
T. fuscoviolaceum
Continent
Asia
Europe
North America
UF Bootstrap
22
41
61
80
100
Continent
Species
0.1
1
2
3
4
5
6
7
8
9
10
11
12
13
14

**P** - *SNF2*

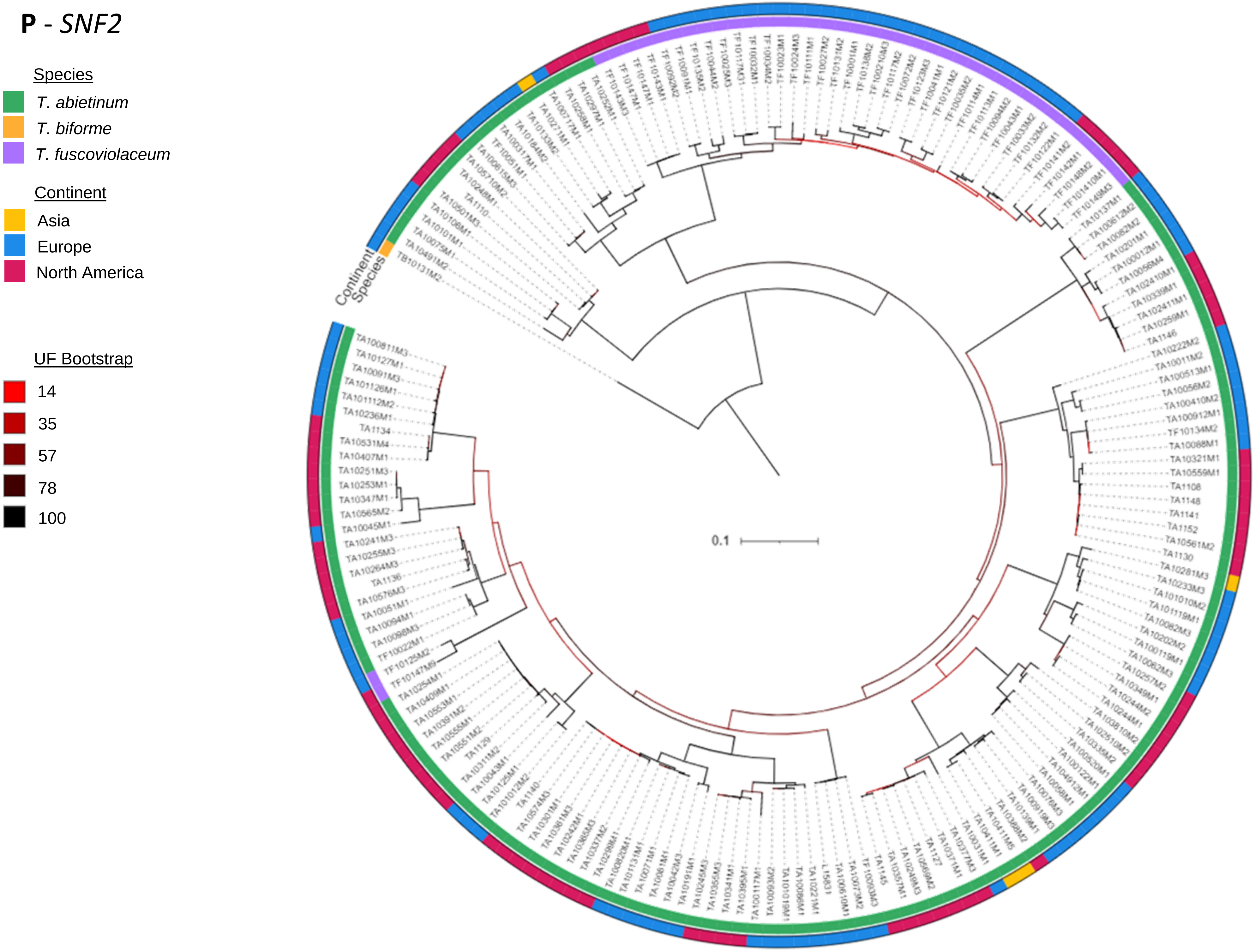
Species
T. abietinum
T. biforme
T. fuscoviolaceum
Continent
Asia
Europe
North America
UF Bootstrap
14
35
57
78
100
Continent
Species
0.1
