## Supplementary figures and images for "Molecular diversity maintained by long-term balancing selection in mating loci defines multiple mating types in fungi"

### Supplementary Figure 1.pdf

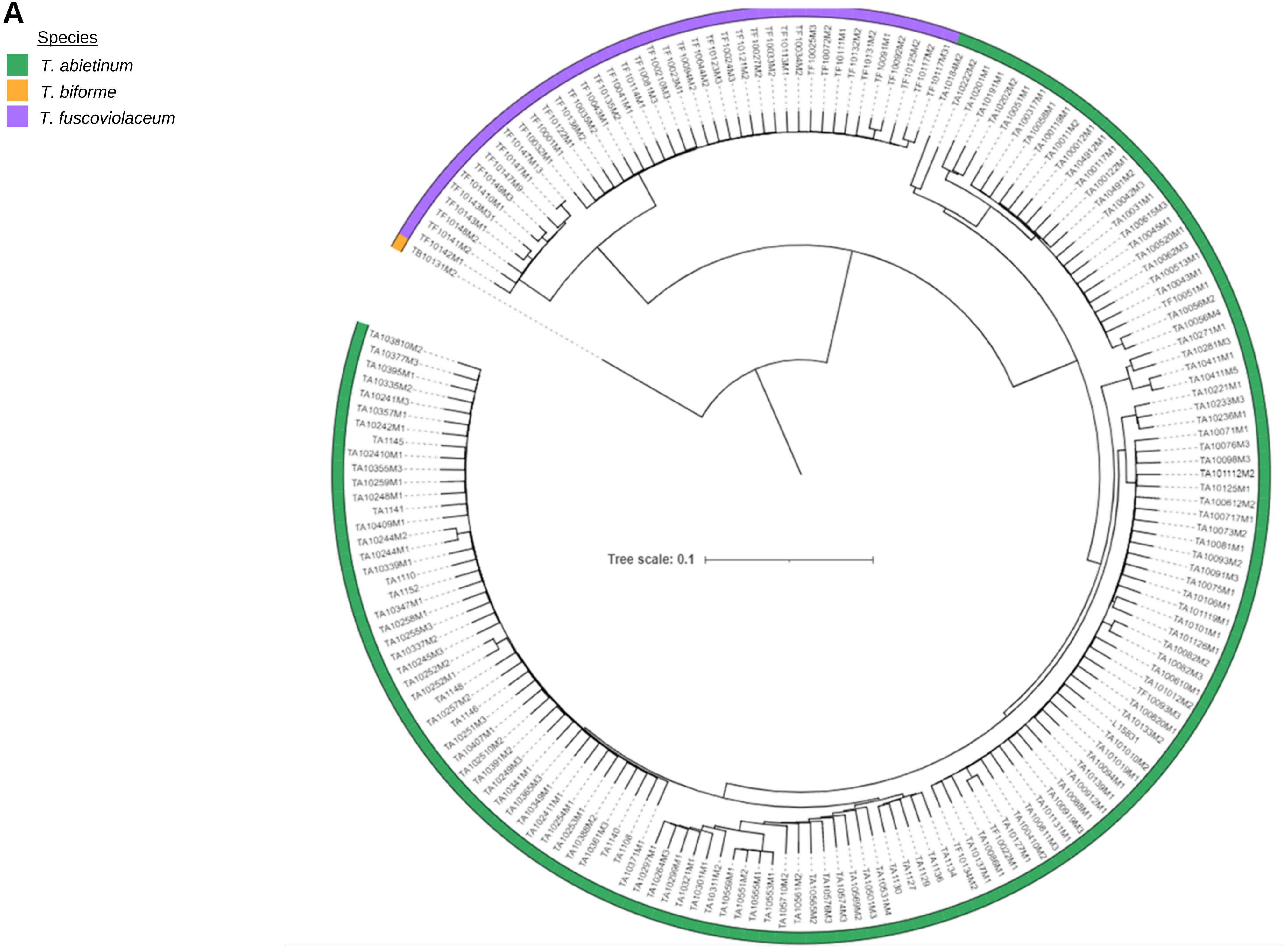

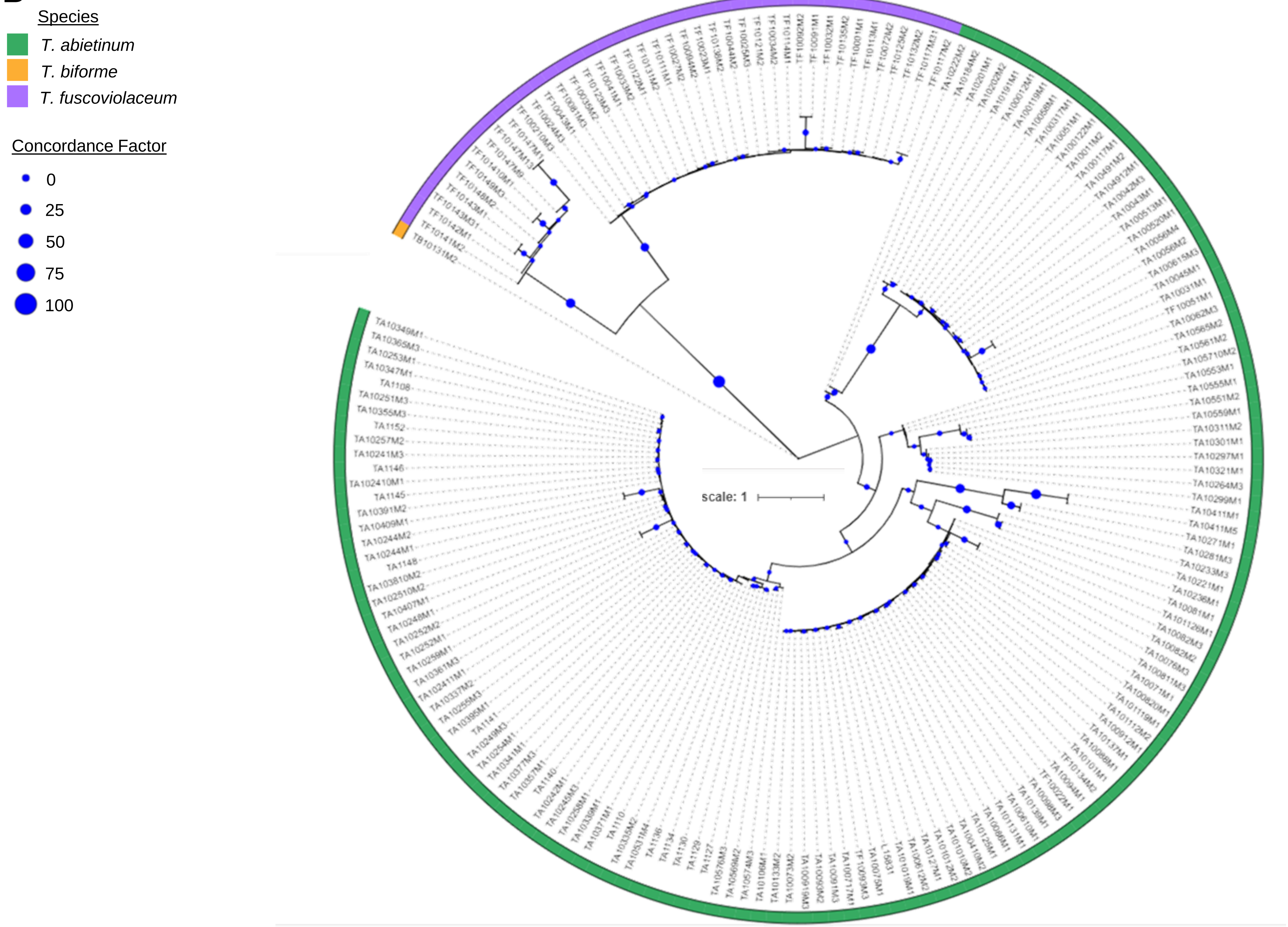

### Supplementary Figure 2.pdf

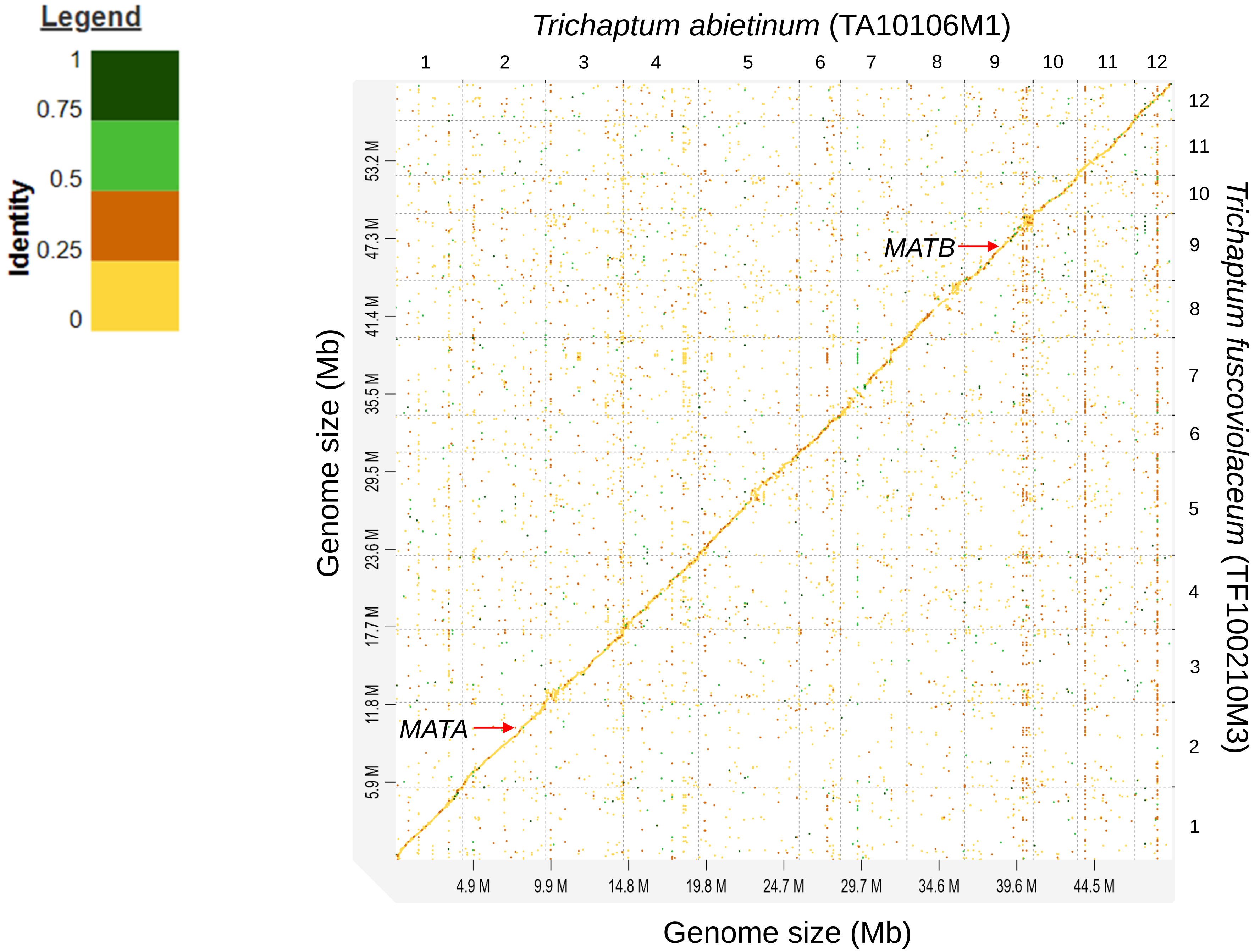

### Supplementary Figure 3.pdf

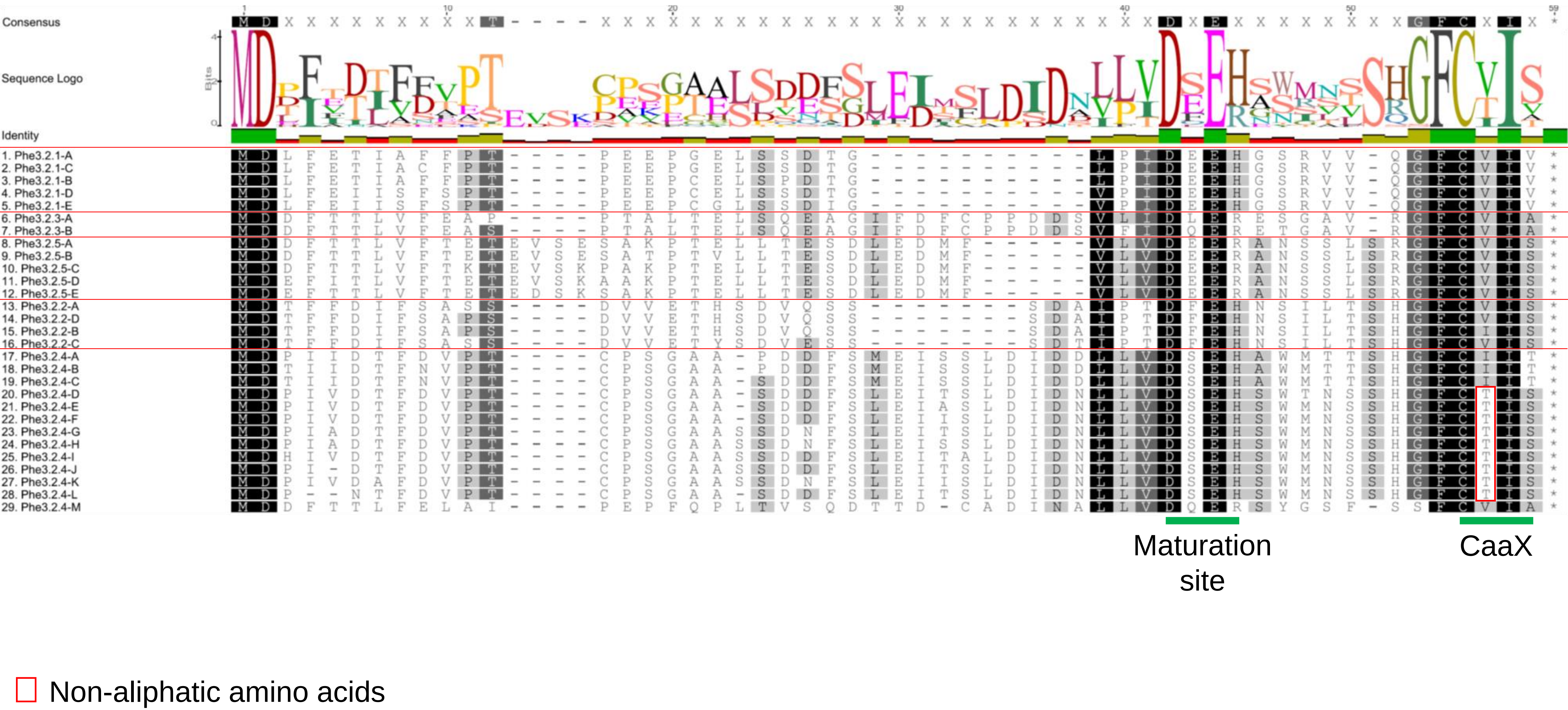

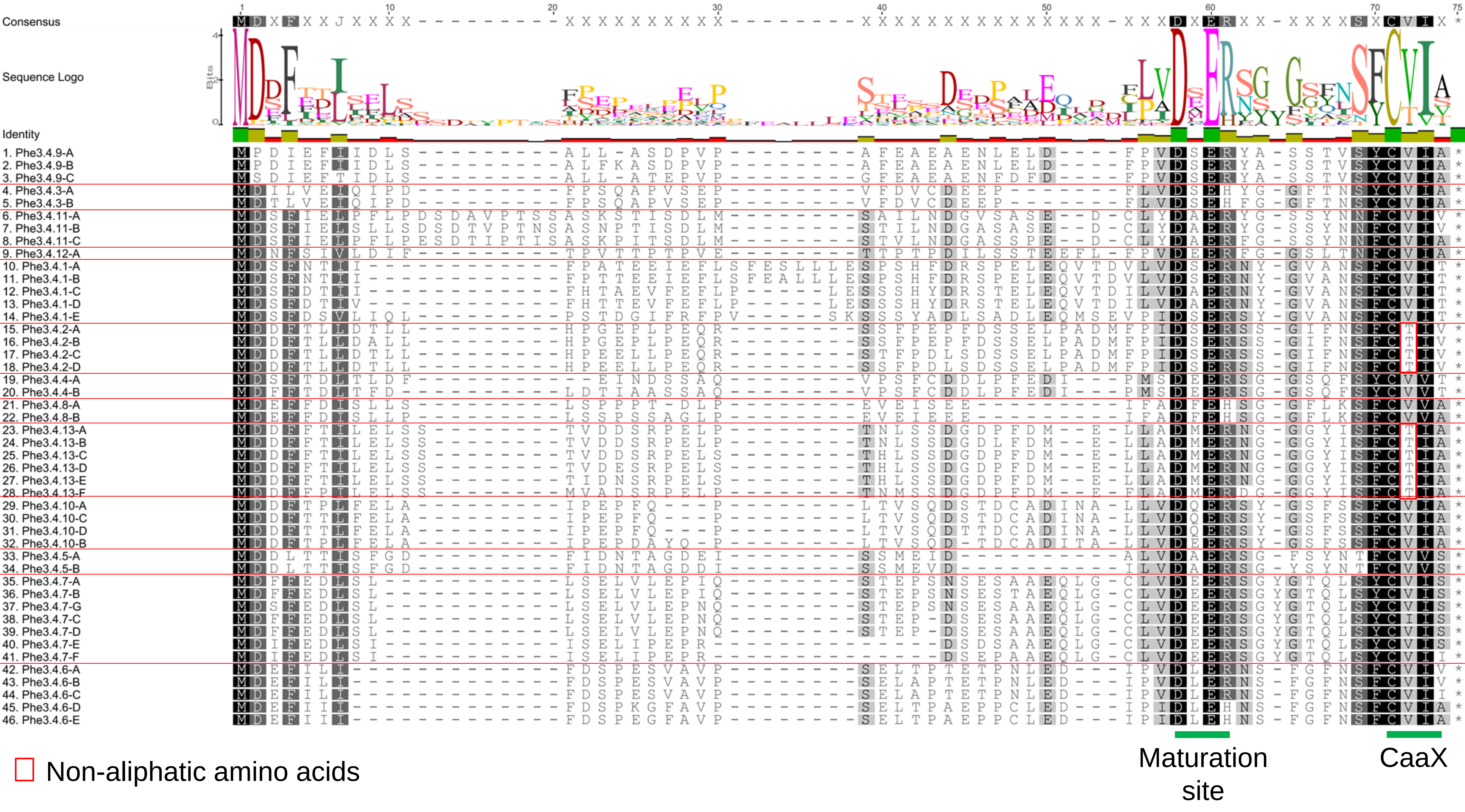

### Supplementary Figure 5.pdf

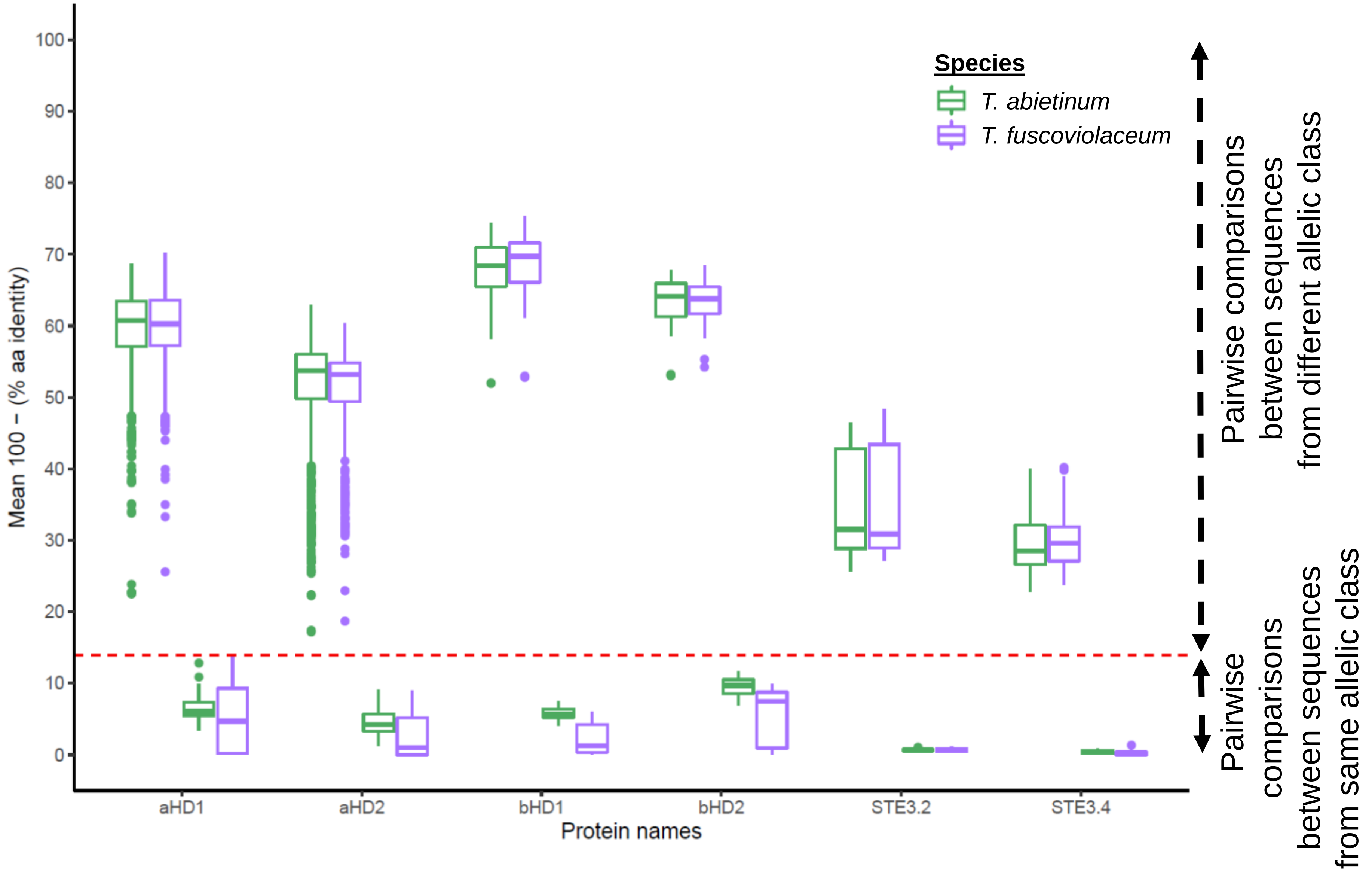

### Supplementary Figure 6.pdf

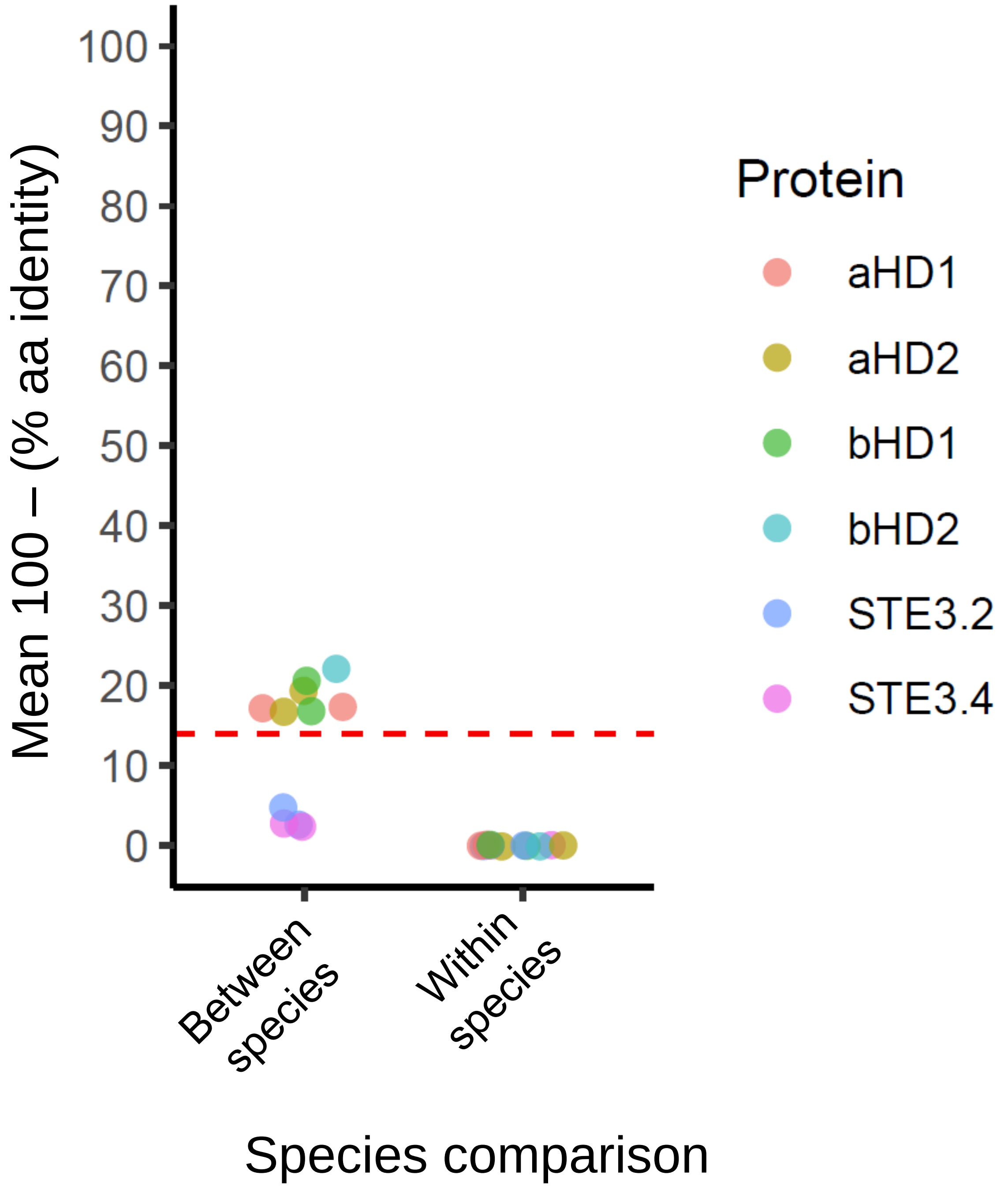

### Supplementary Figure 7.pdf

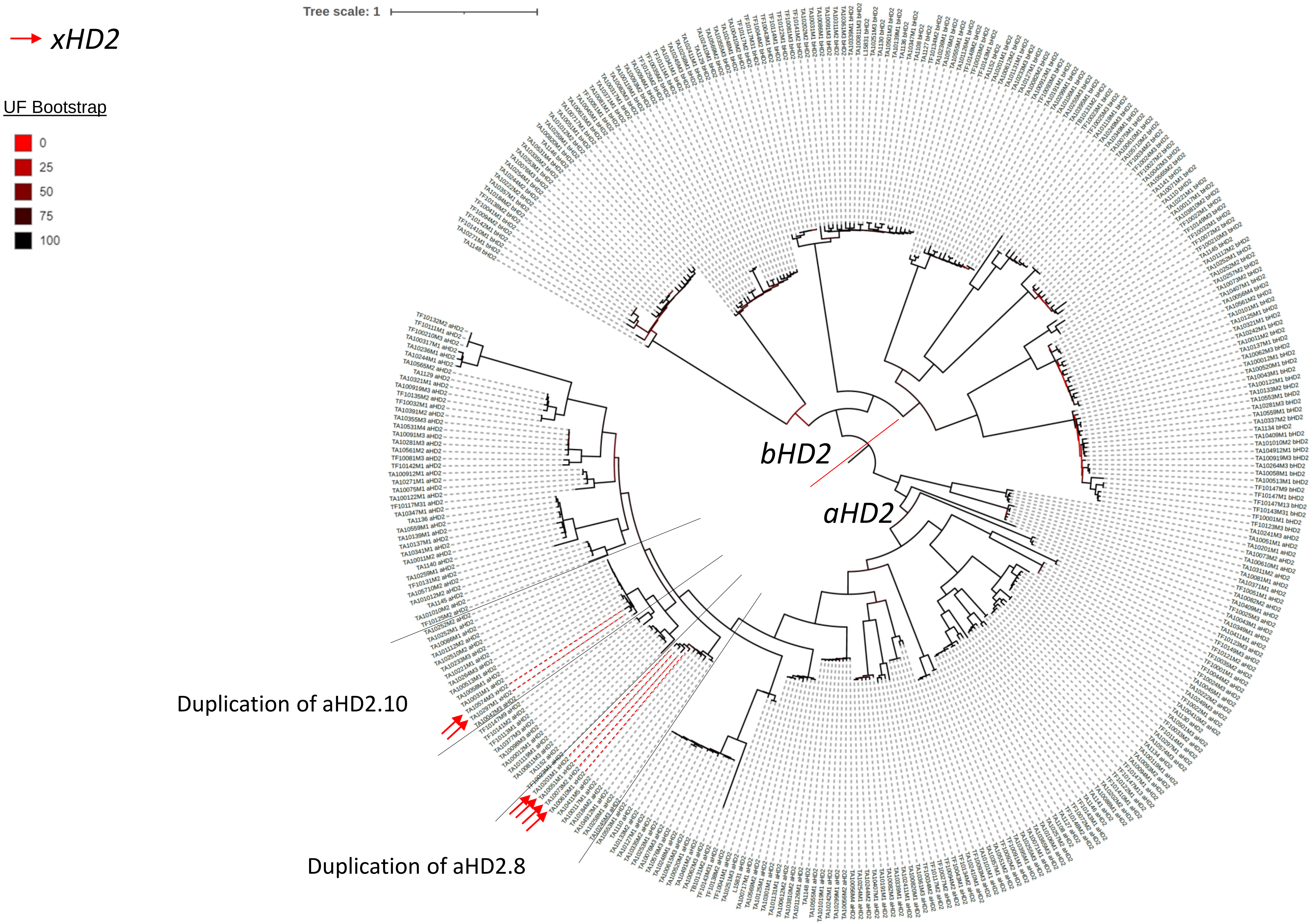

### Supplementary Figure 8.pdf

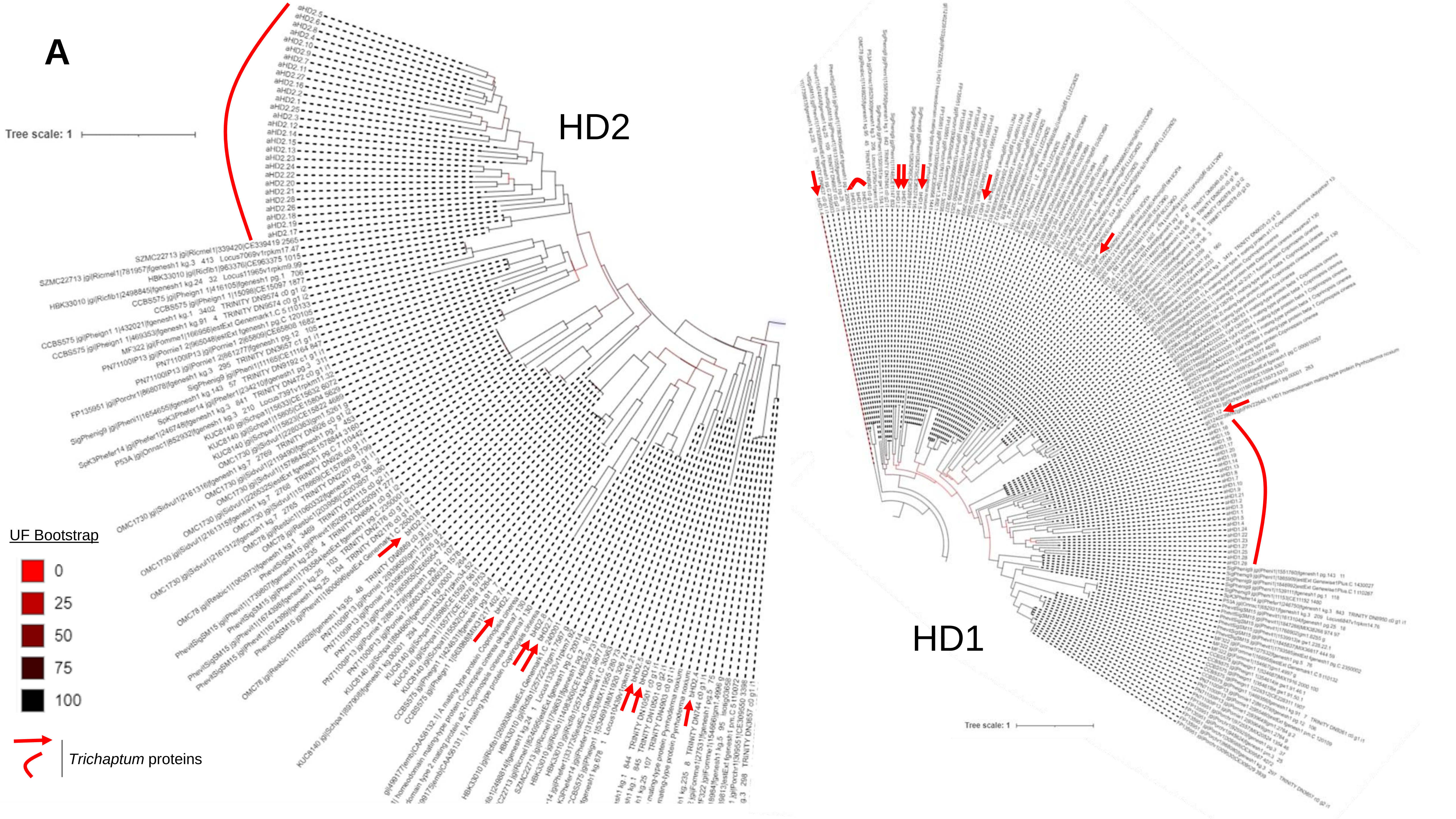

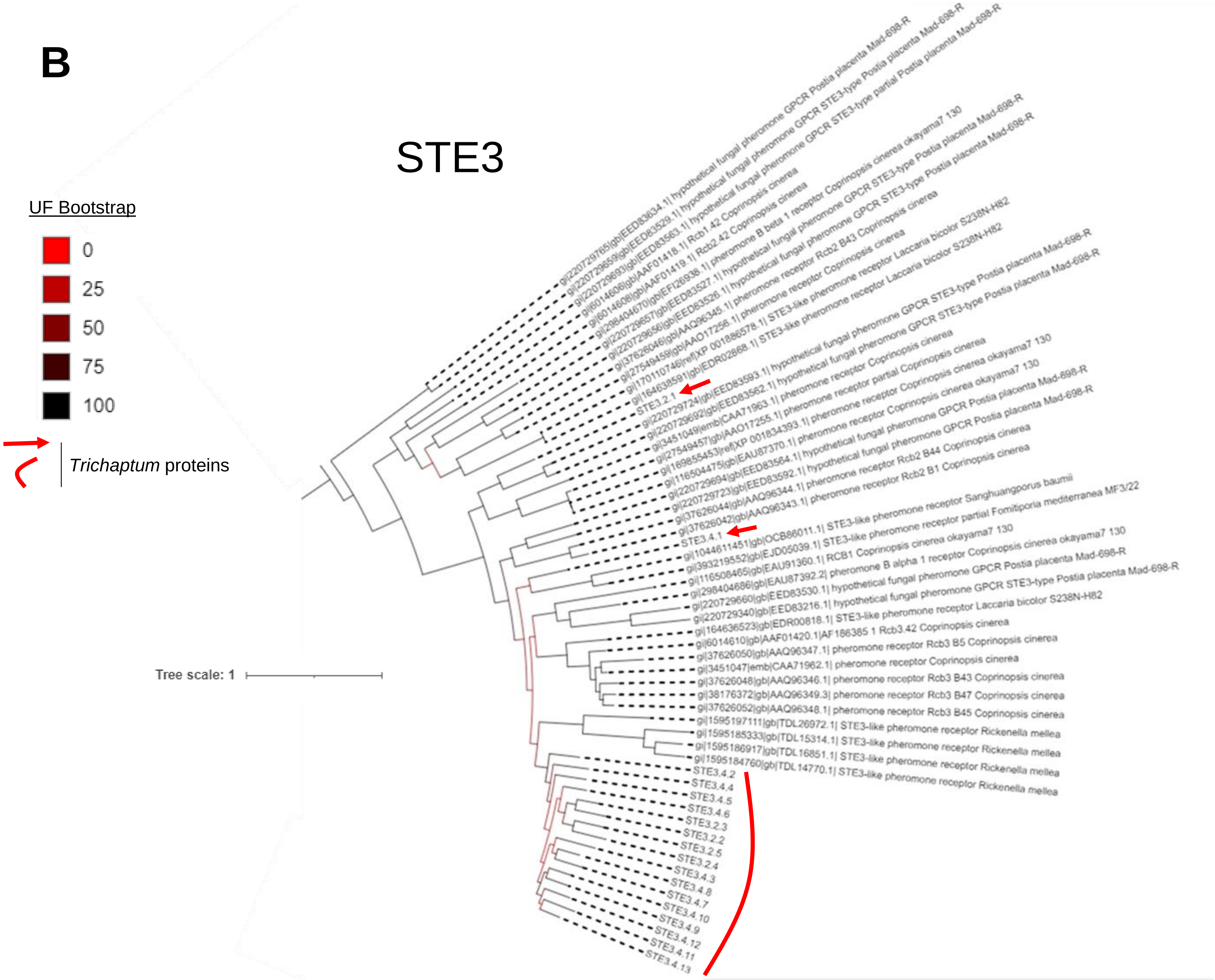

### Supplementary Figure 9.pdf

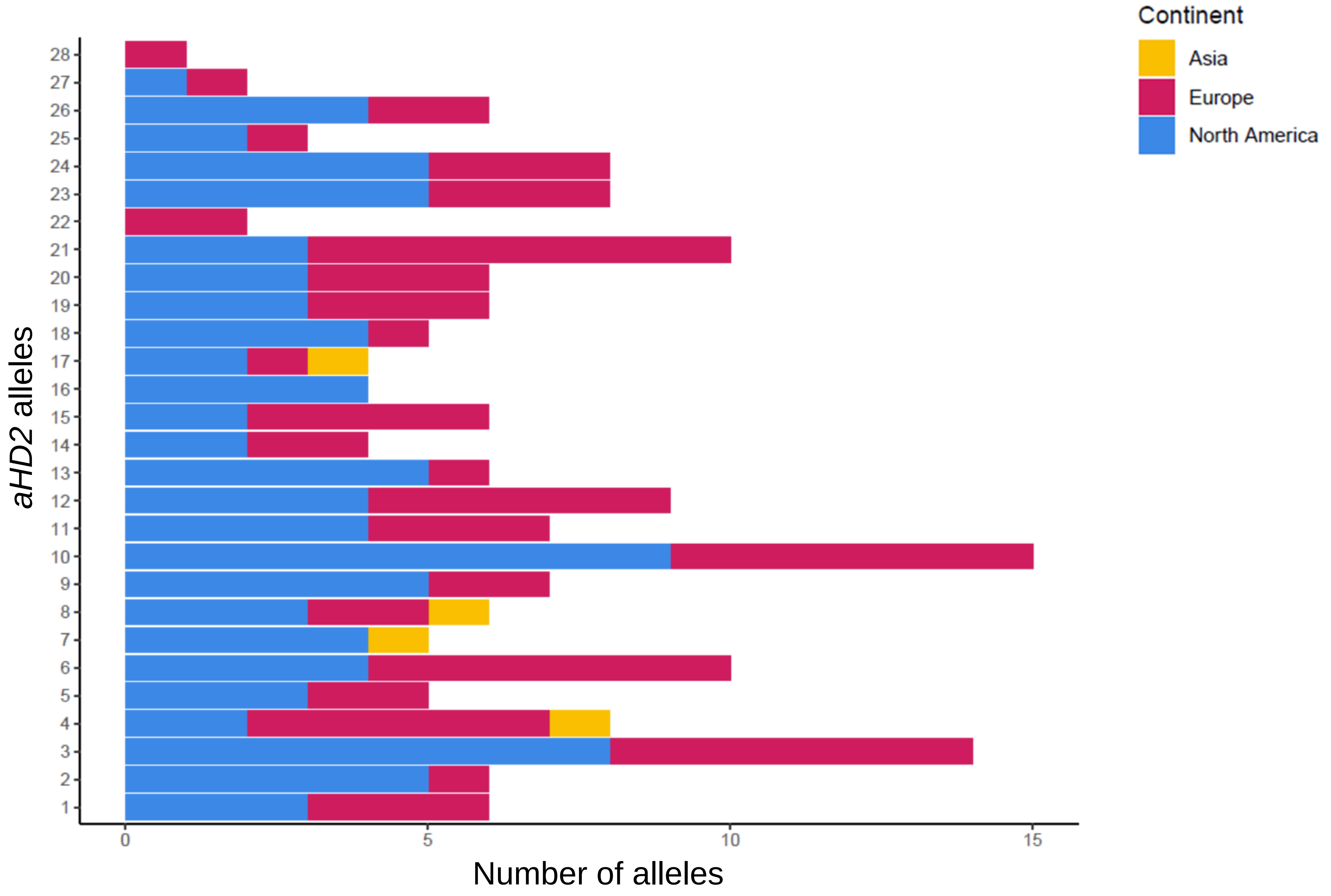

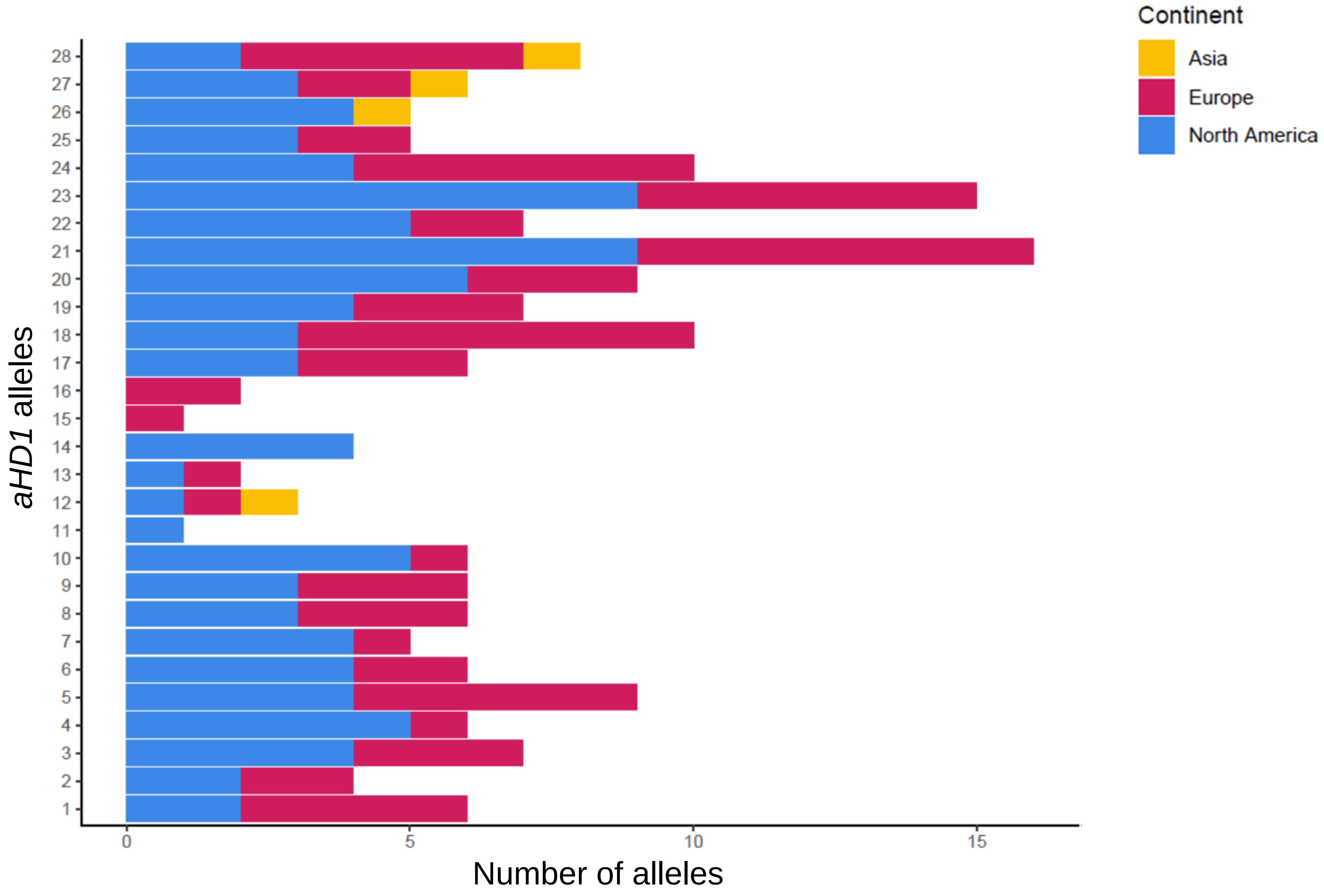

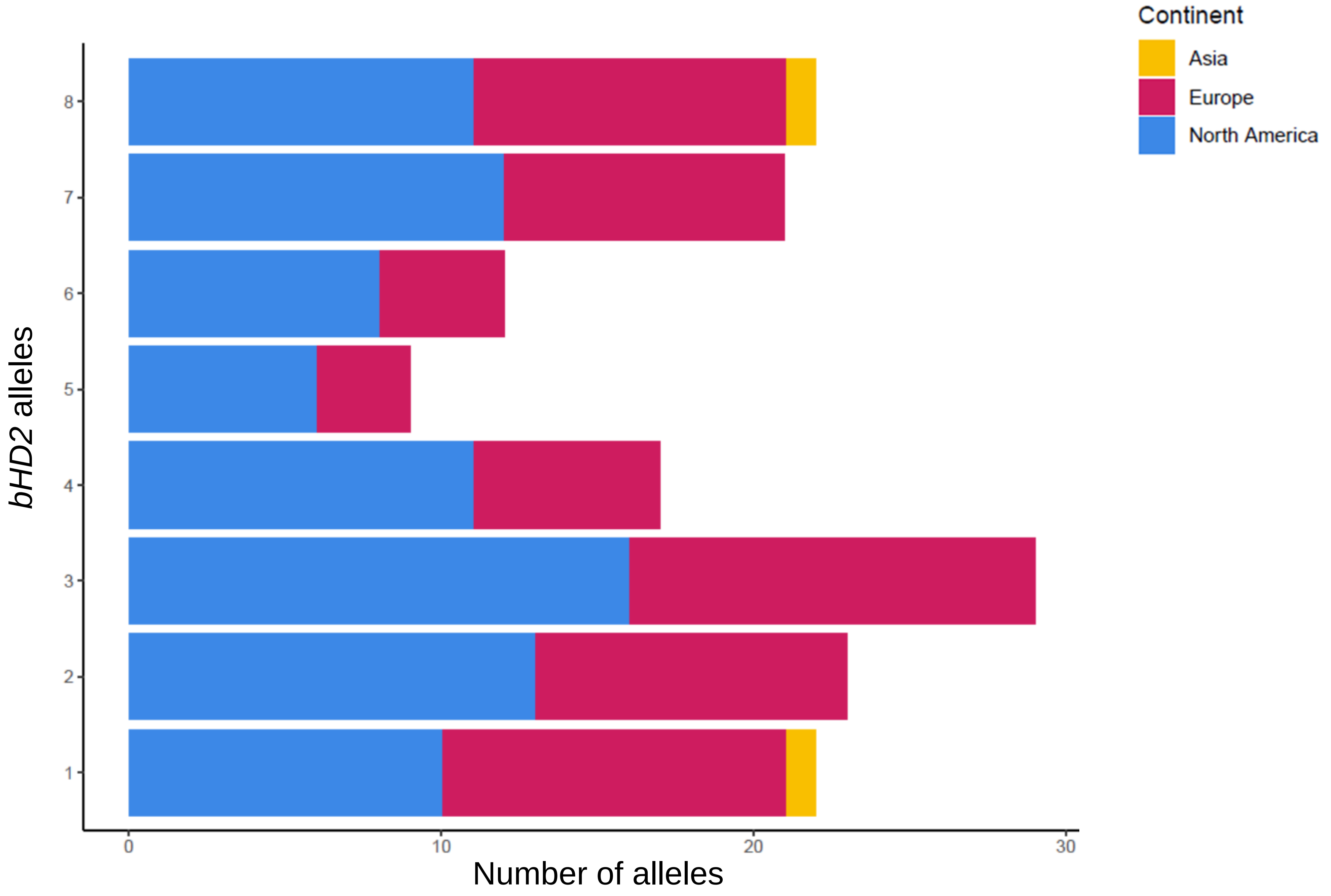

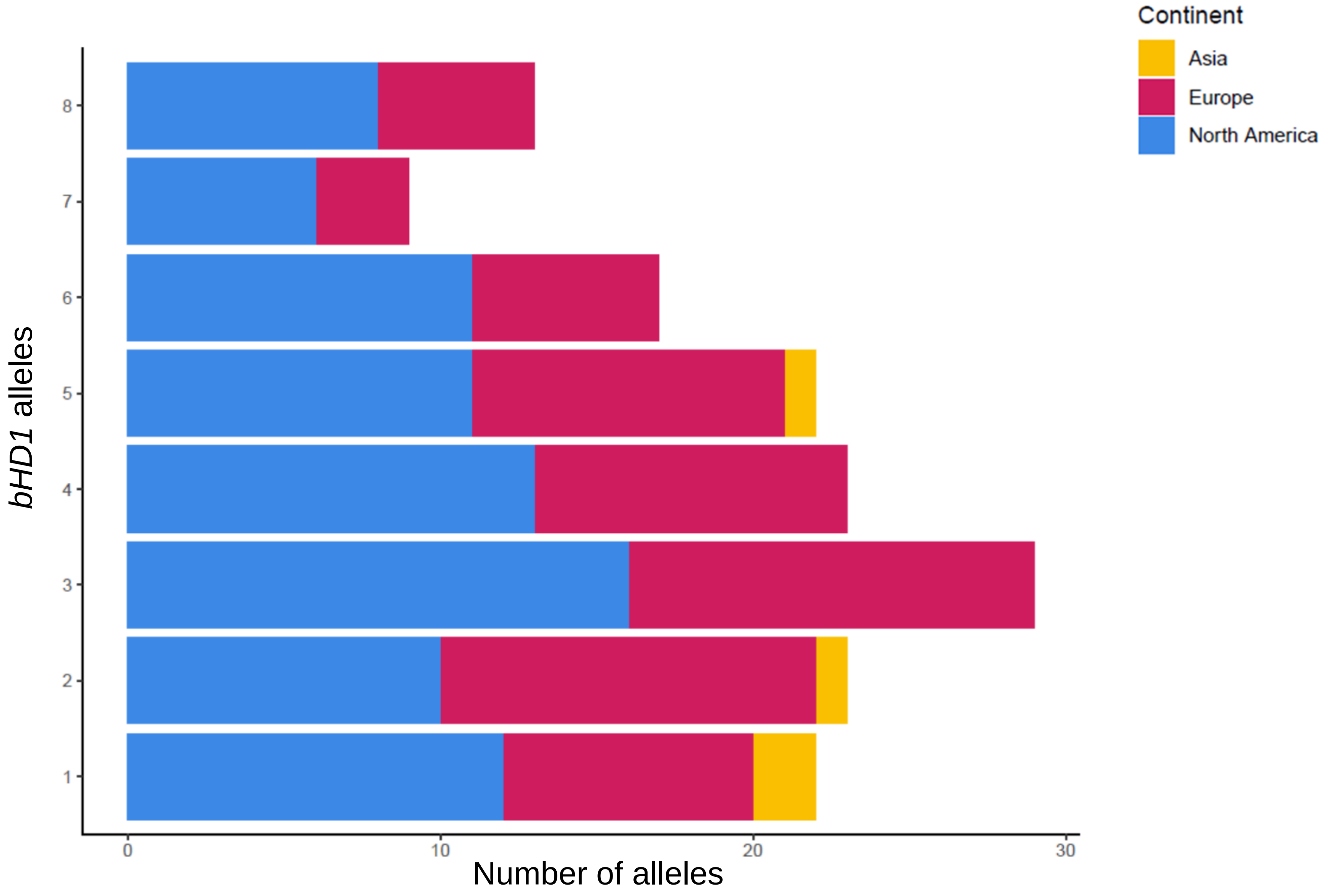

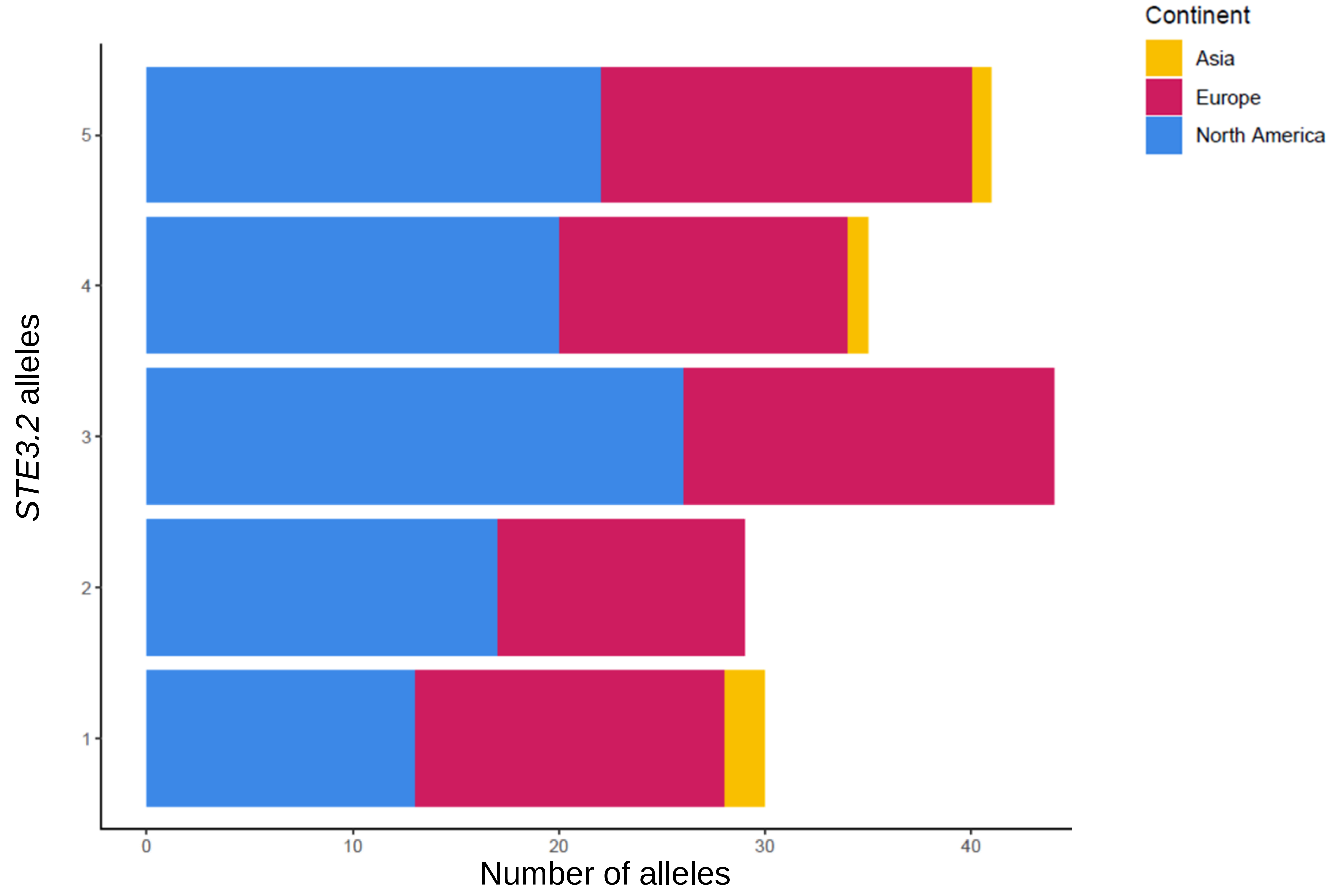

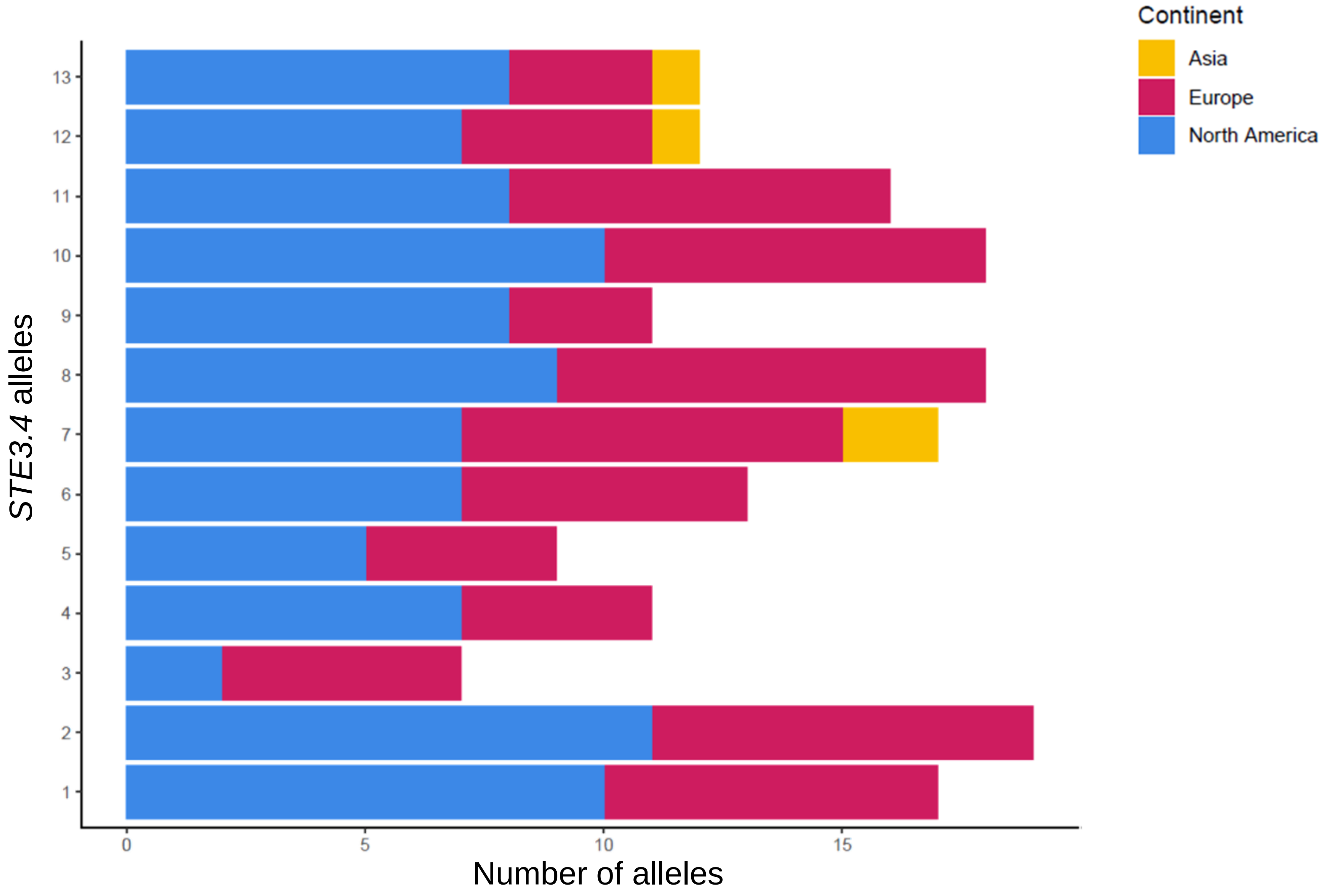

### Supplementary Figure 10.pdf

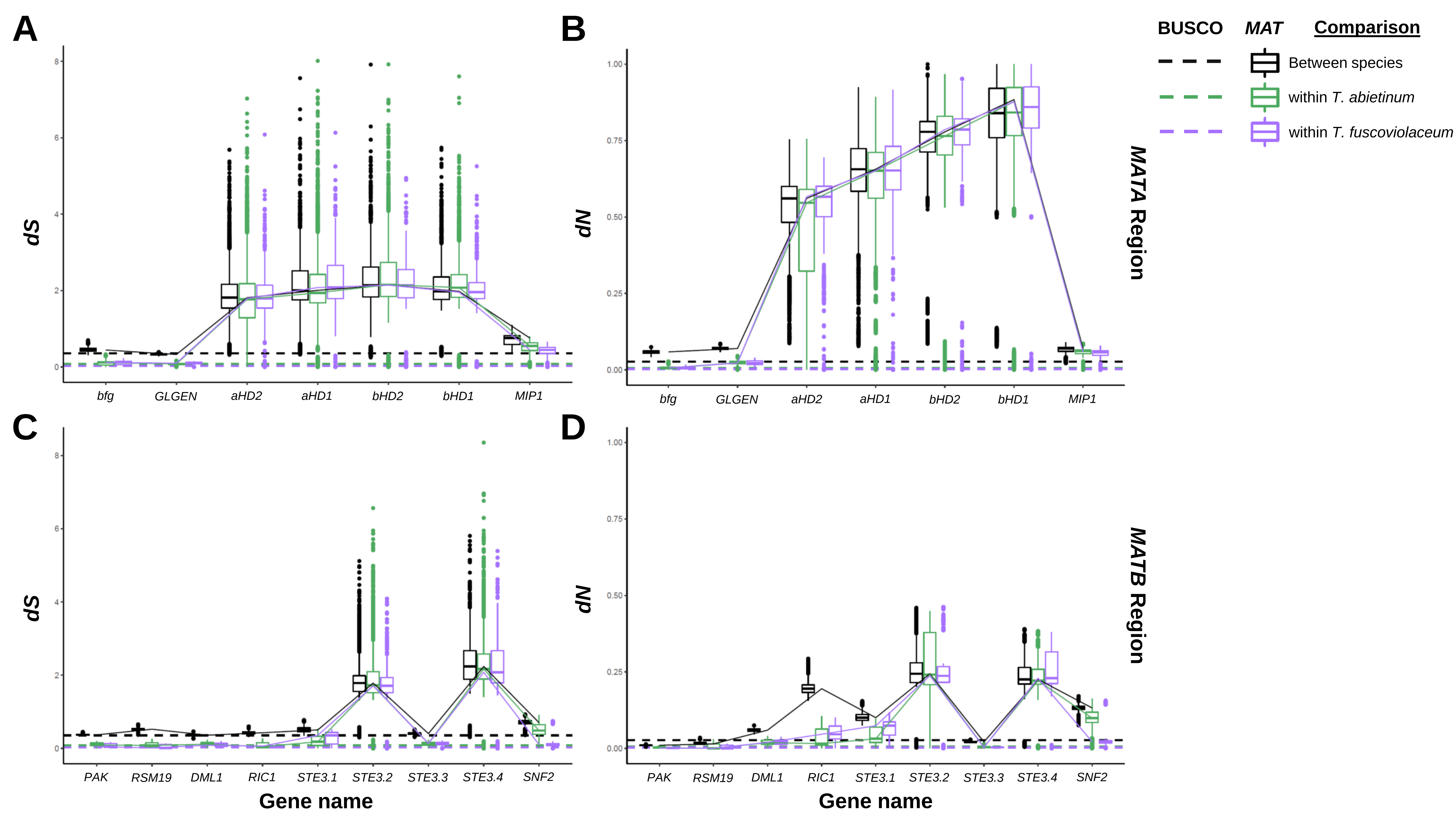

### Supplementary Figure 11.pdf

27296at155619

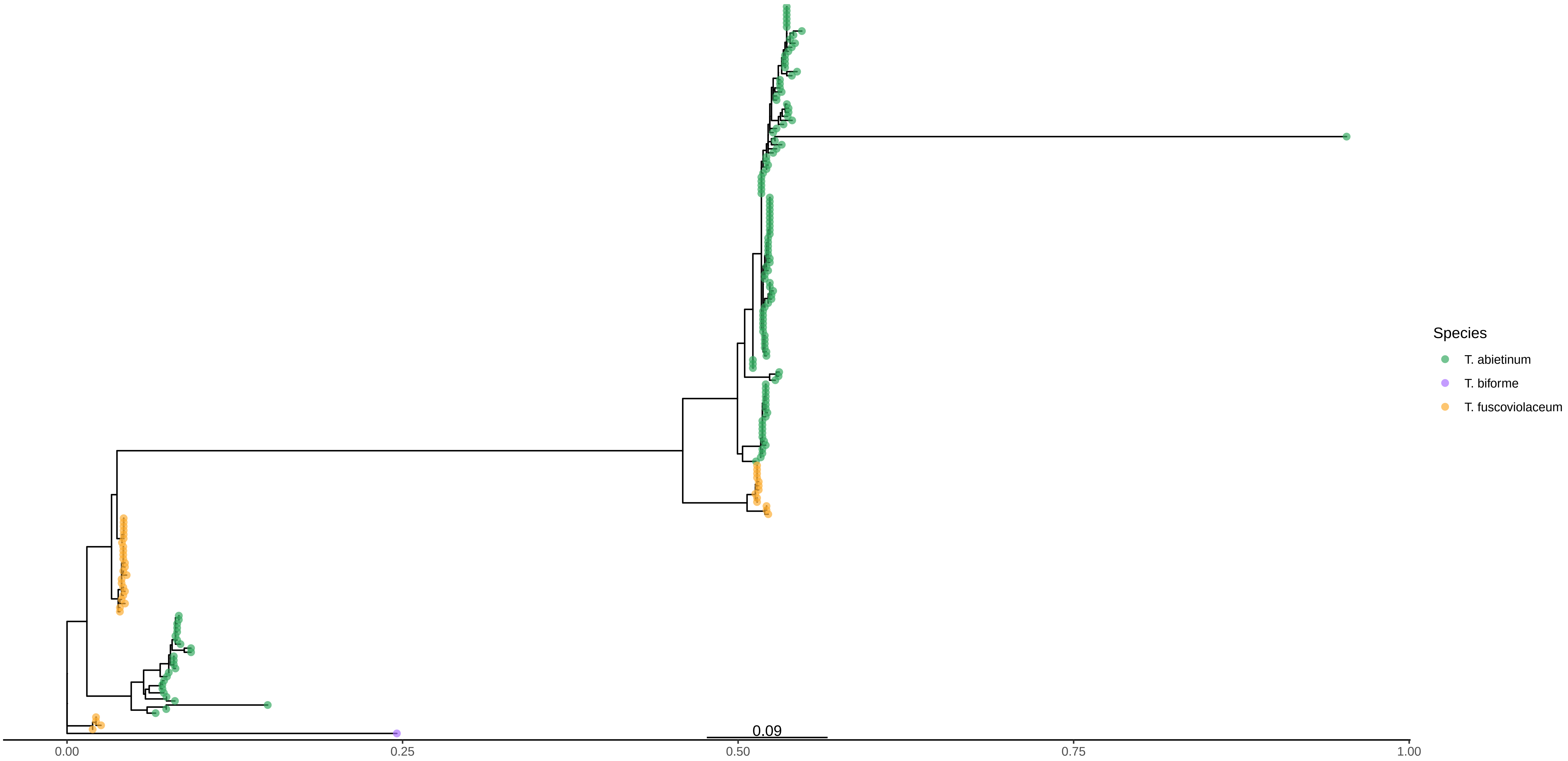

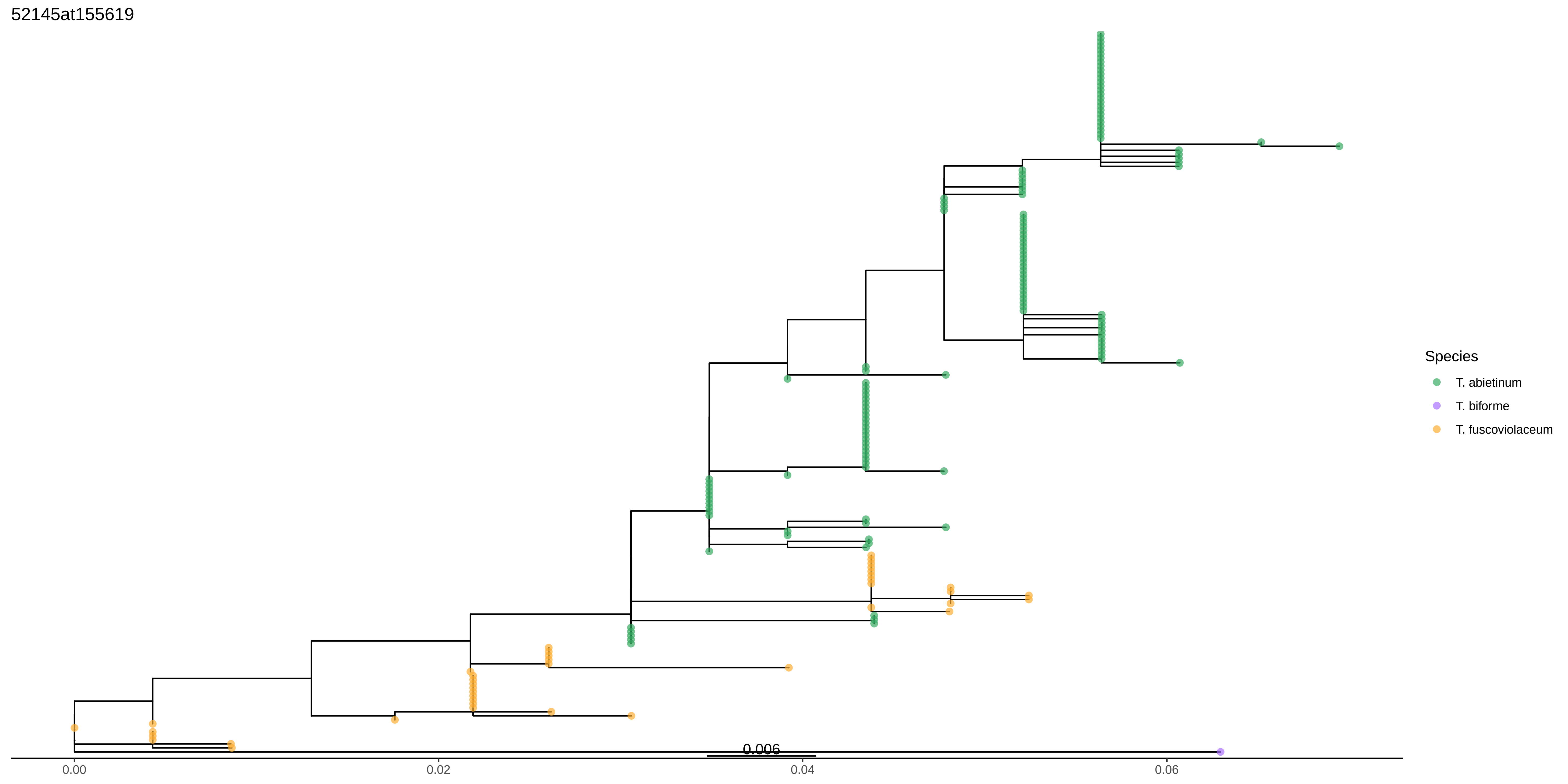

6755at155619

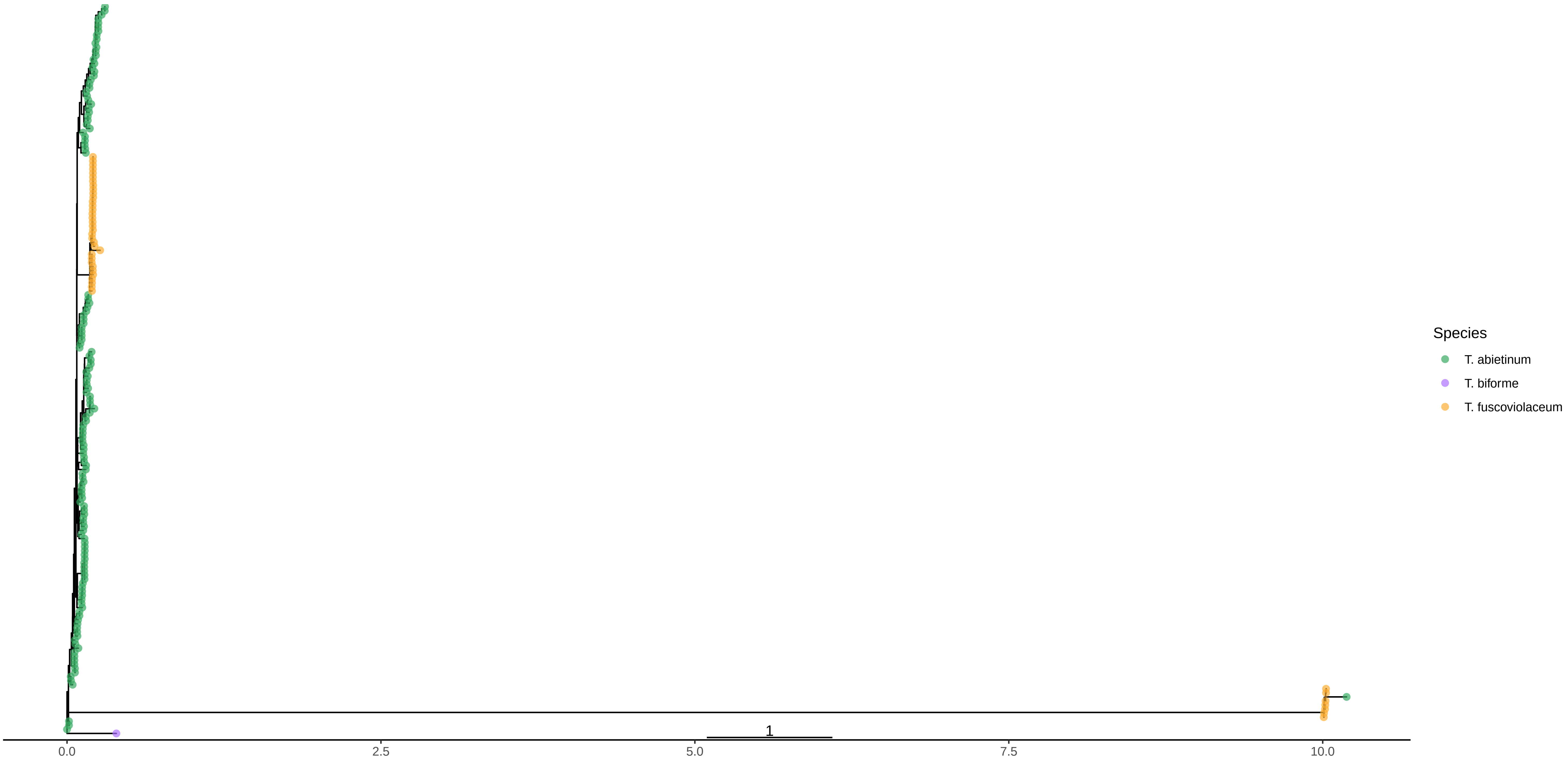

18163at155619

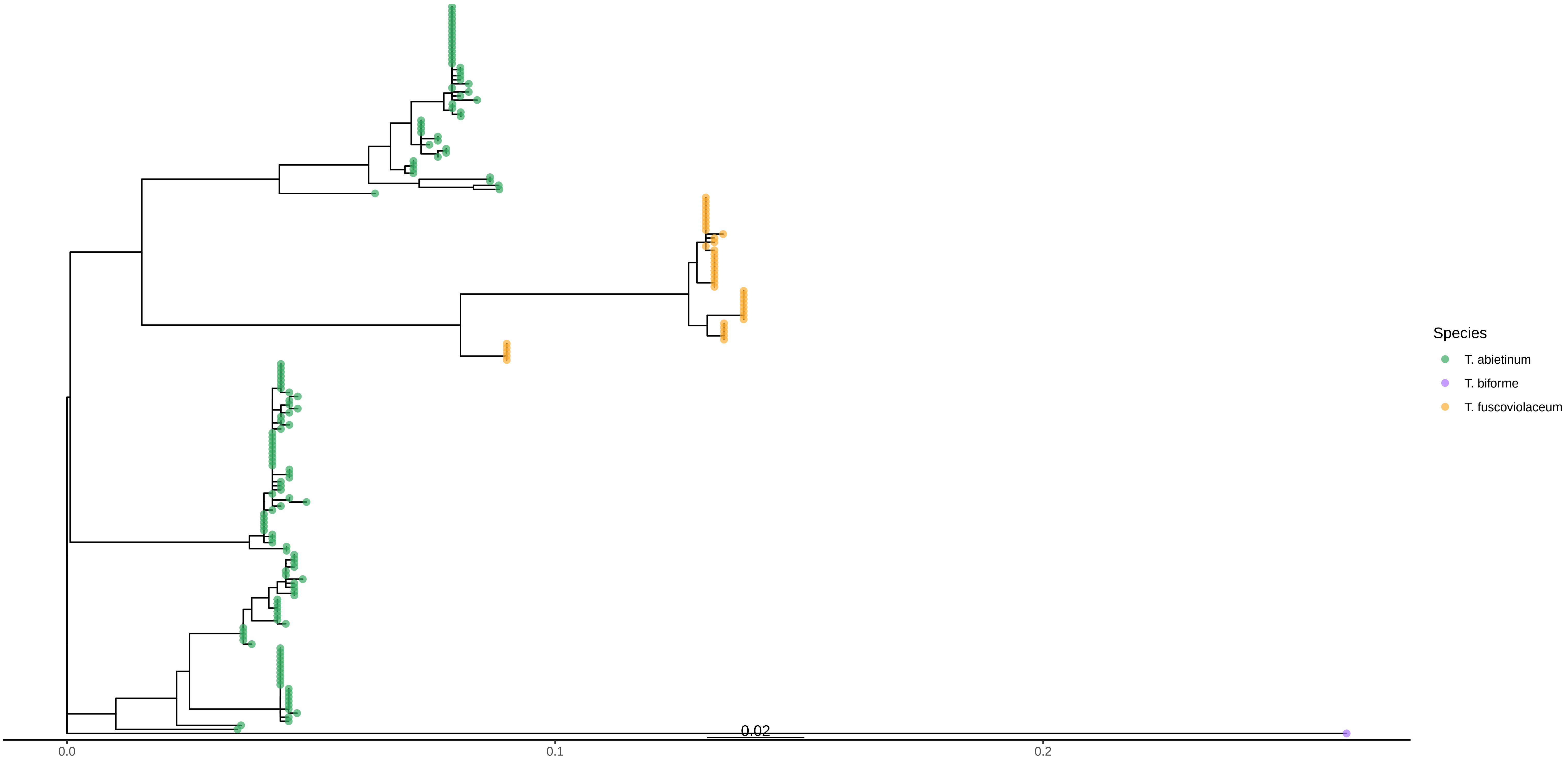

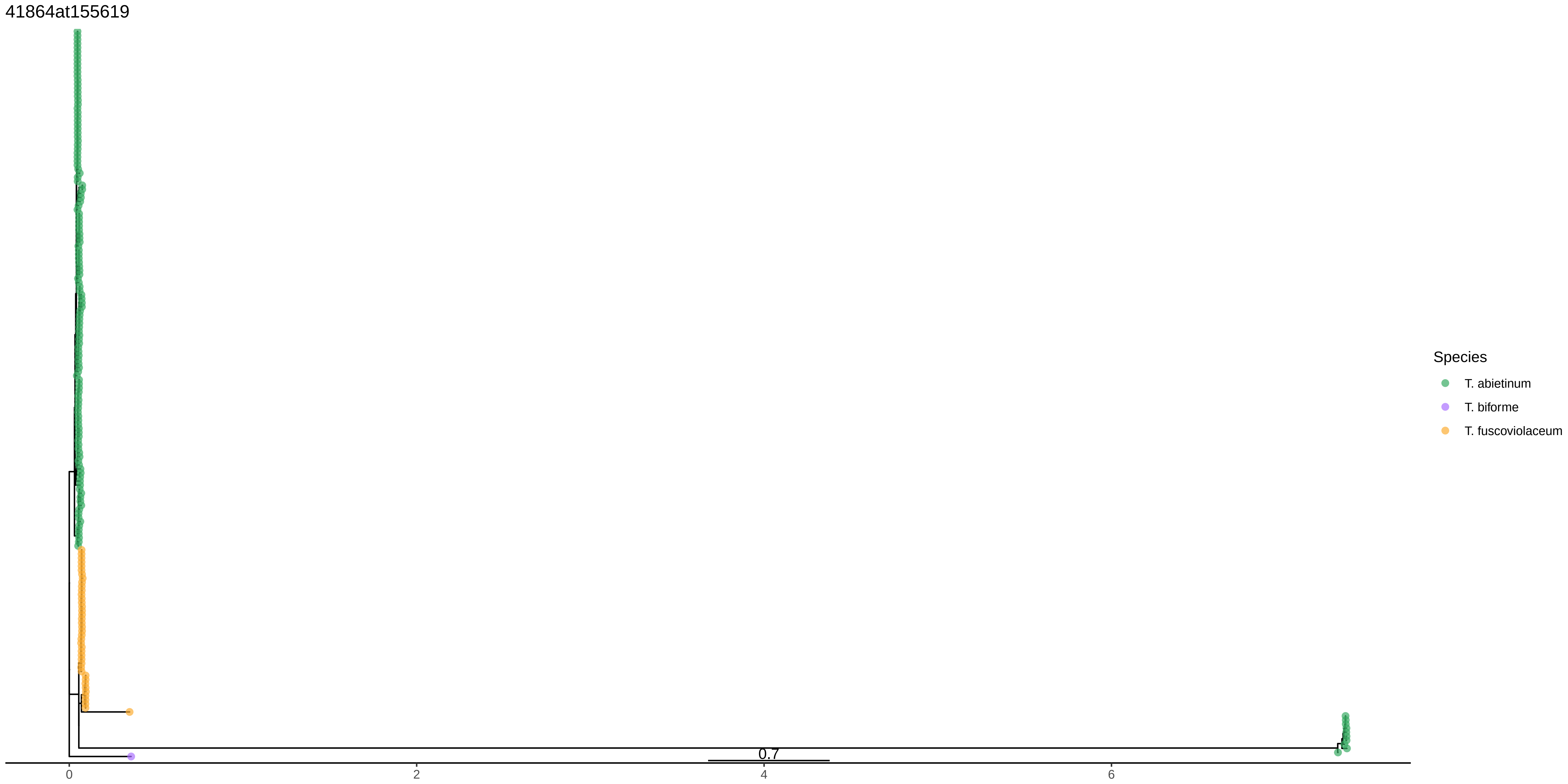

### Supplementary Figure 12.pdf

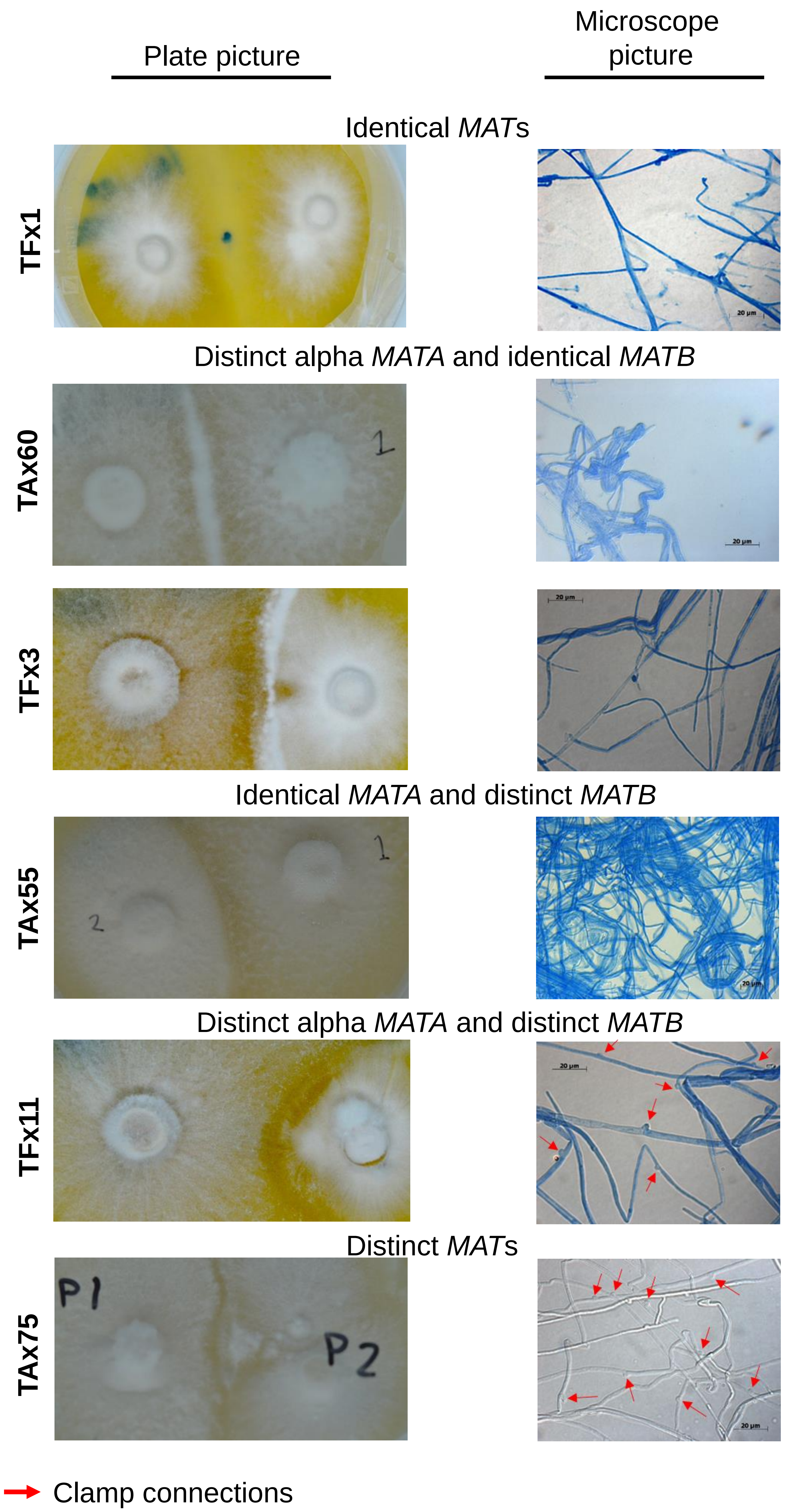
